# Population sparseness determines strength of Hebbian plasticity for maximal memory lifetime in associative networks

**DOI:** 10.1101/2025.06.16.659837

**Authors:** Naomi Auer, Lars Chen, Jakob Stubenrauch, Benjamin Lindner, Richard Kempter

## Abstract

The brain can efficiently learn and form memories based on limited exposure to stimuli. One key factor believed to support this ability is sparse coding, which can reduce overlap between representations and minimize interference. It is well known that increased sparseness can enhance memory capacity, yet its impact on the speed of learning remains poorly understood. Here we analyze the relationship between population sparseness and learning speed — specifically, how the learning speed that maximizes memory capacity depends on the sparseness of the neural code, and how this in turn affects the network’s maximal capacity. To this end, we study a feedforward network with Hebbian and homeostatic plasticity and a two-state synapse model. The network learns to associate binary input-output pattern pairs, where sparseness corresponds to a small fraction of active neurons per pattern. The learning speed is modeled as the probability of synaptic changes during learning. The results presented in this manuscript are based on both network simulations and an analytical theory that predicts expected memory capacity and optimal learning speed. For both perfect and noisy retrieval cues, we find that the optimal learning speed indeed increases with increasing pattern sparseness — an effect that is more pronounced for input sparseness than for output sparseness. Interestingly, the optimal learning speed stays the same across different network sizes if the number of active units in an input pattern is kept constant. While the capacity obtained at optimal learning speed increases monotonically with output sparseness, its dependence on input sparseness is non-monotonic. Overall, we provide the first detailed investigation of the interactions between population sparseness, learning speed, and storage capacity. Our findings propose that differences in population sparseness across brain regions may underlie observed differences in how quickly those regions adapt and learn.

**Author summary:** The brain can efficiently learn and form memories based on limited exposure to stimuli. One key factor believed to support this ability is the way neural circuits encode and organize information about the external world. Population sparseness refers to the phenomenon in which, in many brain regions, only a small subset of neurons is active at any given time or in response to a particular stimulus. Sparse codes are believed to reduce overlap between representations and minimize interference and can thus enhance the storage capacity of a network. Here we investigate the effect of population sparseness on the speed of learning input-output associations in a network model with Hebbian learning. We find that the learning speed that yields the maximal capacity increases with increasing sparseness. The maximal capacity increases for sparser input and output representations but in the first case only up to a certain point before sparseness becomes destructive. These findings could contribute to explaining why some brain regions learn faster than others.

## Introduction

Across many regions of the brain, information seems to be represented by the activity of small fractions of the total number of neurons instead of single neurons or whole populations of neurons. In particular, the dentate gyrus (DG) is known to be dependent on very population sparse activity; estimates of population sparseness based on the expression of immediate early genes in the DG of rodents propose a percentage of active cells in the one-digit range [1–3]. Other hippocampal subfields, such as CA1 and CA3, also use sparse coding, but their activity patterns are believed to be less sparse than in DG [2, 4–6]. Outside the hippocampus, population sparseness has been investigated most extensively in the visual system, because it can explain various visual response properties [7, 8]. Estimates for the sparseness level cover a wide range of values, which may be due to different recording techniques, quantification measures, and behavioral paradigms. However, there is a consensus that only a fraction of the neurons available in principle in a particular visual area of the cortex is used to represent a visual stimulus [9–14]. Further, population sparseness has been quantified in the auditory system [15–17], the olfactory system [18–21], the somatosensory system [22], and the amygdala [23]. In general, representations appear to become sparser along the hierarchy of sensory processing [8, 16, 24–27].

In addition to the comprehensive experimental evidence, there are several reasons why sparse coding is favorable from a theoretical point of view. Famously, due to the reduced overlap between patterns, sparse representations can increase the memory capacity of associative neural networks. The dependence of the capacity on the fraction of active neurons was first shown by [28]. Later, [29] found that sparse patterns yield a higher capacity than dense patterns also in the Hopfield net [30, 31]. In 1987, [32] calculated the theoretical maximum capacity, and soon after [33] proposed a weight configuration that can achieve a capacity close to the theoretical optimum. The beneficial effect of sparseness on the storage capacity of different network structures has since been studied extensively and also confirmed for palimpsest-like networks [34–41].

Sparser representations are also favorable because they seem to enable faster learning, which is due to the reduced overlap between representations that reduces interference between patterns. In contrast to the well-established increased storage capacity, higher plasticity or faster learning as benefits of sparser coding are hardly discussed in the literature. For example, the hippocampus, which is well known for its highly sparse representations, is also known to be involved in rapid learning with high plasticity of synapses, serving episodic and spatial memory. A high turnover of dendritic spines as an indicator for high plasticity has been reported for the mouse hippocampus [42]. Studies using single-trial learning paradigms showed that the hippocampus can encode and store information after only a single exposure to an experience [43, 44]. This is often called one-shot learning. Research with rodents [45–47] and humans [48–51] demonstrates that concept cells can develop very quickly (within seconds to minutes), and that hippocampal neurons can rapidly form and reorganize place fields or episodic representations during single learning episodes.

Compared to the hippocampus, the neocortex seems to have lower population sparseness. According to the theory of complementary learning systems in the context of systems memory consolidation, the neocortex is assumed to be a slow learning system, which needs repetitive exposure to acquire a new memory, for example, in the form of repeated memory replay by the hippocampus during slow-wave sleep [52–54]. This is in line with the observation that plasticity in the neocortex is lower than in the hippocampus [55, 56]. An artificial increase of plasticity in the neocortex has been reported to disrupt memory processes [57]. However, engrams can also form rapidly in the neocortex, either in parallel to engrams in the hippocampus or even without hippocampal support [43, 58, 59]. Fast memory formation in the neocortex was observed, for example, for memories related to prior knowledge in rodents [60, 61] and during the learning of new words in humans [62]. A recent analysis of a large fMRI dataset suggests that under specific conditions the human parahippocampal cortex (PHC), a part of the neocortex, can exhibit even higher plasticity than the hippocampus [63]. Even though various areas of the brain, in particular the hippocampus, are known for exhibiting sparse representations and fast learning, the association between these two features has not been properly analyzed so far.

In [64], it was briefly outlined that a smaller fraction of active cells allows for a higher learning rate and hence faster convergence in a gradient descent learning algorithm, but how this relates to the capacity of the network was not investigated. Furthermore, it is not immediately clear how this translates to biologically more plausible learning paradigms. [15] generated neuronal activity patterns based on sparse or dense firing rate distributions. A sequence of patterns alternating between a fixed target pattern and changing non-target patterns was used to train a single neuron with Hebbian synapses. The ratio of the target pattern response and the non-target pattern responses converges faster and to a larger value if the patterns were sampled from the sparse distribution. To the best of our knowledge, there is no deeper investigation of the dependence of the speed of learning on population sparseness.

In the present paper, we investigate the dependence of the optimal strength of plasticity (interpreted as the learning speed of the network) on the sparseness of representations in a scenario with Hebbian and homeostatic plasticity; furthermore, we explain the impact of learning speed and sparseness on the memory lifetime. To simplify the approach as much as possible, we use a heteroassociative feedforward network of neurons with binary activations, connected by two-state synapses; the network learns pairs of input-output patterns in a supervised fashion. Based on the underlying assumption that the memory capacity of the network should be maximal, we examine the role of fractions of active neurons and probabilities of synaptic changes in numerical simulations and analytical derivations.

This manuscript is organized as follows: We first introduce the model and the learning rule in more detail and explain how the capacity of the network is quantified. Then, we explain the relationship between the optimal learning speed and the sparseness of both the input and the output patterns. We also show how choosing the optimal learning speed for each sparseness value impacts the maximal memory capacity. The next section provides a more intuitive understanding of these results based on analytical considerations. Further, we discuss the effect of a change of the network size and analyze whether the fraction of active neurons or the absolute number of active neurons is the critical parameter. Finally, we add noise to the input pattern during retrieval in order to generalize our results to the biologically more realistic case of non-perfect retrieval cues.

## Results

### Network model, learning paradigm, and quantification of capacity

We are interested in the effect of population sparseness on the optimal learning speed of an associative network, and we hypothesize that sparse representations facilitate a large capacity while learning fast. This hypothesis is based on the following considerations: In order to establish an association between two neuronal assemblies, the synapses between the neurons that are part of one assembly and the neurons of the second assembly need to be strengthened. If *f* is the fraction of the total number of neurons that forms an assembly (activation ratio), the number of potential connections between neurons of the two assemblies scales with *f* ^2^. A low activation ratio is interpreted as high population sparseness, while a large activation ratio means dense representations and hence a low population sparseness (Fig 1A). Learning an association between two sparser assemblies involves only a smaller fraction of synapses. In this case, the memory capacity can be larger because each association affects fewer synapses. On the other hand, an association between denser assemblies involves the modification of a larger fraction of synapses, which can lead to catastrophic interference and lower capacity. For example, for sparser assemblies where only 1% of the cells is active (*f* = 0.01), only a fraction *f* ^2^ = 1*/*10^4^ of the synapses is changed, while for denser assemblies with *f* = 0.5, a quarter (*f* ^2^ = 0.25) of all synapses potentially has to be updated.

**Fig 1.**
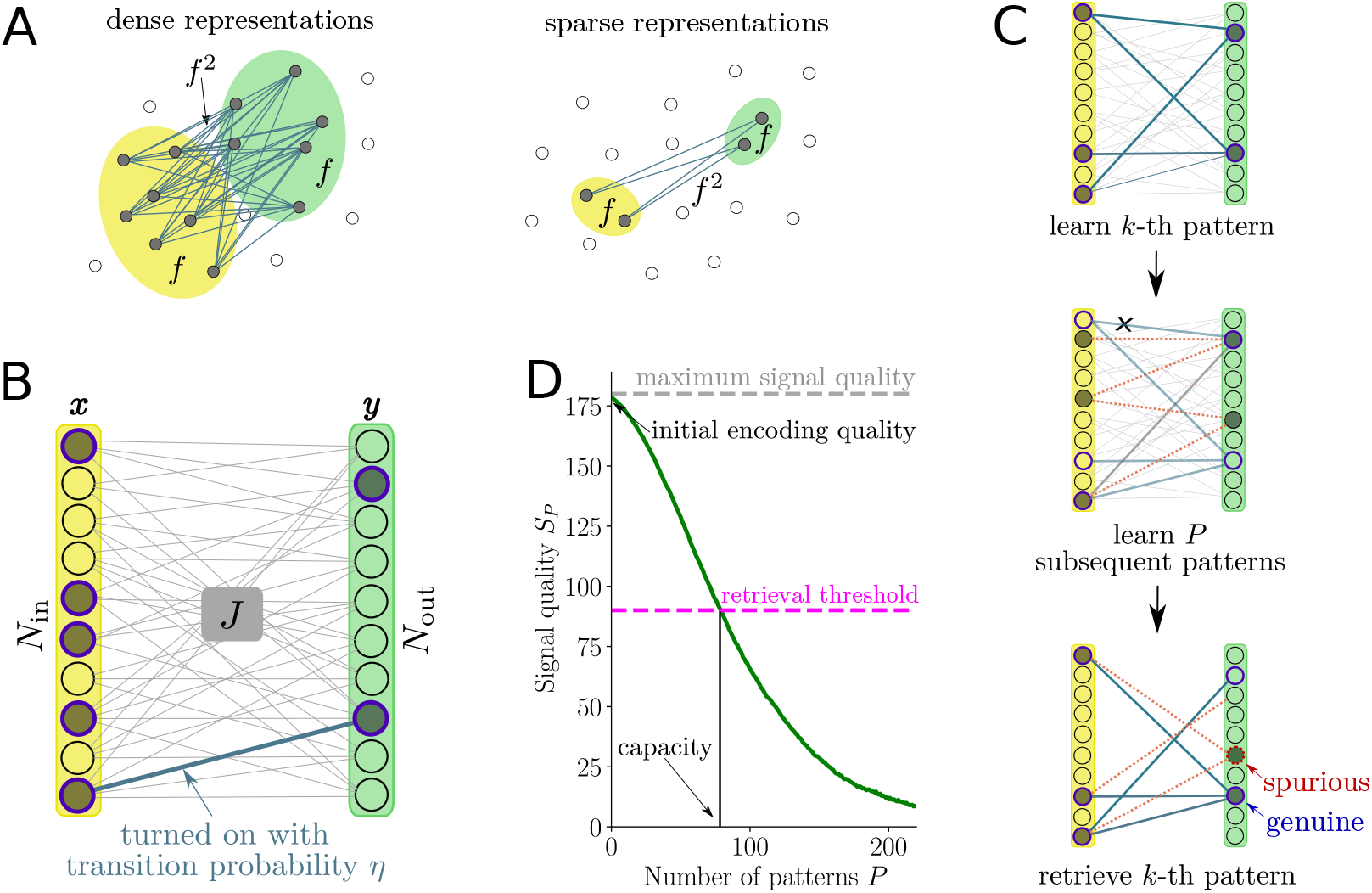
Sparse vs dense representations and quantification of capacity. (**A**) In a network of neurons with an activation ratio *f*, which is the number of neurons in an assembly divided by the total number of neurons in the network, the number of connections potentially involved in an association between two assemblies scales with *f* ^2^. The number of relevant connections is thus smaller for sparse representations (left) than for dense representations (right). (**B**) Illustration of the network model with an input layer (yellow) of size *N*_in_ (black circles) with input patterns **x** (indicated by gray discs and blue circles) of activation ratio *f*_in_ and an output layer (green) of size *N*_out_ with output patterns **y** (gray discs and blue circles) of activation ratio *f*_out_ that are connected by a binary weight matrix *J* with a morphological connectivity *c*_*m*_ (not shown) and a functional connectivity *c* (lines between input and output layer) (both normalized per output unit). During learning, the connections between active units are turned on with transition probability *η*; a homeostasis mechanism maintains the functional connectivity *c*. (**C**) Schematic illustration of the learning and retrieval processes. After learning the *k*-th pattern, *P* additional patterns are learned. In order to determine the memory signal quality of the *k*-th pattern at time *P*, input pattern **x**^[*k*]^ is presented to the network again to calculate an output pattern **ŷ**^[*k*]^(*P*), which can then be compared to the target output pattern **y**^[*k*]^. (**D**) The quality of the memory signal (signal quality) is defined as *H*_avg_ − *H*(**y**^[*k*]^, **ŷ**^[*k*]^(*P*)) (green line). It decays with the number of subsequently stored patterns *P*. The average Hamming distance *H*_avg_ between two random *f*_out_-sparse vectors of length *N*_out_ (dashed grey line) is an upper bound of the average signal quality. The retrieval threshold is marked as *T*_*S*_ (dashed magenta line). The capacity of the network is defined as the number of patterns *P* for which the signal quality reaches the retrieval threshold *T*_*S*_.

We investigate this hypothesis in a hetero-associative one-layer feed-forward network with *N*_in_ input and *N*_out_ output units (1B), similar to the architecture of a Willshaw network [28]. A memory is a one-way association between an input pattern **x** and an output pattern **y**. Units have two states and can be either active (1) or inactive (0). The input and output activation ratios *f*_in_ and *f*_out_ are defined as the fraction of active units in the input representation and the output representation, respectively. We denote the number of active input units in an input pattern **x** as *M*_in_ = *f*_in_*N*_in_; similarly, the number of active output units in a target output pattern **y** is *M*_out_ = *f*_out_*N*_out_. We typically assume *M*_in_ ≫1 and *M*_out_ ≫ 1. Input and output layers of the network are connected by a weight matrix *J* with binary entries. For the construction of *J*, we distinguish between morphological and functional connectivity. In biological neural systems, the number of morphological (or structural) synapses per unit is constrained. Therefore, we assume a morphological connectivity 0 *< c*_*m*_ ≤ 1. The morphological connectivity is normalized per output unit, thus each output unit is targeted by exactly *c*_*m*_*N*_in_ morphological connections chosen randomly among all possible connections from input units. Entries of the weight matrix *J* that do not correspond to a morphological connection are permanently set to zero. We have a total number of *c*_*m*_*N*_in_*N*_out_ morphological connections between the input and the output units, and each connection can be functional (on, value 1) or silent (off, value 0). The weight matrix is initialized such that each output unit is targeted by exactly *cN*_in_ connections that are functional (thin grey lines in Fig 1C top), and we call *c* the functional connectivity. Note that the functional connectivity is bounded by the morphological connectivity: 0 *< c* ≤ *c*_*m*_.

The network learns a sequence of uncorrelated input and output pattern pairs (**x**^[*k*]^, **y**^[*k*]^)_*k∈*{0,1,…}_, which are pairs of randomly drawn binary vectors that fulfill the sparseness constraint given by *f*_in_ or *f*_out_. Since there is no explicit time component, presenting one pattern pair to the network can be interpreted as one step in time. For each pattern, the weights of the connection matrix *J* are updated in the following way: First, we consider a connection between an active input unit (filled grey circle in yellow layer in Fig 1C top) and an active output unit (filled grey circle in green layer in Fig 1C top). Such a connection can only change if there is a morphological connection. A connection that was already functional stays on, i.e., it is not changed (thin blue line originating from last input unit in Fig 1C top). A connection that was silent is turned on with transition probability *η*. These connections are depicted as thick blue lines in Fig 1C top. We interpret the transition probability 0 ≤ *η* ≤ 1 as the learning speed (or learning strength) of the network. For a high transition probability *η*, many connections are updated at once and the memory is strongly engraved into the network.

Furthermore, a homeostasis mechanism maintains the functional connectivity *c*. Previously functional connections are turned off in order to balance the additional connections turned on during a learning step. Otherwise all morphological connections would eventually become functional. In principle, there are many possible ways to normalize the number of functional connections and provide a homeostatic stabilization. Inspired by studies suggesting that neurons monitor the total excitation that they receive and the summed synaptic surface area per length of dendritic segment is conserved over time [65, 66], we apply a post-synaptic homeostasis step after each learning step. The functional connectivity *c* is maintained per output unit by randomly silencing the necessary number of connections from inactive input units to the active output units. To this end, we must assume *f*_in_ ≤ *c* (see Methodsfor details). Connections targeting inactive output units are not changed. This combination of Hebbian potentiation and homeostatic depression realizes a learning rule that is similar to the ones suggested by [35] and by [36].

To asses memory capacity, we quantify changes of the memory trace of the *k*-th pattern pair after learning *P* subsequent pattern pairs. With increasing *P*, the memory trace deteriorates more until the memory is not retrievable anymore according to some criterion. We always understand the time point *P* as relative to the *k*-th pattern. So after learning the *k*-th pattern, *P* additional patterns (illustrated for *P* = 1 in Fig 1C center; units active in the *k* + 1st pattern are shown as filled grey circles, units active in the *k*-th pattern are shown as circles with dark blue stroke) are learned by turning on connections that are relevant for these patterns and turning others off, potentially also connections that were relevant for the *k*-th pattern. Connections that are turned on in this step are shown as dashed orange lines and connections that are silenced are shown as light blue lines that are crossed out. The network hence realizes a palimpsest memory system [67] where older memories are erased by newer memories due to the homeostasis mechanism.

In order to quantify memory capacity, we distinguish two types of output units: Output units are either *genuine* with respect to the *k*-th pattern, hence active in the original target output **y**^[*k*]^ or *spurious* with respect to the *k*-th pattern, hence inactive in the target output **y**^[*k*]^. During retrieval, genuine output units might become inactive and spurious units might become active. At any time step *P* after learning the *k*-th pattern, the retrieval of the *k*-th input-output pattern pair can be tested by presenting the original input pattern and calculating the output with the current weight matrix *J* ^[*k*+*P*]^ (Fig 1C bottom). Some of the relevant connections of the *k*-th pattern are still functional (blue lines) and provide input to the genuine output units (blue output units in Fig 1C, bottom). Other connections from the same input units might be functional due to learning subsequent patterns (dashed orange lines) or due to the initial weights before learning the *k*-th pattern. These connections provide input to spurious output units (red output unit in Fig 1C bottom).

Output units are modeled as McCulloch-Pitts neurons [68], which sum the weighted inputs and compare them against an activation threshold 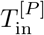. Their output is calculated by

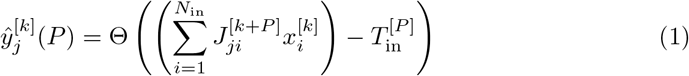

where

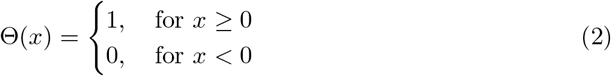

is the Heaviside step function. The activation threshold 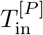 is adjusted in every time step such that the calculated output pattern has the same activation ratio *f*_out_ as the target output. This phenomenologically describes latent competition where *M*_out_ = *f*_out_*N*_out_ winners take all and the rest is deactivated. Note that this choice of threshold is motivated by the comparability of results based on fixed sparseness values. It is not necessarily an optimal activation threshold in that it maximizes capacity or other performance metrics. The difference between the output **ŷ**^[*k*]^(*P*) obtained with the weight matrix *J* ^[*k*+*P*]^ at time (or pattern) *P* and the original target output **y**^[*k*]^ of this input pattern is measured by the absolute number of bit flips between **ŷ**^[*k*]^(*P*) and **y**^[*k*]^, termed the Hamming distance

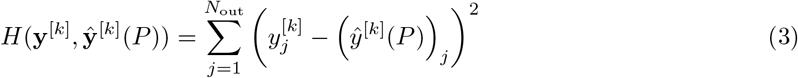

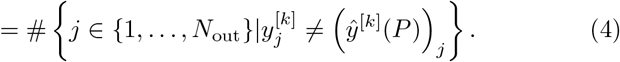

The larger the number of subsequent patterns *P* that have been learned since the one that is being tested for retrieval, the more the obtained output will differ from the target output because the summed inputs to the genuine units become too small compared to the summed inputs of the spurious units.

To quantify the strength of the memory after *P* time steps, we define the signal quality

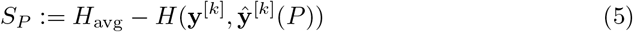

where the first term *H*_avg_ := 2*N*_out_*f*_out_(1 − *f*_out_) is the average Hamming distance between two random *f*_out_-sparse patterns (Fig 1D, dashed grey line) and the second term is the Hamming distance between the calculated output representation **ŷ**^[*k*]^(*P*)) and the target **y**^[*k*]^ (Fig 1D). For two random *f*_out_-sparse patterns, an error occurs if either the bit of the first pattern is 1 (with probability *f*_out_) and the bit of the second pattern is 0 (with probability 1 − *f*_out_) or the first bit is 0 (with probability 1 − *f*_out_) and the second one is 1 (with probability *f*_out_); the probability for a deviation between the two entries is thus the sum over the probabilities of the two distinct events. Summed over all entries, this gives *H*_avg_.

We define the memory capacity *P*\* of the network as the lifetime of a memory stored in the network, which is the time that passes until the signal of the memory has decayed to the point that the memory cannot be successfully retrieved anymore. Alternatively, it is the number of patterns that can be stored in addition to the first pattern, without destroying the retrieval of the first pattern. The capacity is determined by introducing a retrieval threshold *T*_*S*_ (Fig 1D, dashed magenta line), which represents the error tolerance between the obtained output and the target. The number of subsequently stored patterns for which the signal quality reaches the retrieval threshold can be interpreted as the lifetime of the memory or the memory capacity *P*\* of the network, defined by the equation

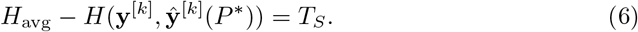

We define the threshold

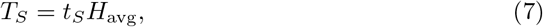

with *t*_*S*_ ∈ (0, 1), as a fraction of the maximal signal quality which is the Hamming distance between two random *f*_out_-sparse vectors *H*_avg_. A fixed number of wrongly activated units thus has a more severe destructive effect if the total number of active output units *M*_out_ = *N*_out_*f*_out_ is small. Whenever not further specified, we choose *t*_*S*_ = 0.5 and thus *T*_*S*_ = *N*_out_*f*_out_(1 − *f*_out_).

### Theory on distributions of dendritic sums and calculation of capacity

The memory capacity of the network can be calculated numerically in simulations but it can also be described analytically in a probabilistic sense. Given a particular input pattern **x**^[*k*]^, the summed input that an output unit receives is called the dendritic sum of this output unit. Output units can potentially be either genuinely or spuriously active in the calculated output **ŷ**^[*k*]^(*P*) depending on whether they are active or inactive in the target output **y**^[*k*]^. For each output unit, the dendritic sum hence follows either the distribution of dendritic sums of spurious output units (red distribution in Fig 2A, will be called *spurious distribution*) or the distribution of dendritic sums of genuine output units (blue distributions in Fig 2A, will be called *genuine distribution*).

**Fig 2.**
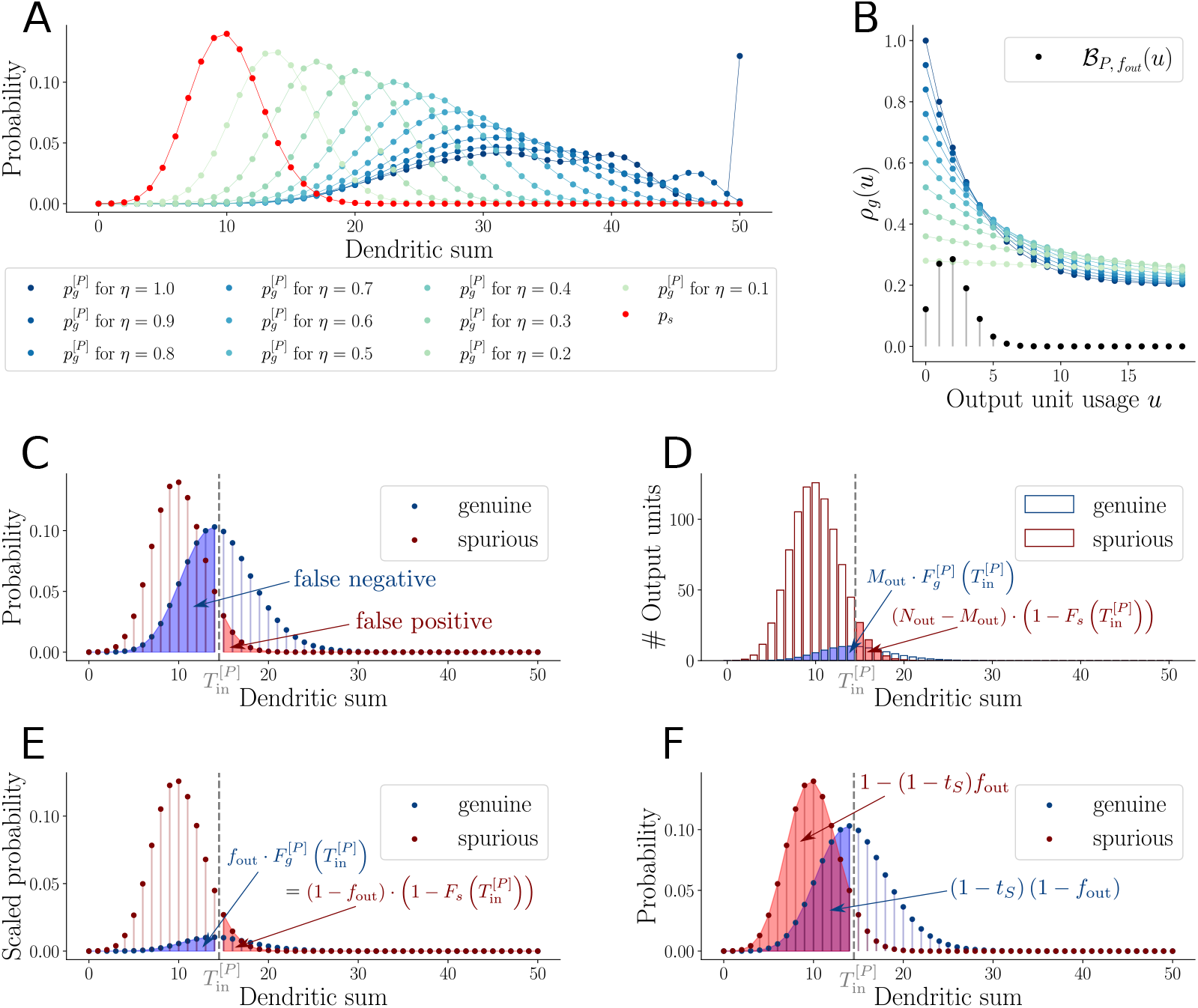
Distributions of dendritic sums and theory on capacity. (**A**) The smaller the transition probability *η*, the closer the PMF of the dendritic sums of genuine output units (blue) is to the PMF of the dendritic sums of spurious output units (red). (**B**) The probability of a functional connection to a genuine output unit decays with the output unit usage *u*. The probability 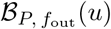 that an output unit is active *u* times across *P* patterns is the weight of the corresponding binomial PMF in Eq. (9). Different shades of blue represent different transition probabilities *η* (same as in (**A**)). In (**A**) - (**B**), *P* = 20. (**C**) Spurious units with a dendritic sum larger than the activation threshold 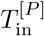 (red area) are wrongly activated (false positive); genuine units with a dendritic sum smaller than 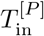 are wrongly not activated (false negative). (**D**) shows a histogram of the dendritic sums of output units. The expected Hamming distance is the sum of falsely deactivated genuine units (blue area) and falsely activated spurious units (red area). In (**E**), the PMFs of dendritic sums of genuine (blue) and spurious (red) units are scaled by *f*_out_ and 1−*f*_out_, respectively. To respect the Balance Equation, the mass of the part of the genuine distribution left of the activation threshold 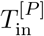 (blue area) must correspond to the mass of the part of the spurious distribution right of 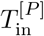 (red area). (**F**) At capacity *P* = *P*\*, the activation threshold 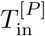 corresponds to the 1 − (1 − *t*_*S*_)*f*_out_-quantile of the spurious distribution and to the (1 − *t*_*S*_)(1 − *f*_out_)-quantile of the genuine distribution. In (**C**) - (**F**), *P* = 108. Further parameters: *N*_in_ = *N*_out_ = 1000, *f*_in_ = 0.05, *f*_out_ = 0.1, *c* = 0.2, *c*_*m*_ = 1, *t*_*S*_ = 0.5.

The probability mass function (PMF) of the dendritic sum of a spurious unit is

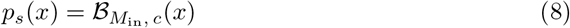

(see Fig 2A red, derived in Methods, see Eqs. (34) and (42)), where ℬ_*n,p*_ denotes a binomial distribution with *n* trials and success probability *p*. The spurious distribution is centered at *M*_in_*c* where *M*_in_ = *N*_in_*f*_in_. We note that *M*_in_*c* is constant in *P* due to the constant functional connectivity *c* that ensures a fixed number of functional connections per output unit; when a random input pattern is applied, on average, a fraction *f*_in_ of the *cN*_in_ connections receives input. The spurious distribution does not change when new uncorrelated patterns are learned, and thus the spurious distribution *p*_*s*_ does not depend on the number of patterns *P*.

In contrast, the PMF of the distribution of the dendritic sums of genuine units depends on *P*; it is given by

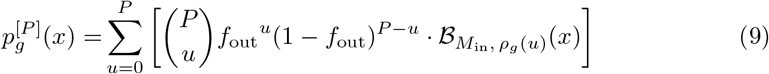

(see Fig 2A blue) with the probability for a connection to a genuine unit to be functional given that the output unit is active *u* times across the whole pattern set

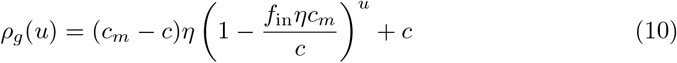

(see Fig 2B blue, derived in Methods, see Eqs. (34) and (57)). Immediately after learning a specific pattern, the probability of the relevant connections to a genuine output unit to be functional is larger than the average functional connectivity *c*, i.e., in Eq. (10) we have *ρ*_*g*_(0) = (*c*_*m*_ − *c*)*η* + *c*. Due to the homeostatic plasticity, the probability *ρ*_*g*_ is reduced every time this output unit is active in subsequent patterns, which is reflected in the term 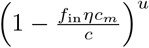 where *u* is the output unit usage. For *u* → ∞, we have *ρ*_*g*_(*u*) → *c*. The probability *ρ*_*g*_ enters Eq. (9) in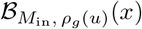, which is the distribution of the dendritic sums if the output unit was active *u* times in the other patterns. The distribution of the output unit usage *u*, hence the number of times that an output unit is active across a set of *P* → ∞ patterns, explains the weight 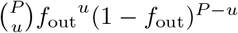 of a binomial distribution in Eq. (9) (see Fig 2B). For *P* → ∞, the genuine distribution converges to the spurious distribution. Distributions of dendritic sums obtained from numerical simulations match the distributions in Eqs. (8) and (9) very well (see example in S5 Fig).

For an analytical description of the expected Hamming distance 𝔼(*H*(**y**^[*k*]^, **ŷ**^[*k*]^(*P*))) between a target output pattern **y**^[*k*]^ and the corresponding output pattern **ŷ**^[*k*]^(*P*) calculated during retrieval, we need to calculate the expected number of output neurons that are in the wrong state in **ŷ**^[*k*]^(*P*) compared to **y**^[*k*]^. The activation threshold *T*_in_ (dashed grey line in Fig 2C) determines which output units are activated and which are not. If the distributions of dendritic sums of genuine and spurious output units are far apart and do not overlap, which can be the case during the first steps after learning a pattern, hence for small *P*, the activation threshold *T*_in_ can perfectly separate the two distributions, and then the calculated output is identical with the target output. If the two distributions are close to each other, a part of the spurious output units is wrongly activated (red area under red distribution in Fig 2C, false positive) and a part of the genuine output units is wrongly inactivated (blue area under blue distribution in Fig 2C, false negative). These two errors both contribute to the Hamming distance between the target output pattern and the calculated output pattern but they are weighted differently, as will be explained in the following (see Methods for a detailed derivation). In order to derive an actual error contribution from these distributions, we have to consider how many units have dendritic sums following either of the two distributions. As mentioned, there are *M*_out_ active units in the target output pattern. There are hence *M*_out_ output units whose dendritic sum follows the genuine distribution and *N*_out_ − *M*_out_ output units whose dendritic sum follows the spurious distribution. The false negative error and the false positive error thus have to be weighted by *M*_out_ and *N*_out_ −*M*_out_, respectively. It follows that the expected Hamming distance can be expressed as

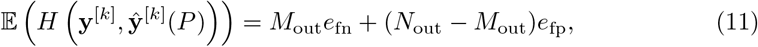

where *e*_fn_ and *e*_fp_ denote the error probabilities due to false negatives (blue area) and false positives (red area), respectively.

Let 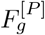 and *F*_*s*_ denote the cumulative distribution functions (CDF) of the distributions of dendritic sums of genuine units and spurious units, respectively. Then, we can express the expected Hamming distance as

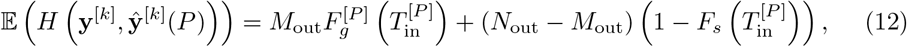

see Fig 2D (derived in Methods).

As discussed in the previous section, we always adjust the activation threshold 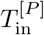 such that the output activation ratio of the target output pattern **y**^[*k*]^ is maintained in the calculated output **ŷ**^[*k*]^(*P*). This restriction is respected if the Balance Equation

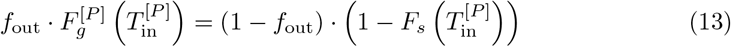

is fulfilled (see Fig 2E), i.e., if the contribution to the Hamming distance of the wrongly inactive genuine units *M*_out_*e*_fn_ equals the contribution of the wrongly active spurious units 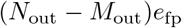 (see derivation in Methods).

With the Balance Equation, we can express 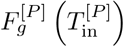 in terms of 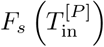 and the expected Hamming distance simplifies to

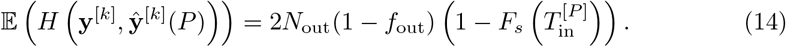

For the capacity, we need to determine the number of patterns *P* for which the signal quality *S*_*P*_ = *H*_avg_ − *H* (**y**^[*k*]^, **ŷ**^[*k*]^(*P*)) equals the retrieval threshold *T*_*S*_ = *t*_*S*_*H*_avg_ (see Methods for details), i.e., we have to solve the equation

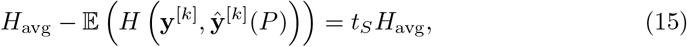

which can be simplified to

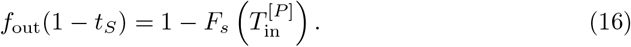

Using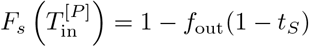 in the Balance Equation yields

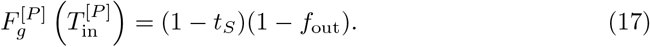

We now have two Equations (16) and (17) in two variables *P* and 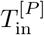. Solving them gives us the number of patterns *P* (and the corresponding activation threshold 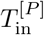) for which the signal quality *S*_*P*_ equals the retrieval threshold. This *P* is the capacity (or memory lifetime) *P*\*.

Since the dendritic sums follow discrete distributions (see Eqs. (8) and (9)), their CDFs 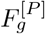 and *F*_*s*_ are not invertible. If we approximate the distributions by appropriate continuous distributions, Eqs. (17) and (16) can be reduced to a single equation

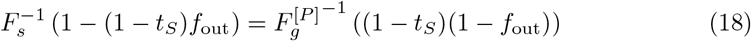

in *t*_*S*_. The capacity *P*\* corresponds thus to the number of patterns *P* for which the 1 − (1 − *t*_*S*_)*f*_out_-quantile of the spurious distribution equals the (1−*t*_*S*_)(1−*f*_out_)-quantile of the genuine distribution (see Fig 2F). An analytical solution to Eq. (18) is derived in the Methods.

### Effect of activation ratio on optimal learning speed and capacity

Let us now quantify how the memory capacity *P*\* depends on the activation ratio of the input pattern *f*_in_, the activation ratio of the output pattern *f*_out_, and on the transition probability *η*. Fig 3A shows the signal quality as a function of the number of patterns *P* for several transition probabilities. The signal quality always decays with time, i.e., with the number of subsequently stored patterns. The larger the transition probability, the faster the decay because the number of connections that is updated in each time step is larger and the old memories are thus overwritten more quickly. In the long run, the signal quality decays to zero. The initial signal quality (shown at *P* = 0 in Fig 3A) strongly depends on the transition probability because it determines how many connections are used for the initial memory storage. For a higher transition probability, the memory is more strongly engraved into the system and, initially, can be retrieved better. If the transition probability is too low, too few connections are updated to recall a memory even right after storing it (grey circles in Fig 3A). The transition probability that yields the highest capacity, which we call the optimal transition probability, hence depends on a trade-off between a high initial encoding and a slow decay of the memory signal (Fig 3A).

**Fig 3.**
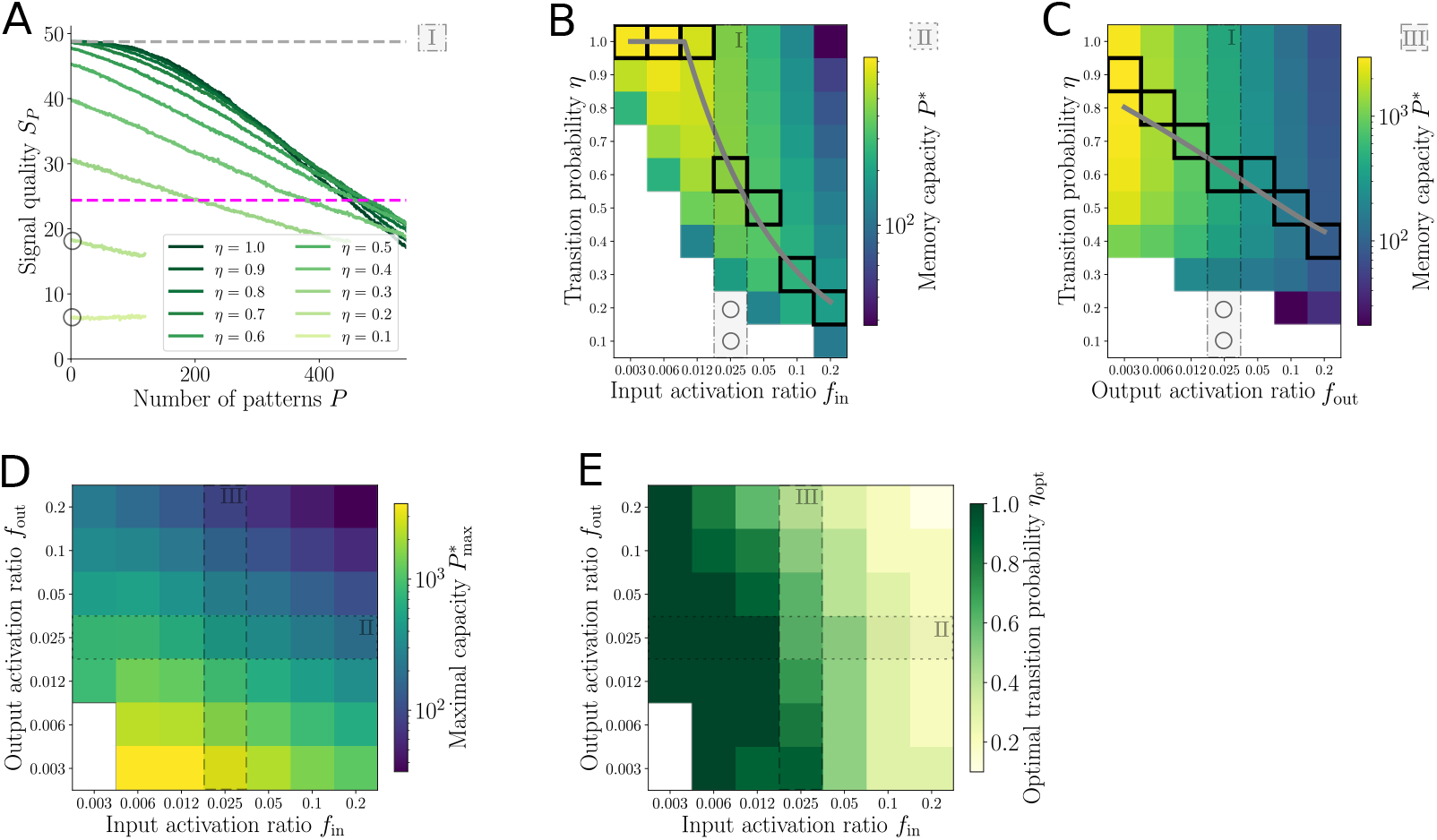
Effect of activation ratio on optimal learning speed and capacity. (**A**) Signal quality decaying with number of patterns *P* for various transition probabilities *η*. While small *η* values cause low initial signal quality, large *η* values cause a fast decay. A transition probability of *η* = 0.6 yields the largest memory capacity because it reaches the retrieval threshold (magenta dashed line), which is chosen as half of the maximal signal quality (grey dashed line), for the largest number of patterns *P*. For approximately *η <* 0.25, the initial signal quality is lower than the threshold and the capacity is zero (grey circles, see also grey circles in (**B**) and (**C**)). (Parameter values: *f*_in_ = *f*_out_ = 0.025.) (**B**) For a given transition probability, the memory capacity increases with decreasing input activation ratio *f*_in_ until it reaches a maximum and then quickly decays to zero for smaller input activation ratios. For a fixed input activation ratio *f*_in_, the optimal transition probability, which is defined as the transition probability that results in the largest memory capacity for the given set of network parameters, increases with decreasing *f*_in_ until it reaches and stays at its largest value of 1 (see black squares). The solid grey line shows the analytically derived optimal transition probability as a function of the input activation ratio *f*_in_. The signal quality traces of the column marked by I are shown in (**A**). White squares indicate a capacity of zero. (Output activation ratio *f*_out_ = 0.025.) (**C**) Same as (**B**) for output activation ratio *f*_out_ instead of input activation ratio *f*_in_. The memory capacity and the optimal transition probability increase with decreasing output activation ratio *f*_out_. Column I is shown in (**A**). (Input activation ratio *f*_in_ = 0.025.) In (**D**), the maximal capacity (obtained with the optimal transition probability *η* shown in (**E**)) for each combination of input activation ratio *f*_in_ and output activation ratio *f*_out_ is shown. For a fixed input activation ratio *f*_in_, the maximal capacity increases monotonically with decreasing *f*_out_. For a fixed output activation ratio *f*_out_, the maximal capacity has a maximum at a small but nonzero *f*_in_. Columns II and III are obtained from the results shown in (**B**) and (**C**), respectively. (**E**) Optimal transition probability increases with decreasing *f*_in_ and with decreasing *f*_out_. The slope is larger for *f*_in_ than *f*_out_. Columns II and III are shown in (**B**) and (**C**), respectively. Further parameter values in (**A**) - (**E**): *N*_in_ = 1000, *N*_out_ = 1000, *c* = 0.2, *c*_*m*_ = 1, *t*_*S*_ = 0.5, *N*_avg_ = 200 (see Methods for details on averaging).

Fig 3B summarizes how the memory capacity *P*\* depends on the input activation ratio *f*_in_ and the transition probability *η* — for fixed *f*_out_. The capacity values of Fig 3A are visible in the marked column (*f*_in_ = 0.025) of Fig 3B. In each column of Fig 3B, hence for a fixed *f*_in_, the largest capacity value that thus corresponds to the optimal transition probability is marked by a black square. The corresponding analytical estimate of the optimal transition probability is displayed by the grey line (Eq. (165) in Methods) For a fixed transition probability *η* (single row in Fig 3B), the capacity *P*\* is non-monotonic as a function of *f*_in_, i.e., it increases with increasing input activation ratio *f*_in_ and then decreases again. As known from the literature [33, 35, 37], denser representations (higher *f*_in_) can lead to lower capacity values. However, if the input representations become too sparse (too small *f*_in_), the maximal possible capacity also drops and goes to zero for very sparse input patterns. This can be explained by the fact that it is not possible for too few active input units to sufficiently excite the respective output units. In this case, a slight change of the weight matrix due to the storage of additional patterns can lead to very different units being activated in the output. In extreme cases — especially if both the transition probability *η* and the input activation ratio *f*_in_ are too small — the pattern is not well enough imprinted from the very beginning (cf. Fig 3A, grey circles, and 3B, white region) and the capacity is zero. This issue is particularly severe if the error tolerance 1 − *t*_*S*_ is small, hence if the retrieval threshold *T*_*S*_ is high, i.e., is close to its maximum *H*_avg_ (not shown here). In this case, the capacity can be zero even for high transition probabilities *η* if the initial signal quality lies below *T*_*S*_.

For a fixed transition probability (single rows in Fig 3C), the capacity increases with decreasing output activation ratio *f*_out_. In Fig 3C, the capacity is evaluated for several output activation ratios *f*_out_ and transition probabilities *η* while the input activation ratio *f*_in_ is fixed. The marked column in Fig 3C shows the same capacity values as the marked column in Fig 3B. A smaller output activation ratio *f*_out_ is beneficial for the capacity because a specific output unit is less likely to be active in subsequent patterns; hence relevant connections are turned off with lower probability. Nevertheless, if the transition probability *η* is too small, as before, the memory is not sufficiently well imprinted from the beginning and the capacity is zero (Fig 3C, white region).

In summary, we find that the sparseness of the input and the output patterns, respectively, have differential effects on the maximal capacity of the network. While the smallest output activation ratio achieves the highest capacity values (Fig 3C), there is a non-monotonic dependence of the maximal capacity on the input activation ratio (Fig 3B).

The non-monotonic dependence of the capacity on *f*_in_ and its monotonic dependence on *f*_out_ are also maintained for the maximal capacity. Fig 3D summarizes the maximal capacity for various pairs of input and output activation ratios *f*_in_ and *f*_out_, which is obtained by using the optimal transition probability for each pair. As for the capacity for a fixed *η* in Fig 3B and C, Fig 3D shows an asymmetric effect of *f*_in_ and *f*_out_ on the maximal capacity.

Furthermore, we find that the optimal learning speed depends on the input as well as the output activation ratio. The sparser the patterns (small *f*_in_ or *f*_out_), the larger the transition probability that allows for the largest capacity (depicted by black squares in Fig 3B and C). This effect is monotonic and has a larger gradient for the input activation ratio (Fig 3B) than the output activation ratio (Fig 3C), which is summarized in Fig 3E for various pairs of *f*_in_ and *f*_out_.

In S4 Appendix, we compare our results with post-synaptic homeostasis to a pre-synaptic normalization and a normalization averaged over all connections and find that the qualitative dependences of the optimal transition probability and the maximal capacity on the activation ratios (compare Fig 5D,E to S4 Appendix) remain unchanged.

### Activation ratios in the distributions of dendritic sums

The intuitive understanding of these qualitatively different effects of the input and the output activation ratios on the capacity can be strengthened by having a closer look at the distributions of the dendritic sums, which determine the signal quality of the memory. We again distinguish distributions of dendritic sums of spurious and genuine units (red and blue, respectively, in Fig 2A) as given in Eq. (8) and Eq. (9), respectively. These explicit expressions for dendritic sums of spurious and genuine units allow us to understand why *f*_in_ and *f*_out_ affect the capacity of the network in different ways.

We first explain the role of *f*_in_, while *f*_out_ is assumed to be fixed: A fixed *f*_out_ implies that the weighting of the individual binomial mass functions in Eq. (9) is fixed for a given *P*. We can hence discuss the role of *f*_in_ in the individual binomials without considering the weighting. The capacity’s non-monotonic dependence on *f*_in_ is a combination of two separate effects working together. In the following, I discuss via what dominating mechanism *f*_in_ can positively correlate with the capacity, which is the case for very small *f*_in_ (left of the grey line in Fig 3B), and what mechanism has to dominate such that *f*_in_ is negatively correlated with the capacity, which is the case for larger *f*_in_ (right of the grey line in Fig 3B). First, the input activation ratio occurs in the number of trials *M*_in_ = *f*_in_ *N*_in_ of the individual binomial distributions in 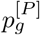 and the number of trials *M*_in_ of the spurious distribution *p*_*s*_. Let us assume for the moment that *f*_in_ was increased only in the number of trials of the binomials, and we ignore the dependence of *ρ*_*g*_ on *f*_in_ in Eq. (10). Such an increase of *f*_in_ in the number of trials pulls the distributions *p*_*g*_ and 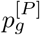 farther apart since both mean values are proportional to *f*_in_. They are also stretched out more (higher standard deviation 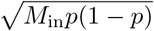 with *p* = *c* or *p* = *ρ*_*g*_(*u*)) but the overall overlap of the two distributions is reduced because the means grow with *M*_in_ and the standard deviations grow only with 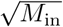 (also see left hand side of Eq. (133) for *P* = 0). This is illustrated in Fig 4A (top: smaller *f*_in_, bottom: larger *f*_in_). Due to the smaller overlap of the two distributions, the Hamming distance is reduced and this yields a larger capacity. For *f*_in_ increasing from zero, this increase of the capacity can be seen for the values left of the grey line in Fig 3B. However, the input activation ratio *f*_in_ also occurs in the probabilities *ρ*_*g*_(*u*) (Eq. (10)) of the individual binomials in the genuine distributions. If only changed at this place, an increased *f*_in_ leads to a faster decay of the probability with increasing *u* (Fig 4B) and hence with the number of patterns *P*. For larger *f*_in_, each of the individual binomials 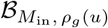 in Eq. (9) and hence also the total genuine distribution converges faster to the spurious distribution, which implies a deteriorative effect on the capacity. This can be seen for the capacity values right of the grey line in Fig 3B. In summary, these two effects of *f*_in_ interact with each other, which can explain the intermediate optimal value of *f*_in_ where the first effect is balanced by the second one to allow for an optimal capacity. For input activation ratios *f*_in_ close to zero, the effect of the change of *f*_in_ in the number of trials of the distributions (8) and (9) dominates over the effect of the change of *f*_in_ in (10). The capacity hence increases with increasing *f*_in_ if *f*_in_ is small (derived in S3 Appendix). This confirms the detrimental effect of a very small *f*_in_ on the network capacity that we discussed in the previous section: If *f*_in_ becomes very small, the number of active input units becomes too low to reliably activate the respective output units. For very large input activation ratios *f*_in_, the effect of a change of *f*_in_ via the probability *ρ*_*g*_ (Eq. (10)) is stronger than the effect via the number of trials and the capacity decreases with increasing *f*_in_.

**Fig 4.**
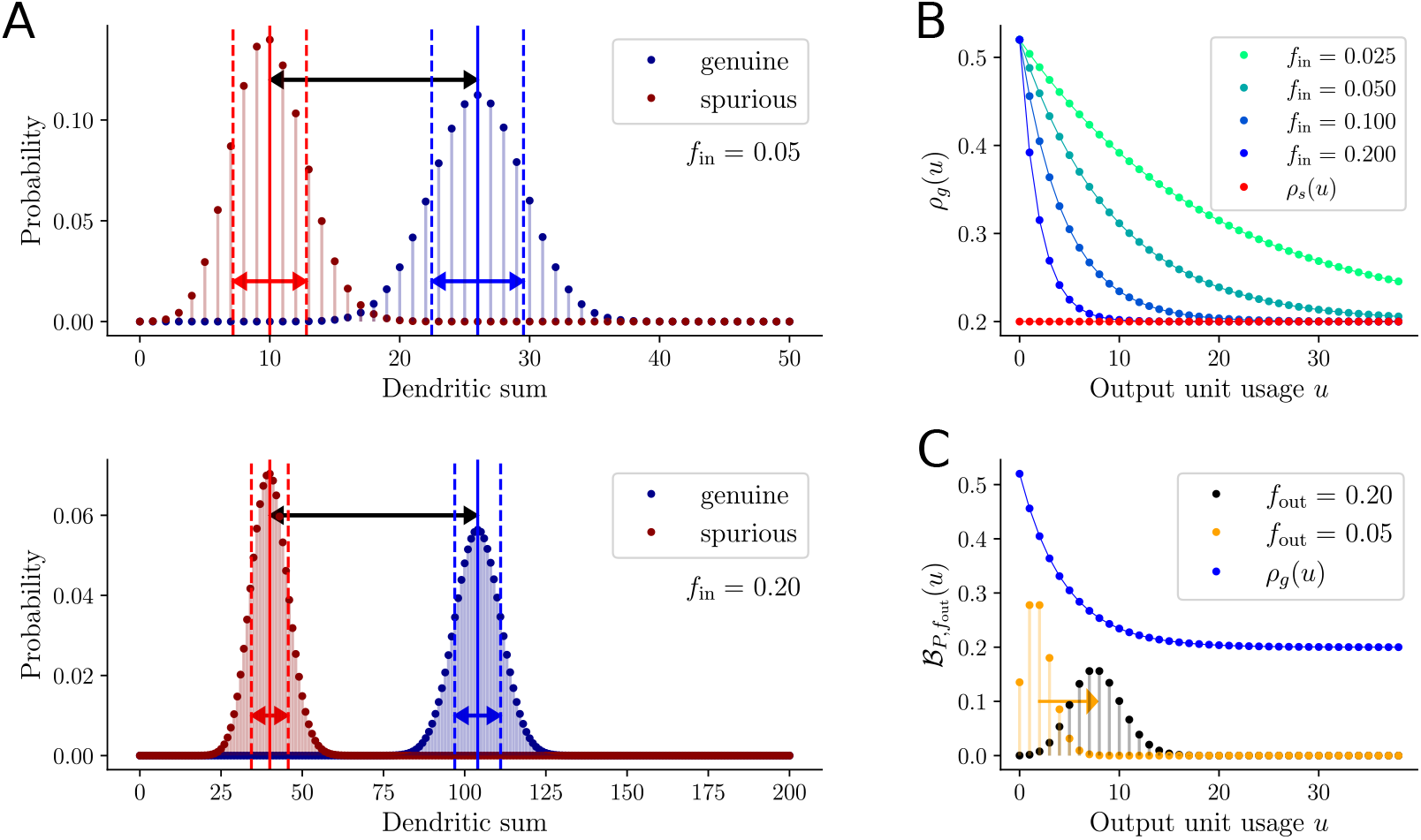
Role of the activation ratios in the distributions of dendritic sums. **(A)** Distributions of dendritic sums of genuine (blue) and spurious (red) output units. In the bottom panel, *f*_in_ is four times larger than in the top panel. For a larger input activation ratio *f*_in_, the difference between the means of the distributions (black arrows) is larger. The standard deviations (red and blue arrows) are also larger for larger *f*_in_ (note the different scales of the *x*-axes in the two panels) but the increase is less than for the means. The overall overlap of the two distributions is smaller for larger *f*_in_. (**B**) The probability *ρ*_*g*_ of a connection to a genuine output unit to be functional as a function of the output unit usage *u* decays faster and hence approaches *ρ*_*s*_ (red) faster for larger input activation ratio *f*_in_. (**C**) The weight distribution 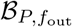 moves to the right if *f*_out_ is increased (black: *f*_out_ four times larger than in orange). The largest weights hence affect binomials 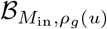 (in Eq. (9)) with smaller *ρ*_*g*_(*u*) values (blue). Further parameters: *N*_in_ = 1000, *f*_out_ = 0.1, *c* = 0.2, *c*_*m*_ = 1, *P* = 0, *η* = 0.4.

**Fig 5.**
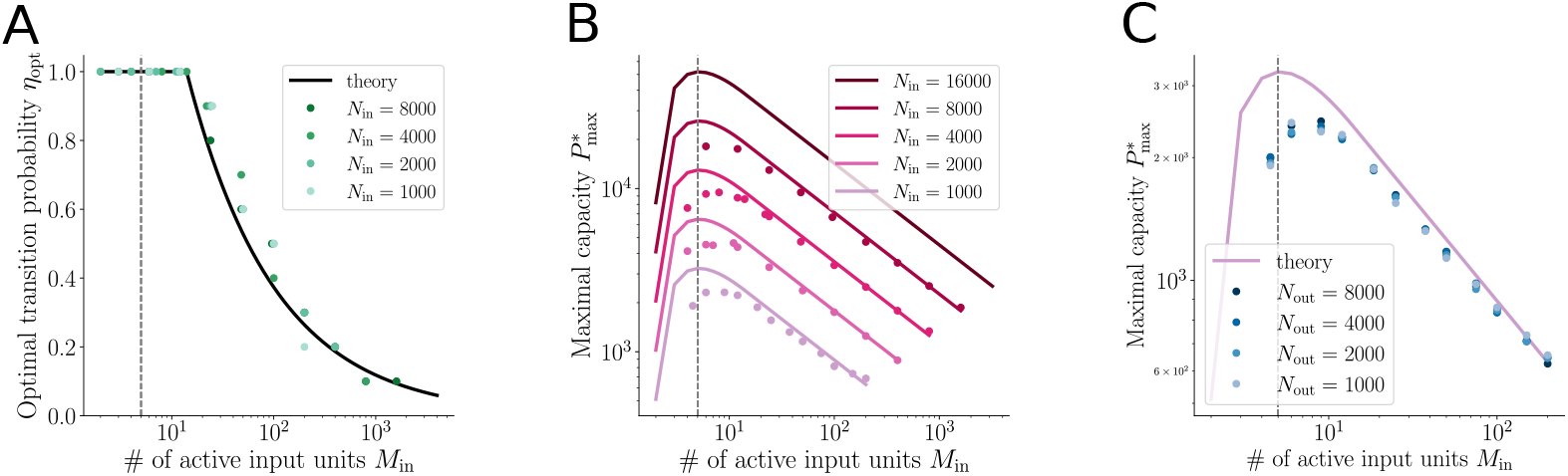
Role of network size and number of active units. Results from numerical simulations are depicted by dots, theoretical results are depicted by solid lines. (**A**) The optimal transition probability *η*_opt_ as a function of *M*_in_ monotonically decreases from a plateau at one to zero. It does not depend on the input layer size *N*_in_. The overall highest capacity for a fixed network size is obtained for an *M*_in_ which yields an optimal *η* of one (grey dashed line). (**B**) shows the maximal capacity 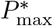 as a function of the number of active input units *M*_in_ = *f*_in_*N*_in_ for *N*_in_ = 1000, 2000, 4000, 8000 and 16000. The size of the output layer is fixed to *N*_out_ = 1000 in the simulations. The maximal capacity as a function of *M*_in_ scales linearly with *N*_in_. The grey dashed line marks the number of active input units *M*_in_ that yields the highest capacity. The highest capacity for a fixed network size is obtained at the same *M*_in_ for any network size. In (**C**), the output layer size *N*_out_ (= 1000, 2000, 4000, 8000) is varied instead of the input layer size *N*_in_. The number of input units *N*_in_ is fixed to 1000. The size of the output layer *N*_out_ does not have an impact on the maximal memory capacity 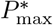 of the network. Further parameter values in (**A**)-(**C**): *f*_out_ = 0.006, *c* = 0.2, *c*_*m*_ = 1, *t*_*S*_ = 0.5, *N*_avg_ = 200.

The upper limit of *f*_in_ is *c* (see first section on condition *f*_in_ ≤ *c*). The capacity for this upper limit of the input activation ratio *f*_in_ can be derived for the special case *c*_*m*_ = 1, *η* = 1. If *f*_in_ takes its maximal value *c*, the probability *ρ*_*g*_(*u*) (Eq. (10)) equals *c* and the genuine (Eq. (9)) and spurious distributions (Eq. (8)) become identical. This makes it impossible to differentiate between genuine and spurious units and the capacity is hence zero. In total, the intuition that we have developed here thus matches the numerical results discussed in the previous section.

Let us now turn to the output activation ratio *f*_out_, which affects the capacity also in two ways. We now assume a fixed *f*_in_. First, *f*_out_ is the success probability of the binomial distribution, which constitutes the weights in the genuine distribution (Eq. (9)). An increased *f*_out_ pushes the center of mass of the distribution to larger values of *u* (Fig 4C). The largest weights thus are given to larger *u* values, which correspond to binomials with smaller probabilities *ρ*_*g*_(*u*). This emphasizes the parts of 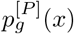 that are more to the left of *x* ∈ [0, *M*_in_], which culminates in a shift to the left of the center of mass of the overall genuine distribution. In this sense, increasing *f*_out_ increases the overlap of the genuine distribution with the spurious distribution and hence decreases the capacity of the network. At the same time, *f*_out_ directly impacts the placement of the activation threshold because we are always enforcing *M*_out_ = *f*_out_*N*_out_ active output units. This is discussed in detail in the Methods. The effect on the capacity of a change of the activation threshold due to an increase of *f*_out_ is unknown (also see Methods). We speculate that this effect is small in the parameter range that we investigated numerically and that the effect of *f*_out_ via the weight distribution discussed above dominates the dependence of the capacity on *f*_out_.

### Role of network size and number of active units

So far, we have considered a fixed network size. We have seen how the capacity and the optimal transition probability depend on the input and the output activation ratio. Here, we discuss how these results depend on network sizes.

Assuming that the average number of functional connections per output unit is large but not too large (*M*_in_*c* ≫ 1 and *M*_in_(1 − *ρ*_*g*_(⌊*f*_out_(*P* + 1) ⌋)) ≫ 1) and *f*_in_*ηc*_*m*_*/c* ≪ 1, the capacity *P*\* of the network can be analytically approximated as

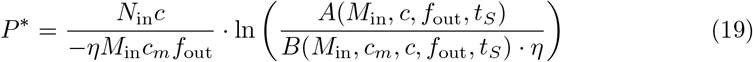

with *A* and *B* being two functions that are defined in Equations (147) and (148) in the Methods section (derived in Methods, see Eq. (159)). Note that both *A* and *B* are independent of the input and output network size *N*_in_ and *N*_out_. They only depend on the number of active input units *M*_in_ and the output activation ratio *f*_out_.

Under the assumption 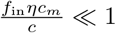, the optimal transition probability can be approximated by

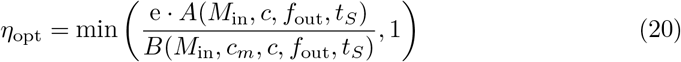

(black line in Fig 5A, derived in Methods, see Eq. (165)). It is independent of the input layer size *N*_in_ and the output layer size *N*_out_. The maximal capacity 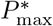 for a fixed set of network parameters is obtained by using the optimal transition probability *η*_opt_ in Equation (19):

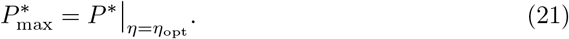

We find analytically and numerically that the size of the input layer strongly impacts the maximal capacity of the network (see Fig 5B): The capacity *P*\* scales linearly with the input layer size *N*_in_. Since *η*_opt_ is independent of *N*_in_, the maximal capacity 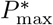 also scales linearly with *N*_in_ (Eq. (19) and Fig 5B). The maximal capacity 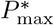 depends non-monotonically on the number of active input units *M*_in_. We find that, for any input layer size, the highest capacity is reached for the same number of active input units *M*_in_ (Fig 5B, grey line). According to Eqs. (19) and (21)), the maximum over *M*_in_ of the maximal capacity 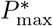 depends on the output activation ratio *f*_out_, the functional and morphological connectivities *c* and *c*_*m*_ and of course, via *t*_*S*_, also on the retrieval threshold *T*_*S*_ = *t*_*S*_*H*_avg_. Since, for a fixed network size *N*_in_, the input activation ratio *f*_in_ = *M*_in_*/N*_in_ is simply a scaled version of *M*_in_, the maximal capacity 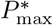 also depends non-monotonically on *f*_in_. We have seen examples of this non-monotonic dependence on *f*_in_ in Fig 3D. An increase of the retrieval threshold *T*_*S*_ yields a larger optimal transition probability and, of course, a smaller capacity (S1 Fig). If the morphological connectivity *c*_*m*_ is increased, *η*_opt_ decreases while 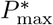 increases (S2 Fig). An increase of the functional connectivity *c* increases both *η*_opt_ and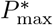, at least for large enough *M*_in_ (S3 Fig).

As a probability, the optimal transition probability is bounded between zero and one. It is a monotonically decreasing function of *M*_in_. For small values of *M*_in_, it takes the maximal value one. When *M*_in_ becomes large enough, the expression e*A/B* becomes smaller than one and *η*_opt_ becomes strictly decreasing as a function of *M*_in_ (Fig 5A). This monotonicity of *η*_opt_ as a function of *M*_in_ could not be formally shown with our theory but has been confirmed in all evaluations of Equation (20) and all numerical simulations that were performed. In the range where *η*_opt_ *<* 1, the maximal capacity 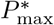 is monotonically decreasing as a function of *M*_in_ (shown in the Methods) (compare Fig 5A and B). It follows that, if the maximal capacity has a maximum as a function of *M*_in_ (for any given network size), it is reached within the range where we have *η*_opt_ = 1 (Fig 5A, the grey line marks the *M*_in_ that yields the largest capacity). Note that, depending on the network parameters and the retrieval criterion, it can occur that there is no *M*_in_ ≥ 1 for which *η*_opt_ = 1. In this case, the dependence of the maximal capacity on *M*_in_ (as well as *f*_in_) is monotonic and the largest capacity is obtained for *M*_in_ = 1.

As can be seen from Eqs. (19) and (21) and as discussed in previous sections, the capacity depends on the output activation ratio *f*_out_. For adequately large *N*_out_, the size of the output layer, on the other hand, does not affect the storage capacity of the network (Eq. (19) and Fig 5C, also see Discussion.

### Noisy input patterns during retrieval

We previously (Fig 3A) found that the transition probability that leads to the largest capacity depends on a balance between a large initial signal quality and a slow decay of the signal quality with the number of subsequently stored patterns. The top panel of Fig 6A also shows the signal quality as a function of the number of patterns but for a different set of input and output parameters (*f*_in_ = *f*_out_ = 0.1) than Fig 3A. Let us now investigate the effect of noisy input patterns during retrieval on the optimal transition probability and the maximal capacity of the network.

**Fig 6.**
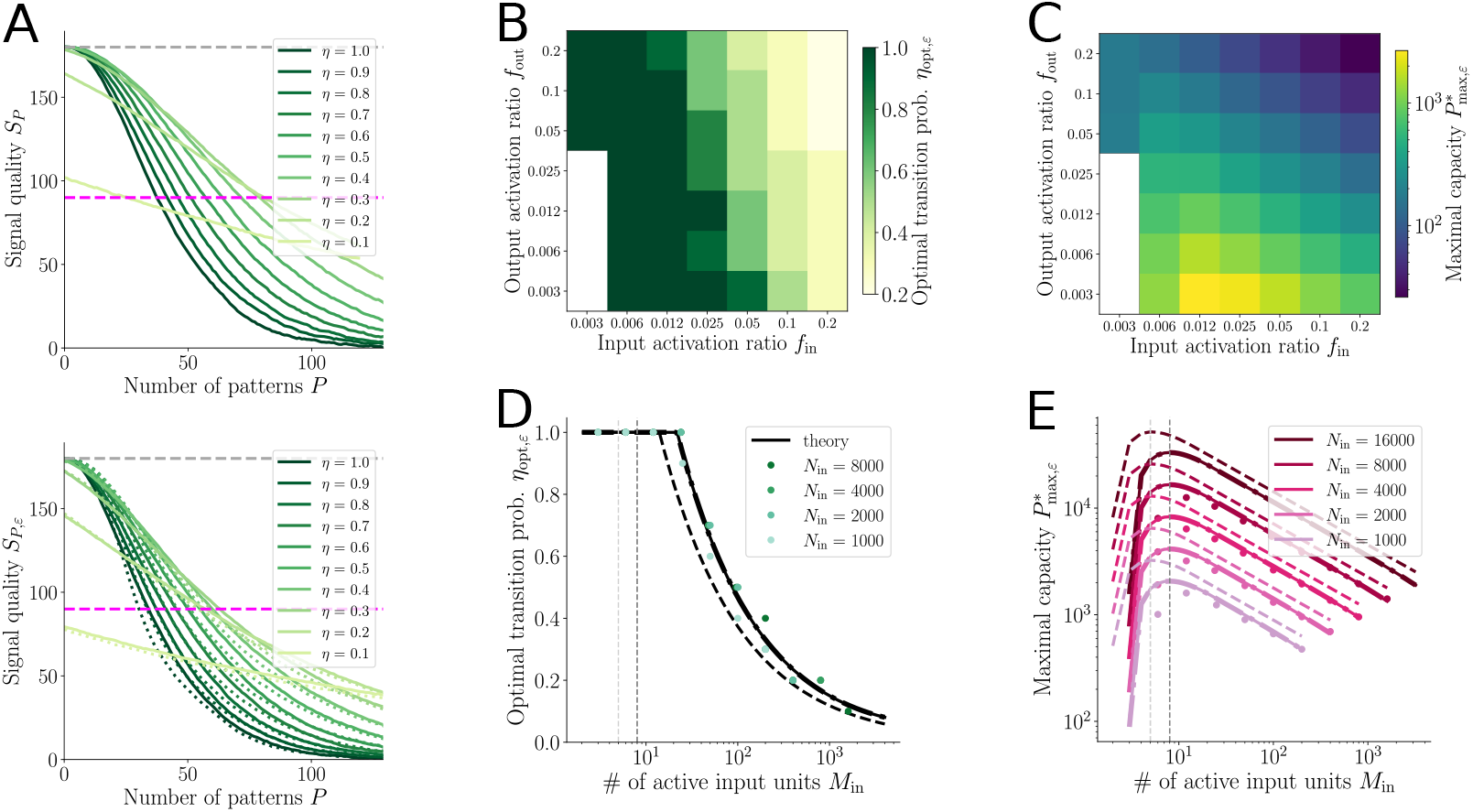
With noisy input patterns during retrieval, maximal capacity decreases and optimal transition probability increases. The noise level is set to *ε* = 0.2 in the entire figure. (**A**) The memory signal quality decays with the number of patterns learned by the network. For smaller transition probabilities *η*, the signal quality *S*_*P*_ starts lower but decays more slowly than for larger *η*. Top: No noise during retrieval. The largest capacity is attained for *η* = 0.2. Bottom: With noise on the input patterns during retrieval. With noise, the signal quality *S*_*P,ε*_ is lower than without noise, particularly for small transition probabilities *η*. All capacity values with noise are slightly smaller than the corresponding ones without noise. The largest capacity is now attained for *η* = 0.3. In (**A**) the solid lines show results from simulations. The dotted lines are an approximation of the signal quality with noise *S*_*P,ε*_ where the signal quality without noise *S*_*P*_ is multiplied by a factor obtained from theory (see S2 Appendix). In (**A**), *f*_in_ = *f*_out_ = 0.1. (**B**)-(**C**) The optimal transition probability *η*_opt,*ε*_ and the maximal memory capacity 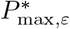 as functions of input and output activation ratios *f*_in_ and *f*_out_ obtained with noise during retrieval behave qualitatively similar to the maximal memory capacity 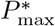 and the optimal transition probability *η*_opt_ obtained without noise (cf. Fig 3D and E). In (**A**)-(**C**), *N*_in_ = *N*_out_ = 1000. (**D**) The optimal transition probability *η*_opt,*ε*_ with noise (solid line, Eq. (22)) is slightly higher than without noise (dashed line, Eq. (20)). The dash-dotted line shows an approximation of *η*_opt,*ε*_ as a multiple of *η*_opt_ (Eq. (24)). The dark grey and light grey lines mark the *M*_in_ that yields the highest capacity with and without noise, respectively. With noise, the optimal *M*_in_ is larger than without noise. All black lines are derived from theoretical considerations, the colored dots show results from simulations (*N*_out_ fixed to 1000). (**E**) Noise on the input patterns during retrieval reduces the maximal capacity. The maximal capacity scales with the input layer size *N*_in_. Dashed lines show the maximal capacity without noise (Eq. (21)), solid lines show the maximal capacity with noise (Eq. (25)), dash-dotted lines show an approximation of the maximal capacity with noise for small *ε* (Eq. (26)) and dots show numerical results (*N*_out_ fixed to 1000). Further parameter values in (**A**)-(**E**): *c* = 0.2, *c*_*m*_ = 1, *t*_*S*_ = 0.5, *N*_avg_ = 200.

The input pattern that is presented to the network during the retrieval phase does not necessarily have to be the same pattern that was used for training. From a biological perspective, it makes sense to assume that the input during retrieval is not exactly the same but a noisy version of the original input pattern. The noisy input patterns are created by deactivating a particular fraction *ε* of the active input units while activating the same number of inactive input units. The input activation ratio *f*_in_ is hence maintained. Adding noise to the input pattern during retrieval reduces the memory capacity by deteriorating the signal quality right from the beginning (Fig 6A, top vs bottom). This shift in the initial signal quality is particularly strong for small transition probabilities *η* because the memory relies on few connections to provide a strong input to the output units. If some of the input units that are supposed to be active in a pattern are inactive and others are activated instead, it is likely that many of the genuine output units are not activated anymore. The effect is less drastic for large transition probabilities for which the memory is more reliably imprinted into many connections and does not get lost just because some input units are wrongly activated. However, we find that the change in the signal quality due to noise can neither be explained by a constant shift nor by a constant multiplicative scaling. The change depends on the number of patterns in a more complex way, see S2 Appendix. There we derived an analytical description of the factor that has to be applied to the signal quality *S*_*P*_ calculated without noise during retrieval to obtain an estimate of the signal quality *S*_*P,ε*_ with noise.

Despite the complex relationship between *S*_*P*_ and *S*_*P,ε*_, the qualitative dependence of the optimal transition probability *η*_opt,*ε*_ on the input and output activation ratios remains the same as without noise (Fig 6B, compare to Fig 3E) — it increases with decreasing activation ratios. Further, adding noise to the input pattern during retrieval maintains the non-monotonic effect of the input activation ratio and the monotonic effect of the output activation ratio on the maximal capacity (Fig 6C, compare to Fig 3D).

We extend our theory to retrieval with noisy input patterns and find that the optimal transition probability can again be described as

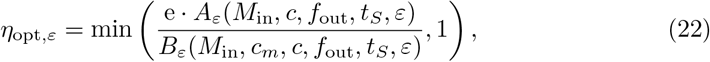

where *A*_*ε*_ and *B*_*ε*_ (Eqs. (182) and (183)) now additionally depend on the noise level *ε* (derived in Methods). For a fixed set of parameters, noise during retrieval increases the optimal transition probability:

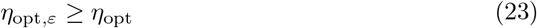

(Fig 6D, compare solid line to dashed line). It can be approximated as a scaled version of the optimal transition probability without noise:

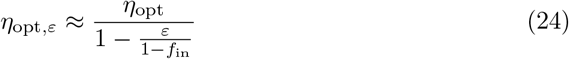

(Fig 6D, dash-dotted lines, also see S4 Fig. As discussed in previously, the optimal transition probability corresponds to a sweet spot that realizes both a good initial encoding of the memory and a slow decay of the memory trace. The additional noise impacts both aspects but since we find that *η*_opt,*ε*_ ≥ *η*_opt_, the noise has a more detrimental effect on the initial signal quality than on the decay of the signal quality.

The maximal capacity can again be theoretically described as

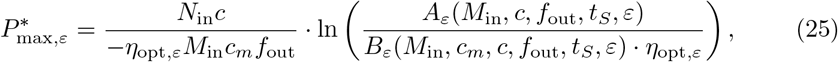

(Fig 6E, solid lines, derived in Methods, see also S4 Fig). It can further be approximated by

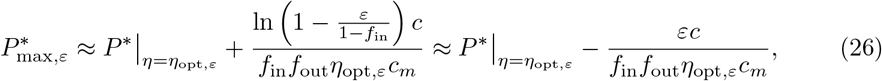

where 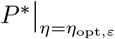 is the capacity without noise (Eq. (19)) evaluated for *η* = *η*_opt,*ε*_ and the last approximation step assumes *ε* ≪ 1 and *f*_in_ ≪ 1 (Fig 6E, dash-dotted lines, see also S4 Fig; derived in Methods). Hence, the maximal capacity with noise during retrieval is the maximal capacity without noise (evaluated at *η*_opt,*ε*_) shifted down by a fixed amount that scales linearly with *ε*. The *M*_in_ that yields the highest capacity is again independent of *N*_in_ but it increases with the noise level *ε* (Fig 6E, vertical dashed lines, see also S4 Fig).

We can conclude that in a system with noisy input patterns, the capacity is lower than in a system without noise. However, with noise, it is beneficial for the capacity to learn with a higher transition probability.

## Methods

In the following, we give a detailed account of the analytical derivations and numerical implementations used to obtain the presented results. First, we derive the distributions of dendritic sums, which provides the basis for the subsequent derivation of analytical expressions for the memory capacity. Finally, we describe the numerical implementation of the algorithm. Table 2 provides a summary of the main parameters.

**Table 1.** Deviation between theory and simulations: Maximal capacity obtained from theory divided by maximal capacity obtained from simulations for various values of *N*_in_ and *f*_out_ averaged across all *M*_in_ values shown in Fig 22 except the two smallest *M*_in_ = 3 and *M*_in_ = 6 (which does not fulfill the condition *M*_in_ ≫ 1 and for which the fit is thus not accurate enough). Deviation calculated based on Eq. (152) (corresponding to dashed lines in Fig 22). In brackets: Deviation calculated based on (159), i.e. with the Taylor approximation (156) (corresponding to solid lines in Fig 22).

| | $f_{\text{out}} = 0.1$ | $f_{\text{out}} = 0.025$ | $f_{\text{out}} = 0.006$ |
| --- | --- | --- | --- |
| $N_{\text{in}} = 1000$ | 1.21 (1.36) | 1.34 (1.53) | 1.44 (1.67) |
| $N_{\text{in}} = 2000$ | 1.13 (1.21) | 1.22 (1.32) | 1.33 (1.45) |
| $N_{\text{in}} = 4000$ | 1.11 (1.16) | 1.16 (1.22) | 1.22 (1.28) |
| $N_{\text{in}} = 8000$ | 1.06 (1.09) | 1.11 (1.14) | 1.24 (1.18) |

**Table 2.**
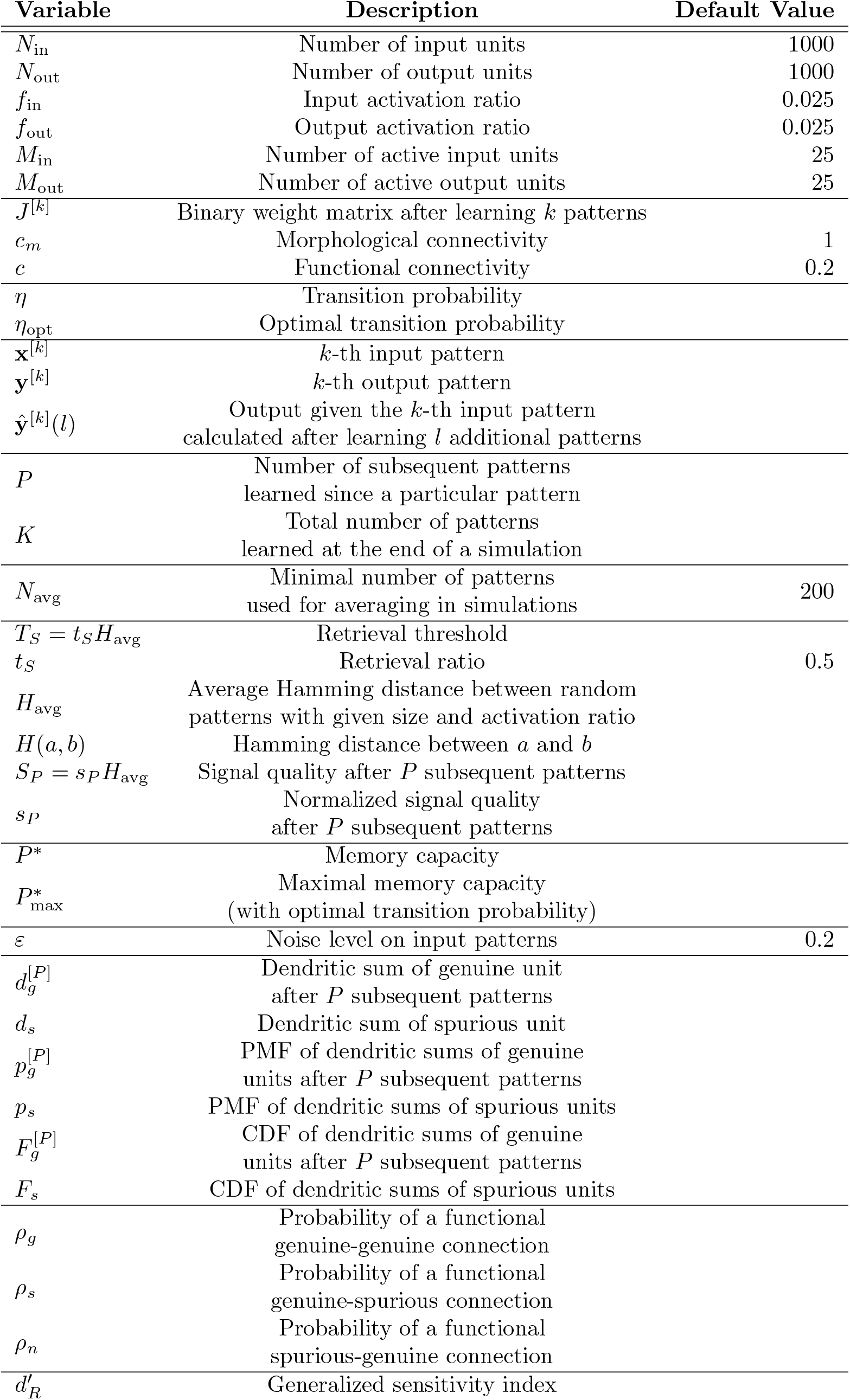
Description of variable names and their default values.

| Variable | Description | Default Value |
| --- | --- | --- |
| $N_{\text{in}}$ | Number of input units | 1000 |
| $N_{\text{out}}$ | Number of output units | 1000 |
| $f_{\text{in}}$ | Input activation ratio | 0.025 |
| $f_{\text{out}}$ | Output activation ratio | 0.025 |
| $M_{\text{in}}$ | Number of active input units | 25 |
| $M_{\text{out}}$ | Number of active output units | 25 |
| $J^{[k]}$ | Binary weight matrix after learning $k$ patterns | |
| $c_m$ | Morphological connectivity | 1 |
| $c$ | Functional connectivity | 0.2 |
| $\eta$ | Transition probability | |
| $\eta_{\text{opt}}$ | Optimal transition probability | |
| $\mathbf{x}^{[k]}$ | $k$-th input pattern | |
| $\mathbf{y}^{[k]}$ | $k$-th output pattern | |
| $\hat{\mathbf{y}}^{[k]}(l)$ | Output given the $k$-th input pattern<br>calculated after learning $l$ additional patterns | |
| $P$ | Number of subsequent patterns<br>learned since a particular pattern | |
| $K$ | Total number of patterns<br>learned at the end of a simulation | |
| $N_{\text{avg}}$ | Minimal number of patterns<br>used for averaging in simulations | 200 |
| $T_S = t_S H_{\text{avg}}$ | Retrieval threshold | |
| $t_S$ | Retrieval ratio | 0.5 |
| $H_{\text{avg}}$ | Average Hamming distance between random<br>patterns with given size and activation ratio | |
| $H(a, b)$ | Hamming distance between $a$ and $b$ | |
| $S_P = s_P H_{\text{avg}}$ | Signal quality after $P$ subsequent patterns | |
| $s_P$ | Normalized signal quality<br>after $P$ subsequent patterns | |
| $P^*$ | Memory capacity | |
| $P_{\text{max}}^*$ | Maximal memory capacity<br>(with optimal transition probability) | |
| $\varepsilon$ | Noise level on input patterns | 0.2 |
| $d_g^{[P]}$ | Dendritic sum of genuine unit<br>after $P$ subsequent patterns | |
| $d_s$ | Dendritic sum of spurious unit | |
| $p_g^{[P]}$ | PMF of dendritic sums of genuine<br>units after $P$ subsequent patterns | |
| $p_s$ | PMF of dendritic sums of spurious units | |
| $F_g^{[P]}$ | CDF of dendritic sums of genuine<br>units after $P$ subsequent patterns | |
| $F_s$ | CDF of dendritic sums of spurious units | |
| $\rho_g$ | Probability of a functional<br>genuine-genuine connection | |
| $\rho_s$ | Probability of a functional<br>genuine-spurious connection | |
| $\rho_n$ | Probability of a functional<br>spurious-genuine connection | |
| $d'_R$ | Generalized sensitivity index | |

First, we briefly recap the network model and the learning paradigm. The network is presented with a sequence of input/output-pattern pairs (**x**^[*k*]^, **y**^[*k*]^), *k* ∈ {0, 1, …}. Input and output patterns are binary vectors of length *N*_in_ and *N*_out_, respectively, and of activation ratio *f*_in_ and *f*_out_, respectively. Hence, they consist of *M*_in_ = *f*_in_*N*_in_ or *M*_out_ = *f*_out_*N*_out_ active units and *N*_in_ − *M*_in_ or *N*_out_ − *M*_out_ inactive units, respectively.

The input and the output layer of the network are connected by a weight matrix *J*. Per output unit, a number *c*_*m*_*N*_in_ connections (*c*_*m*_*N*_in_ entries per row of the weight matrix) are randomly chosen to be morphologically available connections. They can be functional (‘on’, value 1) or silent (‘off’, value 0). The functional connectivity is always normalized such that a fixed number *cN*_in_ of morphologically available connections per output unit is functional.

In each learning step, the weight matrix is updated according to the presented pattern pair (**x**^[*k*]^, **y**^[*k*]^). Silent connections between active input and active output units are turned on with transition probability *η*. In addition, we randomly silence the same number of functional connections from inactive input units to active output units such that the total number of functional connections per output unit is maintained. This update step transforms *J* ^[*k−*1]^ into *J* ^[*k*]^.

For any pattern pair that has already been learned, retrieval can be tested by presenting the original input **x**^[*k*]^ and calculating the output as

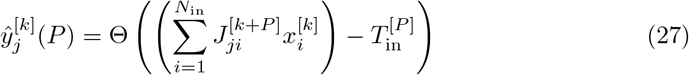

for *j* = 1, …, *N*_out_, where *P* is the number of subsequent patterns that have been learned since pattern *k*, and the activation threshold 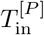 is chosen such that *M*_out_ output units are activated. To determine the decay of the memory signal, this calculated output **ŷ**^[*k*]^(*P*) can be compared to the given output pattern **y**^[*k*]^. The larger *P*, the larger the Hamming distance between the two vectors:

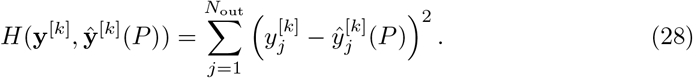

We define the signal quality after *P* subsequent patterns as

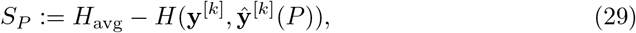

where *H*_avg_ = 2*N*_out_*f*_out_(1 − *f*_out_) is the average Hamming distance between two random *f*_out_-sparse vectors of length *N*_out_.

The retrieval threshold is defined as

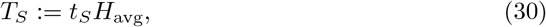

for a fixed *t*_*S*_ ∈ (0, 1). We call a pattern with *S*_*P*_ ≥ *T*_*S*_ retrievable and a pattern with *S*_*P*_ *< T*_*S*_ not retrievable. The maximal number *P* for which *S*_*P*_ ≥ *T*_*S*_, if *S*_*P*_ is averaged across many patterns, is defined as the capacity *P*\* of the network.

### Probabilistic description of dendritic sums

To better understand the learning dynamics of the network and the decay of the quality of the memory signal, which is used to quantify the capacity, we derive a probabilistic description of the dendritic sums of output units [69]. The dendritic sum of the *j*-th output unit is its net input

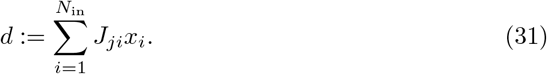

In accordance with [69], we introduce the following nomenclature: In general, a calculated output pattern 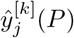 differs from the target output pattern 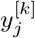. We call output units that are active (1) in the target output *genuine* output units and output units that are inactive (0) in the target output *spurious* units. In the calculated output, on the one hand, genuine units can be correctly active or incorrectly inactive. On the other hand, spurious units can be correctly inactive or incorrectly active. The terms *genuine* and *spurious* do not indicate whether the calculated activity of an output unit is 1 or 0, respectively, but whether it should be 1 or 0, respectively, to match the target output. In this Section, we derive the distributions of the dendritic sums of genuine and spurious output units separately, and these dendritic sums are denoted by symbols *d*_*s*_ (spurious) and *d*_*g*_ (genuine). The shapes of these two distributions (especially their overlap) is related to the Hamming distance between target and calculated output and consequently also to the signal quality. These relationships will become clear later in the Methods.

With respect to a particular pattern pair, we further distinguish between four types of connections:

- genuine-genuine (g-g) connections connect active input to active output units,
- genuine-spurious (g-s) connections connect active input to inactive output units,
- spurious-genuine (s-g) connections connect inactive input to active output units, and
- spurious-spurious (s-s) connections connect inactive input to inactive output units.

Note that the neuronal activity that determines the connection type is here understood as the activity in the original pattern pair (**x**^[*k*]^, **y**^[*k*]^) and not in the calculated output pattern **ŷ**^[*k*]^(*P*).

### Distributions of dendritic sums

In what follows, we calculate the distribution of the net input 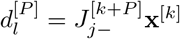 to an output unit *j* where *l* ∈ {*s, g*} denotes a spurious (*s*) or a genuine (*g*) output unit. This input 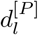 is also called the dendritic sum of output unit *j*. The term 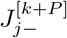 is the *j*-th line of the weight matrix obtained for *P* additional patterns after learning pattern *k*. To understand how the *P* subsequent patterns change the weight matrix, we have to take into account how often the output unit *j* is active across all *P* patterns, which is also called the output unit usage. As suggested by [69], we assume that the number *r* of times an output unit is active in *P* statistically independent patterns follows a binomial distribution ℬ(*P, f*_out_), thus

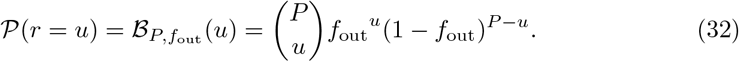

This describes the output unit usage more accurately than the assumption that each output unit is active exactly *f*_out_ · *P* times across a pattern set of size *P*, as assumed in the original paper on the Willshaw network [28].

The dendritic sum of the output unit *j* gets contributions only from connections that originate at one of the *M*_in_ input units that are active in the *k*-th pattern. Independently of their state (functional or silent), connections from inactive input units do not contribute to the net input to an output unit. We define *ρ*_*s*_(*u*) as the probability that a genuine-spurious connection connecting to output unit *j* is functional given that the output unit *j* is active *u* times in the pattern set. Similarly, the probability that a genuine-genuine connection is functional is called *ρ*_*g*_(*u*). The functionality of a connection implicitly takes into account that the connection is morphologically available. Recall that *ρ*_*l*_(*u*), for *l* ∈ {*s, g*}, depends on whether the respective output unit is genuine (*ρ*_*g*_(*u*)), i.e., active in the target output, or spurious (*ρ*_*s*_(*u*)), i.e., inactive in the target output. The derivation of *ρ*_*s*_(*u*) and *ρ*_*g*_(*u*) will be discussed in the following sections.Moreover, the probability that a spurious-spurious or a spurious-genuine connection is functional is less interesting because these connections do not contribute to the dendritic sums due to lack of input. However, these probabilities need to be considered implicitly in the following sections. to be able to determine *ρ*_*s*_(*u*) and *ρ*_*g*_(*u*). Further, the probability of a functional spurious-genuine connection, called *ρ*_*n*_(*u*), is important if the input pattern used for retrieval is noisy. This case will be discussed later.

For a particular output unit usage *u* and statistically independent output units, the dendritic sums 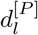 are binomially distributed with

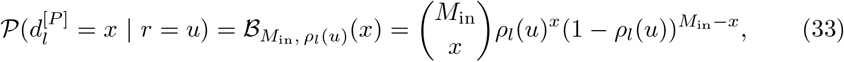

for *l* ∈ {*s, g*}. In the extreme case where all connections from input units that are active in the *k*-th pattern to an output unit are functional, its dendritic sum equals the number of active input units *M*_in_. Combining Eq. (32) and Eq. (33), we obtain

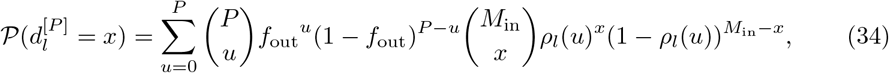

for *l* ∈ {*s, g*}, as the distribution of the dendritic sums 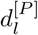. We abbreviate

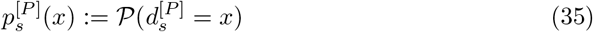

and

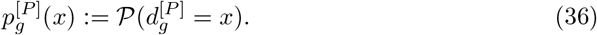

Before calculating the probability *ρ*_*l*_(*u*) of a single genuine-genuine or a single genuine-spurious connection being functional after storing a set of additional patterns in which the output unit was active *u* times, we review the learning rule from a probabilistic perspective and determine important fractions of connections.

### Hebbian and homeostatic update rules from a probabilistic perspective

We assume a morphological connectivity of *c*_*m*_ · *N*_in_ connections per output unit and start from a random choice of *c* · *N*_in_ functional connections targeting each output unit. In each learning step *k*, only connections targeting output units that are active in the current pattern, i.e., genuine-genuine and spurious-genuine connections, are updated. We further can distinguish between four types of connections: They can be either silent or functional before learning the *k*-th pattern and they can either originate from an active or an inactive input unit in the *k*-th pattern. Two types of connections are not updated: If the connection is already functional and the corresponding input unit is active (in addition to the corresponding output unit being active), the connection is protected and remains functional. If the connection is previously silent and the input unit is inactive, the connection always remains silent. The other two types can transition between the two states (functional/silent): Genuine-genuine connections (a fraction *f*_in_ of all connections to one output unit) of the *k*-th pattern that are previously silent (a fraction *c*_*m*_ − *c*) are made functional with a transition probability *η*, i.e., the fraction *f*_in_(*c*_*m*_ − *c*)*η* of all connections becomes functional in one learning step.

For each genuine output unit in the *k*-th pattern, the same number of connections that were made functional is also silenced such that there is a constant fraction *c* of functional connections per output unit after Hebbian learning and homeostatic normalization. Only spurious-genuine connections of the *k*-th pattern (fraction 1− *f*_in_) that are previously functional (fraction *c*) could be silenced due to the homeostasis mechanism. This is a total fraction *c*(1− *f*_in_) of all connections. The probability that such a connection is silenced depends on the number of additional connections that became functional due to the Hebbian mechanism in the same learning step. As discussed above, a fraction *η* · (*c*_*m*_ − *c*)*f*_in_ becomes functional in each step. The probability for a previously functional connection from an inactive input unit to be silenced due to homeostasis is

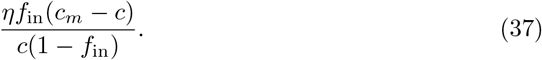

Normalization is possible in this way if this probability is less than or equal to one, which is always fulfilled if we assume that *c* ≥ *f*_in_ (regardless of the values of *η* and *c*_*m*_).

In summary, the numerator in Eq. 37 is the fraction of connections that are made functional in this step and the denominator is the fraction of connections available to be silenced; together, this is the *probability* to be newly silenced. Since the *fraction* of connections made functional, *ηf*_in_(*c*_*m*_ − *c*), equals the *fraction* of connections newly silenced, 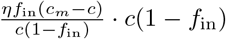, the overall fraction *c* of functional connections is maintained. Since patterns are uncorrelated, these probabilities remain the same for every pattern that is learned, and they will be used several times in the derivations in the following subsections.

### Probability of a functional connection to a spurious unit

First, we consider an output unit that is spurious for the *k*-th pattern. As we show below, the probability *ρ*_*s*_(*u*) for genuine-spurious connections to be functional given that the respective output unit was active *u* times across the patterns 1, …, *P* does not change with *u*. Due to the initialization of turning on *cN*_in_ connections per output unit (blue part in Fig 7A) and the fact that storing the *k*-th pattern does not affect the connections to spurious units, we have that *ρ*_*s*_(0) = *c* (orange part in Fig 7B). Only connections to genuine units in each pattern are turned on or off. Hence, we do not need to distinguish between genuine-spurious and spurious-spurious connections in the *k*-th pattern (as we will have to in the next section where we discuss the probability of a functional connection to a genuine unit). While storing additional patterns, connections targeting output units that are spurious with respect to the *k*-th pattern will be updated because they might not be spurious with respect to the other patterns. Since the *k*-th input pattern is uncorrelated with the other input patterns and since the homeostasis mechanism maintains the functional connectivity level *c* at all times, the probability that a connection to a spurious output unit (with respect to the *k*-th pattern) is on remains *c*.

**Fig 7.**
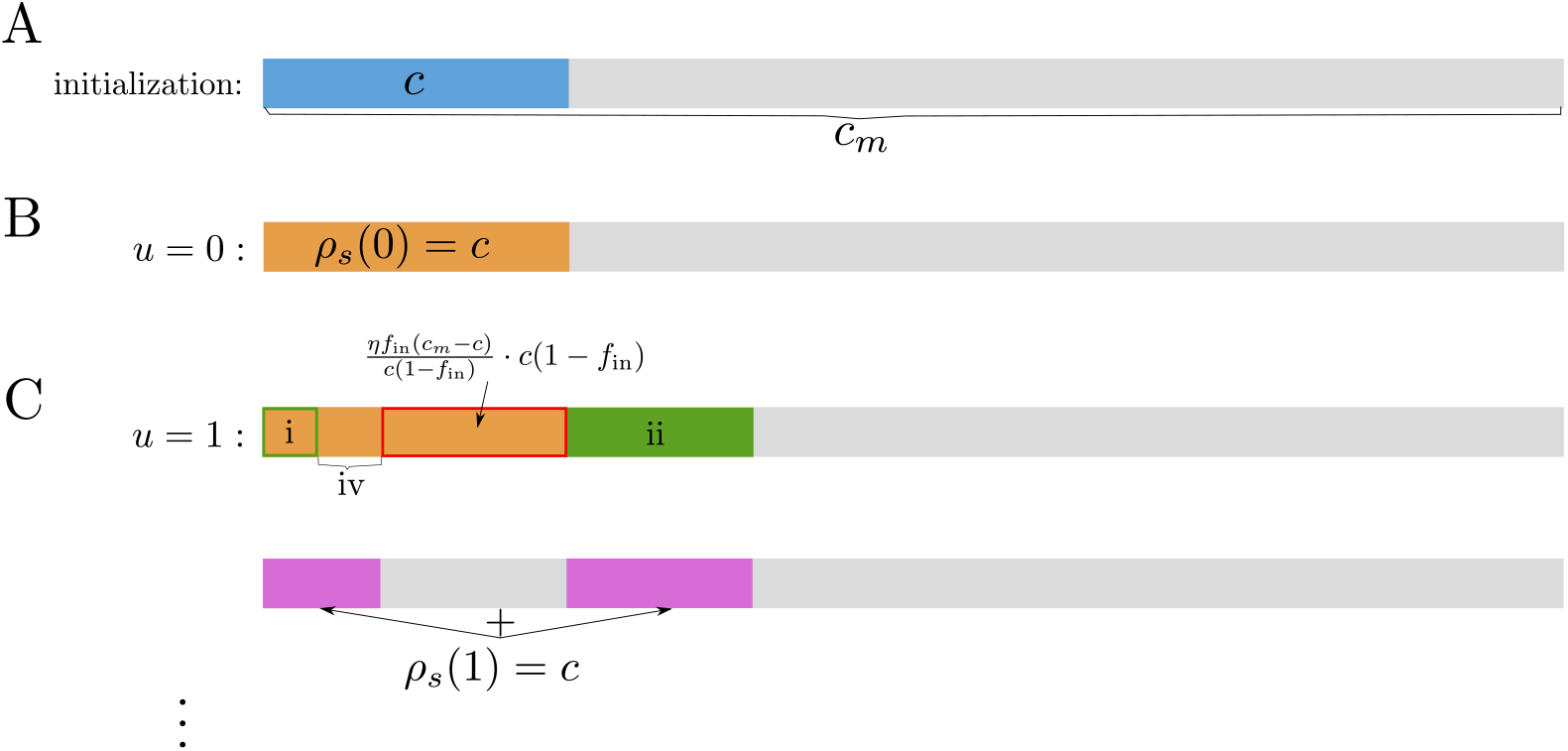
Illustration of the probability of a functional connection to a spurious unit. Fractions of functional connections targeting a spurious output unit if the output unit is active *u* times across the pattern set. Grey parts represent fractions of silent connections. Colored parts represent fractions of functional connections. (**A**) Configuration after initialization. (**B**) Configuration for *u* = 0. (**C**) Configuration for *u* = 1.

Let us explain this result in more detail: We assume that the spurious output unit that we are investigating is active for one additional pattern *k* + *κ* with *κ* ∈ ℕ^+^ fixed. The probability *ρ*_*s*_(1) of a connection to this unit to be functional after the update step can be decomposed using the law of total probability

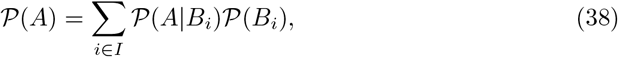

where {*B*_*i*_}_*i∈I*_ is a partition of the sample space. As conditions *B*_*i*_, we distinguish between the four possible combinations that arise from two initial states of the connection and two activity states of the input unit: either the connection is already functional before learning pattern *k* + *κ* (with probability *c*, orange part in Fig 7B) or it is previously silent (but morphologically available) (with probability *c*_*m*_ − *c*, grey part in Fig 7B); furthermore, the originating input unit is either active (with probability *f*_in_) or inactive (with probability 1 − *f*_in_) in pattern *k* + *κ*. These four conditions have the following probabilities:

i. 𝒫 (*‘connection previously functional and input unit active’*) = *cf*_in_
ii. 𝒫 (*‘connection previously silent and input unit active’*) = (*c*_*m*_ − *c*)*f*_in_
iii. 𝒫 (*‘connection previously silent and input unit inactive’*) = (*c*_*m*_ − *c*)(1 − *f*_in_)
iv. 𝒫 (*‘connection previously functional and input unit inactive’*) = *c*(1 − *f*_in_)

Remember that the output unit that we are investigating is spurious with respect to the *k*-th pattern but it is active in pattern *k* + *κ*. For each of the four cases above, we derive the conditional probability of a functional connection based on the update rule that consists of Hebbian and homeostatic learning:

i. If the connection is already functional and the corresponding input unit is active, the connection remains functional: 𝒫 (*‘connection functional’* | *‘connection previously functional and input unit active’*) = 1 (yielding the green frame in Figure 7C)
ii. If the connection is silent and the corresponding input unit is active, the connection is made functional with transition probability *η*: 𝒫 (*‘connection functional’* | *‘connection previously silent and input unit active’*) = *η* (yielding the green area in Figure 7C)
iii. If the connection is silent and the input unit is inactive, the connection remains silent: 𝒫 (*‘connection functional’*|*’connection previously silent and input unit inactive’*) = 0
iv. If the connection is functional and the input unit is inactive, the connection could be silenced due to the homeostasis mechanism (shown by the red frame in Figure 7C) or it could remain functional. As derived in Eq. (37), the probability that it is silenced is *ηf*_in_(*c*_*m*_ − *c*)*/* (*c*(1− *f*_in_)) and the probability that it remains functional is thus 𝒫 (*‘connection functional’* |*’connection previously functional and input unit inactive’*) 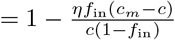 (represented by the orange area without the red frame and without the green frame in Figure 7C).

In total, according to Eq. (38), the probability of the connection to be functional can be expressed as

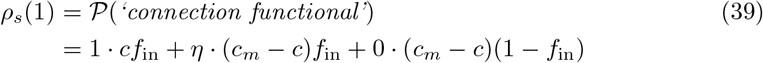

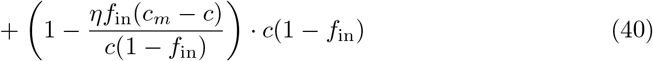

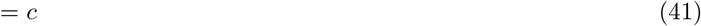

which is represented by the purple parts in Fig 7C.

The preservation of *ρ*_*s*_(*k*) relies on the statistical identity of units in the distribution of patterns. Specifically, every additional pattern in which a particular spurious output unit is active manipulates the connections to the spurious output unit in the same way. It follows that

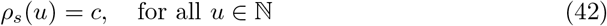

and, in the following, we denote the distribution of the dendritic sums of spurious units for any *P* as

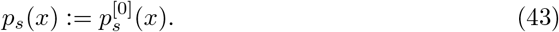

### Probability of a functional connection to a genuine unit

We have seen that the probability *ρ*_*s*_ that a connection to a spurious unit is functional does not change with the output unit usage *u*. The probability *ρ*_*g*_ that a connection to a genuine unit is functional, however, does strongly depend on *u*. In the following, we first derive *ρ*_*g*_(0) and *ρ*_*g*_(1) and then generalize this derivation recursively to *ρ*_*g*_(*u*) for *u >* 1.

### Derivation of *ρ*_*g*_(0) and *ρ*_*g*_(1)

We aim at calculating the probability *ρ*_*g*_(*u*) that a genuine-genuine connection of the *k*-th pattern (Fig 8 left, labeled by *f*_in_) is functional under the condition that the respective output unit is active *u* times in the subsequent patterns *k* + 1, …, *k* + *P*. Here, we compute *ρ*_*g*_(*u*) for *u* = 0, 1. For large numbers of such connections, i.e. for *f*_in_*N*_in_ · *f*_out_*N*_out_ ≫ 1, the probabilities *ρ*_*g*_(0) and *ρ*_*g*_(1) can, by the law of large numbers, be interpreted as fractions. Similarly to describing *ρ*_*g*_(*u*), we can describe the fraction of functional connections originating from input units that are inactive in the *k*-th pattern and targeting one specific genuine output unit, which are called spurious-genuine connections (Fig 8 right, labelled by 1 − *f*_in_). For the output unit being active never or once, we name these fractions *ρ*_*n*_(0) and *ρ*_*n*_(1), respectively. These probabilities of functional spurious-genuine connections are needed for deriving *ρ*_*g*_ but they do not explicitly contribute to the dendritic sum of the output unit because they do not receive any input.

**Fig 8.**
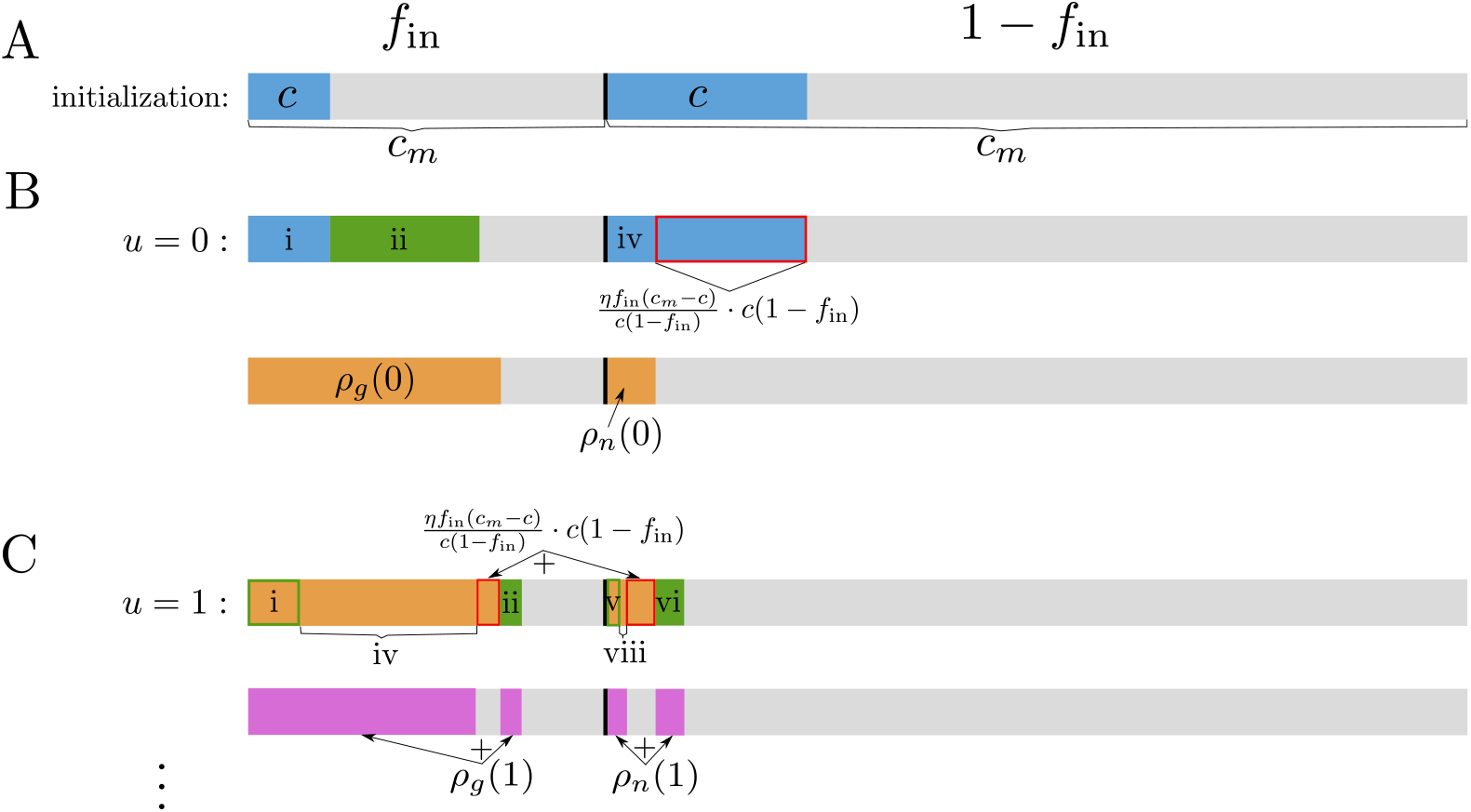
Illustration of the probability of a functional connection to a genuine unit. Fractions of functional connections targeting a genuine output unit if the output unit is active *u* times across the pattern set. Left (below “*f*_in_”): fraction *f*_in_ of connections that originate from an input unit that is active in the *k*-th pattern and will thus contribute to the dendritic sum of the output unit when the *k*-th input pattern is applied — the genuine-genuine connections; right (below “1 − *f*_in_”): fraction 1− *f*_in_ of connections that originate from an input unit that is inactive in the *k*-th pattern — the spurious-genuine connections. Grey areas represent fractions of silent connections. Colored areas represent fractions of functional connections. (**A**) Configuration after initialization. (**B**) Configuration for *u* = 0. (**C**) Configuration for *u* = 1.

As discussed in the previous section, connections targeting a spurious output unit are not modified during learning the *k*-th pattern, and the probability of a functional connection to a spurious unit after learning the *k*-th pattern is the same as the probability at the initialization. Since there is no correlation between input patterns, the probability of functionality is identical for genuine-spurious and spurious-spurious connections. In contrast, the probability of a functional connection to a genuine output unit after learning the *k*-th pattern (i.e., for *u* = 0) depends on whether the connection originates at an input unit that is active in the *k*-th pattern or not. Thus, we separately examine the probability of a functional genuine-genuine connection, *ρ*_*g*_, and the probability of a spurious-genuine connection, *ρ*_*n*_. Note that the two probabilities *ρ*_*g*_ and *ρ*_*n*_, when weighted by the probabilities 𝒫 (‘input unit active’) = *f*_in_ and 𝒫 (‘input unit inactive’) = 1 −*f*_in_, respectively, should, nevertheless, sum up to the functional connectivity *c*:

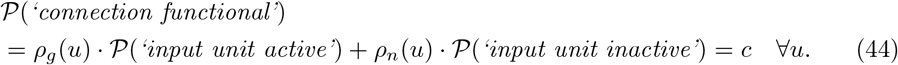

### Output unit never active after learning the *k*-th pattern (u = 0)

For *ρ*_*g*_(0) and *ρ*_*n*_(0), only the storage of the *k*-th pattern plays a role. Starting from a random fraction *c* (thus a total number of *cN*_in_) of functional connections targeting each output unit (blue parts in Fig 8A), we first consider connections only from an active input unit (left part, titled *f*_in_, in Fig 8). We can split up the probability of a genuine-genuine connection to become functional into the probabilities of two mutually exclusive conditions and their corresponding conditional probabilities (see Eq. (38)). We have the two conditions

i. 𝒫 (*‘g-g connection previously functional’*) = *c*
ii. 𝒫 (*‘g-g connection previously silent’*) = *c*_*m*_ − *c*, for each of which we compute the conditional probability of a functional genuine-genuine connection:
  i. If the connection is already functional and the corresponding input unit is active, the connection is protected and remains functional: 𝒫 (*‘g-g connection functional’* | *‘g-g connection previously functional’*) = 1 (yielding the blue part in Figure 8B left)
  ii. If the connection is silent and the corresponding input unit is active, the connection is made functional with transition probability *η*: 𝒫 (*‘g-g connection functional’* | *‘g-g connection previously silent’*) = *η* (yielding the green part in Figure 8B) According to the law of total probability, the probability of a functional genuine-genuine connection is thus

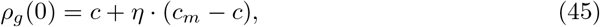

which is depicted by the left orange part in Fig 8B. Analogously, the probability of a functional spurious-genuine connection is derived (right part, titled 1 − *f*_in_, in Fig 8). Given that the corresponding input unit is inactive, the two conditions have the probabilities
  iii. 𝒫 (*‘s-g connection previously silent’*) = *c*_*m*_ − *c*
  iv. 𝒫 (*‘s-g connection previously functional’*) = *c*,
  and the corresponding conditional probabilities are the following:
iii. If the connection is silent and the input unit is inactive, the connection remains silent: 𝒫 (*‘s-g connection functional’* | *‘s-g connection previously silent’*) = 0
iv. If the connection is functional and the input unit is inactive, the connection could either be silenced due to the homeostasis mechanism (represented by the red frame in Figure 8B) or it could remain functional. As before, the probability for case (iv) connections to remain functional is 𝒫 (*‘s-g connection functional’* | *‘s-g connection previously functional’*)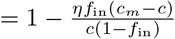 (yielding the blue part without the red frame in Figure 8B right).

Then, the total probability of a functional spurious-genuine connection is

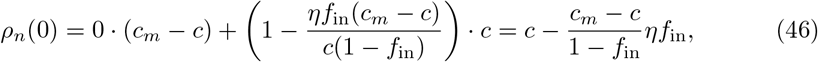

which is represented by the right orange part in Fig 8B. By using Eqs. (45) and (46) in Eq.(44), we see that the total probability of a functional connection to a genuine unit after learning the *k*-th pattern is again

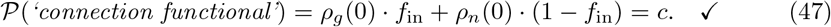

### Output unit active once after learning the *k*-th pattern (u = 1)

Next, we derive *ρ*_*g*_(1) and *ρ*_*n*_(1), which correspond to the probabilities that a genuine-genuine or a spurious-genuine connection (with respect to the *k*-th pattern) is functional under the condition that the respective output unit is active exactly once among all subsequent patterns. The derivation of *ρ*_*g*_(1) and *ρ*_*n*_(1) is more involved because functional connections that originated from an active input unit in the *k*-th pattern (genuine-genuine connections) can be silenced while storing additional patterns, and in turn silent connections originating from inactive input units in the *k*-th pattern (spurious-genuine connections) can be made functional. We assume that the genuine output unit that we are investigating is active only for one pattern *k* + *κ* with *κ* ∈ℕ^+^ fixed.

Let us start with the derivation of *ρ*_*g*_(1). The fraction of genuine-genuine connections being functional after storing pattern *k* + *κ* (in addition to pattern *k*) can be split into four conditions depending on the previous state of the connection (i.e., depending on *ρ*_*g*_(0)) and on the activity state of the corresponding input unit in pattern *k* + *κ*. The four conditions have the probabilities

i. 𝒫 (*‘g-g connection previously functional and input unit active’*) = *ρ*_*g*_(0)*f*_in_
ii. 𝒫 (*‘g-g connection previously silent and input unit active’*) = (*c*_*m*_ − *ρ*_*g*_(0))*f*_in_
iii. 𝒫 (*‘g-g connection previously silent and input unit inactive’*) = (*c*_*m*_ − *ρ*_*g*_(0))(1 − *f*_in_)
iv. 𝒫 (*‘g-g connection previously functional and input unit inactive’*) = *ρ*_*g*_(0)(1 − *f*_in_) and the corresponding conditional probabilities are
  i. If the connection is already functional before storing pattern *k* + *κ* and the corresponding input unit is active, the connection is protected and remains functional: 𝒫 (*‘g-g connection functional’* | *‘g-g connection previously functional and input unit active’*) = 1 (represented by the green frame in Figure 8C left)
  ii. If the connection is silent and the corresponding input unit is active in pattern *k* + *κ*, the connection is made functional with transition probability *η*: 𝒫 (*‘g-g connection functional’* | *‘g-g connection previously silent and input unit active’*) = *η* (yielding the green area in Figure 8C left)
  iii. If the connection is silent and the input unit is inactive, the connection remains silent: 𝒫 (*‘g-g connection functional’* | *‘g-g connection previously silent and input unit inactive’*) = 0
  iv. If the connection is functional and the input unit is inactive, the connection remains functional with probability 𝒫 (*‘g-g syn. functional’* | *‘g-g syn. previously functional and input unit inactive’*)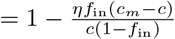 (shown by the orange area without the red frame and without the green frame in Figure 8C left). Using the law of total probability, we thus have

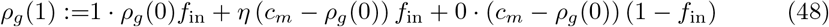

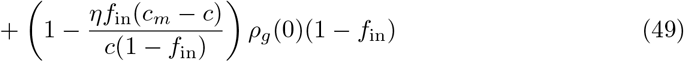

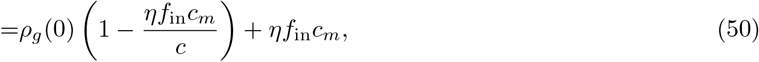

shown by the purple parts in Figure 8C left. Analogously, the probability *ρ*_*n*_(1) of spurious-genuine connections being functional after storing pattern *k* + *κ* consists of the four conditions
  v. 𝒫 (*‘s-g connection previously functional and input unit active’*) = *ρ*_*n*_(0)*f*_in_
  vi. 𝒫 (*‘s-g connection previously silent and input unit active’*) = (*c*_*m*_ − *ρ*_*n*_(0))*f*_in_
  vii. 𝒫 (*‘s-g connection previously silent and input unit inactive’*) = (*c*_*m*_ − *ρ*_*n*_(0))(1 − *f*_in_)
  viii. 𝒫 (*‘s-g connection previously functional and input unit inactive’*) = *ρ*_*n*_(0)(1 − *f*_in_) and the four corresponding conditional probabilities
v. 𝒫 (*‘s-g connection functional’*|*’s-g connection previously functional and input unit active’*) = 1 (represented by the green frame in Figure 8C right)
vi. 𝒫 (*‘s-g connection functional’* | *‘s-g connection previously silent and input unit active’*) = *η* (yielding the green area in Figure 8C right)
vii. 𝒫 (*‘s-g connection functional’*|*’s-g connection previously silent and input unit inactive’*) = 0
viii. P(*‘s-g syn. functional’* | *‘s-g syn. previously functional and input unit inactive’*)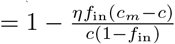 (represented by the orange area without the red frame and without the green frame in Figure 8C right)

According to the law of total probability, the probability of a functional spurious-genuine connection amounts to

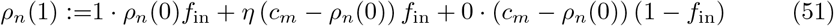

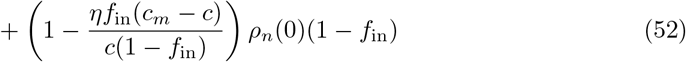

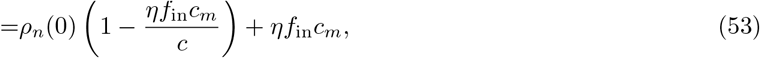

which corresponds to the purple parts in Fig 8C right.

### Generalization for any output unit usage *u >* 0

The probability of a functional connection, if the corresponding output unit is active *u* times across the whole pattern set, can be computed recursively. The conditional probabilities remain the same in every step, while the four conditions always depend on the probabilities *ρ*_*g*_(*u* − 1) and *ρ*_*n*_(*u* − 1) of the previous step. We thus obtain *ρ*_*g*_(2) from *ρ*_*g*_(1) the same way we obtained *ρ*_*g*_(1) from *ρ*_*g*_(0),

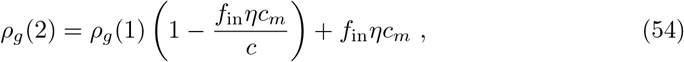

and in general we get

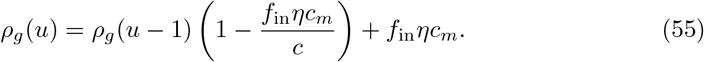

*ρ*_*n*_(*u*) can be derived in the same way and we have

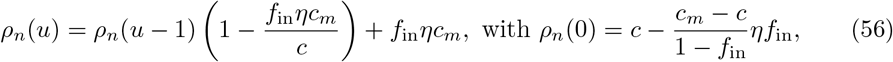

where the only difference from *ρ*_*g*_(*u*) is the initial value *ρ*_*n*_(0) ≠ *ρ*_*g*_(0) (see Eqs. (45) and (46)).

From the recursive description of *ρ*_*g*_(*u*) (Eqs. (45) and (55)), we can derive an explicit description

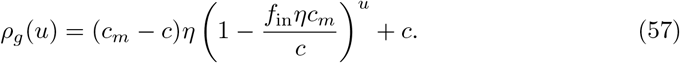

Examples of the probability for a genuine-genuine connection to be functional *ρ*_*g*_(*u*) as a function of the output unit usage *u* are shown in Fig 9. Analogously, the explicit description of *ρ*_*n*_ is

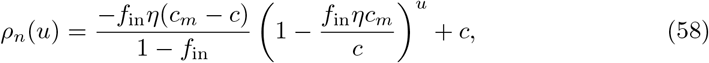

and it will come up again later when we discuss retrieval with noisy cues.

**Fig 9.**
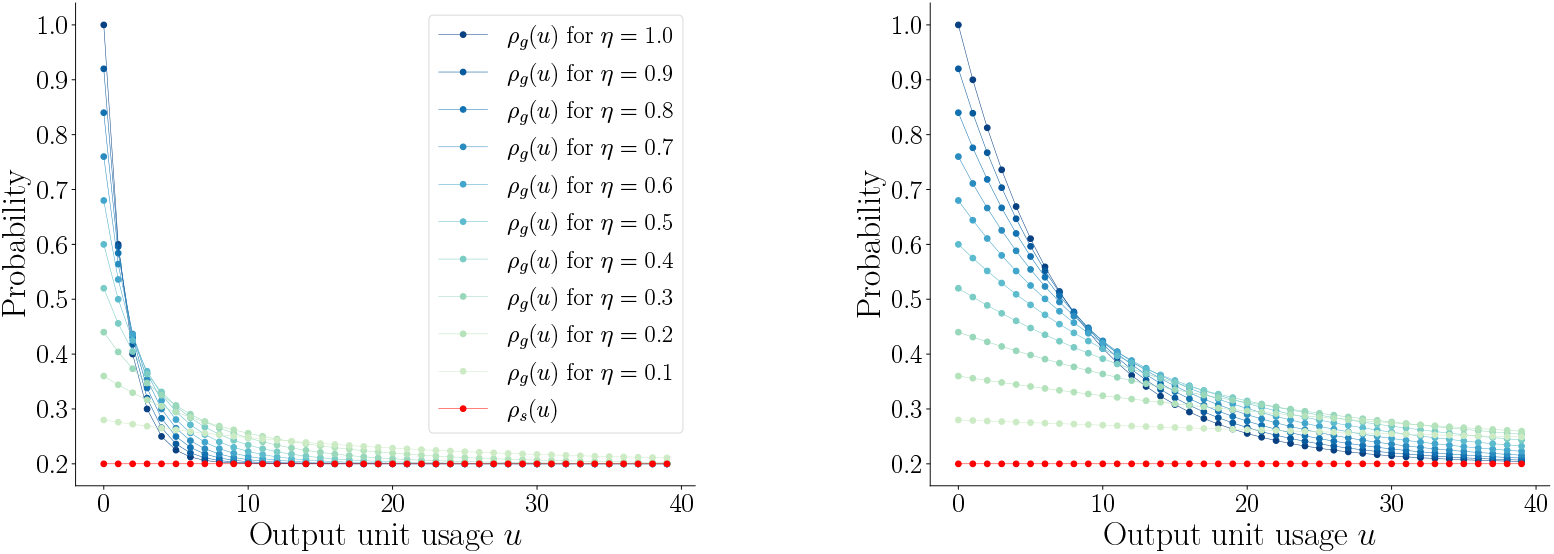
Probabilities of functional connections *ρ*_*g*_(*u*) and *ρ*_*s*_(*u*). Probability *ρ*_*l*_(*u*), *l* ∈ {*s, g*}, of a connection to be functional depending on output unit usage *u* for various transition probabilities *η*. Blue dots: genuine-genuine connections (Eq. (57)); red dots: genuine-spurious connections (Eq. (42)). Left: *f*_in_ = 0.1, right: *f*_in_ = 0.025. Probability *ρ*_*s*_ of genuine-spurious connections to be functional is constant as a function of *u*. Probability *ρ*_*g*_ of genuine-genuine connections to be functional decreases exponentially with *u*. (Other parameters: *c* = 0.2, *c*_*m*_ = 1.)

To summarize, we have now fully characterized the distributions of dendritic sums for spurious and for genuine output units. We insert the probability of a functional connection to a spurious unit *ρ*_*s*_(*u*) = *c* into Eq. (34) and obtain the probability mass function (PMF) of the dendritic sum of a spurious output unit

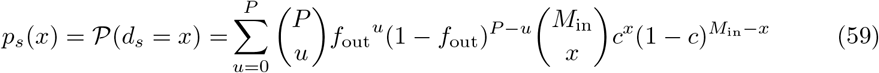

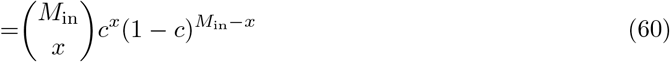

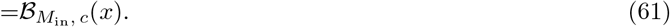

Since *ρ*_*s*_(*u*) is constant as a function of the output unit usage *u*, the distribution of dendritic sums for spurious output units *p*_*s*_ simplifies to a single binomial distribution ℬ_*n, p*_ with parameters *n* = *M*_in_ and *p* = *c*, and *p*_*s*_ does not depend on the number of subsequently learned patterns *P*.

Analogously, we insert 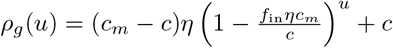 into Eq. (34) and obtain the PMF of the dendritic sum of a genuine output unit

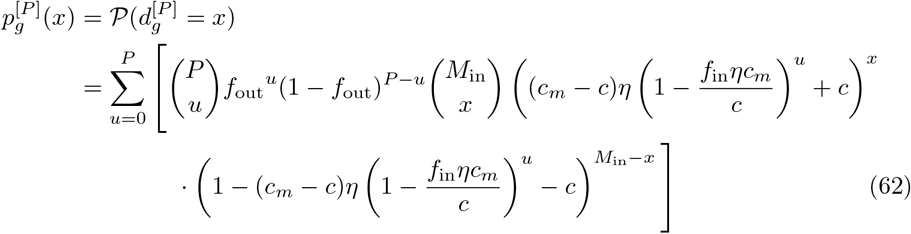

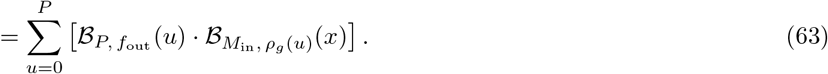

Thus, 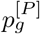 depends on the number of subsequent patterns *P* (in contrast to *p*_s_). It is not a single binomial but a linear combination of binomial PMFs 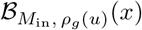, for *u* ∈ {0, …, *P*}, see Fig 10. This multimodality is analyzed in more detail in S1 Appendix. The relation between *p*_*s*_ and 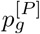 and their dependence on the activation ratios *f*_in_ and *f*_out_ is discussed in the Results. S5 Fig shows a comparison of Eqs. (61) and (63) to distributions of dendritic sums obtained from numerical simulations.

**Fig 10.**
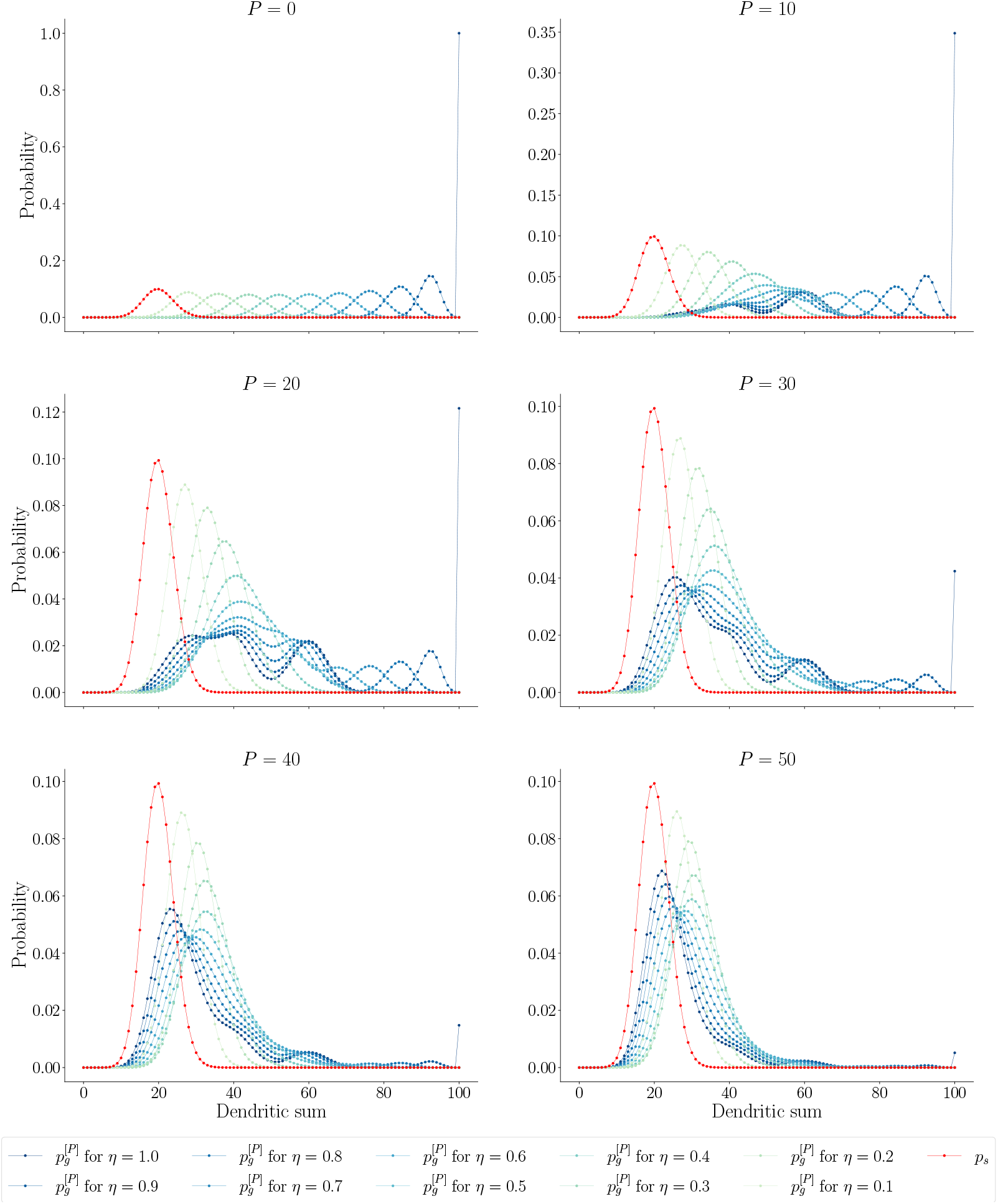
Distributions of dendritic sums. Analytical probability mass functions of dendritic sums after storing *P* = 0, 10, 20, 30, 40, 50 patterns (Eqs. (61) and (63)). Spurious units in red; genuine units in blue shades, for several transition probabilities *η*. With an increasing number of subsequent patterns *P*, the distributions of the dendritic sums of genuine units approach the distribution of the dendritic sums of spurious units. The genuine distributions are multimodal, in particular for large transition probabilities *η* and small numbers of patterns *P >* 0. Other parameters: *N*_in_ = *N*_out_ = 1000, *f*_in_ = *f*_out_ = 0.1, *c* = 0.2, *c*_*m*_ = 1.

Let us finally discuss some limiting cases. We have

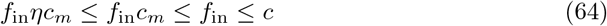

because 0 *< η* ≤ 1, 0 *< c*_*m*_ ≤ 1, 0 *< f*_in_, and *f*_in_ ≤ *c*. Note that the constraint *f*_in_ ≤ *c* stems from the normalization condition as discussed previously. It thus follows that we either have

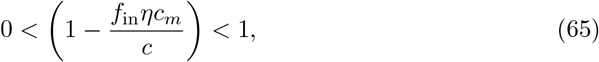

which implies

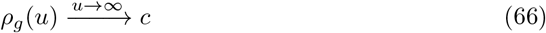

or, for *c*_*m*_ = 1, η = 1, and *f*in = *c*,

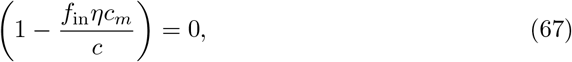

which implies

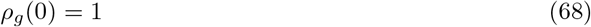

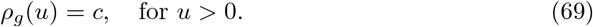

In the second case, the part of the memory stored in the connections targeting one output unit is completely lost as soon as this output unit is active one additional time for another pattern.

### Noisy input patterns during retrieval

So far we always assumed that during recall the original input pattern is shown to the network. Due to the inherent presence of noise in biological systems, it makes sense to assume that the input pattern that is presented to the network during retrieval is not exactly the same as the input pattern used for training. In the following, we discuss the consequences on the previous derivations when a noisy cue is presented to the network by putting noise onto the input pattern. To be able to compare the results in a fair fashion, the input activation ratio *f*_in_ should not be changed, hence a fraction *ε* of the input units that are active in the original pattern (genuine input units) is deactivated while the same absolute number of input units that are inactive in the original pattern (spurious input units) is activated. Thus there is a number *m*_*g*_ = *M*_in_ · (1 − *ε*) of original input units (genuine input units) and a number *m*_*s*_ = *M*_in_ · *ε* of wrong input units (spurious input units) active, in total we have *M*_in_ = *m*_*g*_ + *m*_*s*_ active units [69]. The parameter *ε* is restricted to values that yield *m*_*g*_, *m*_*s*_ ∈ ℕ. Note that, in the numerical simulations, we set each of the genuine input units to zero with probability *ε*, count how many have changed and flip the same amount of spurious input units to one. This means *m*_*g*_ and *m*_*s*_ are not fixed exact numbers but follow distributions. Only *on average, M*_in_*ε* genuine units are deactivated. The additional variability coming from deactivating each genuine input unit with probability *ε* would make the analytical description more involved while hardly affecting the expected result. It is therefore neglected in the theory derived in this section. Instead, we assume that, for generating a noisy cue, exactly *M*_in_*ε* of the genuine input units are deactivated and *M*_in_*ε* of the spurious input units are activated. Note that, nevertheless, in both the numerical simulations and the theory, it is always ensured that the absolute number of active units *M*_in_ in the noisy input pattern that is presented to the network does not change.

Noisy cues alter the distribution of the dendritic sums of genuine output units but not of spurious output units. The probability of a functional genuine-genuine connection is the same as before, *ρ*_*g*_. The probability of a functional spurious-genuine connection has already been introduced as *ρ*_*n*_(*u*) (Eq. (58)). Analogously to *ρ*_*g*_(*u*) in Eq. (55) and Eq. (57), *ρ*_*n*_(*u*) can be expressed recursively as

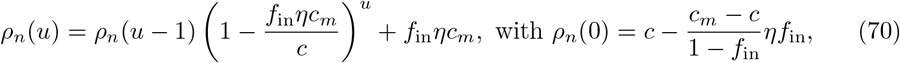

or explicitly as

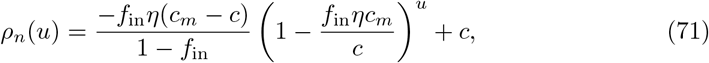

and is shown in Fig 11. Just like *ρ*_*g*_(*u*), *ρ*_*n*_(*u*) depends on the number of times *u* an output unit has been active across the pattern set. The probability of a functional spurious-genuine connection *ρ*_*n*_ is lower than the probability of a functional genuine-genuine connection *ρ*_*g*_ (and even lower than the functional connectivity *c*, see Figs 11 and 9).

**Fig 11.**
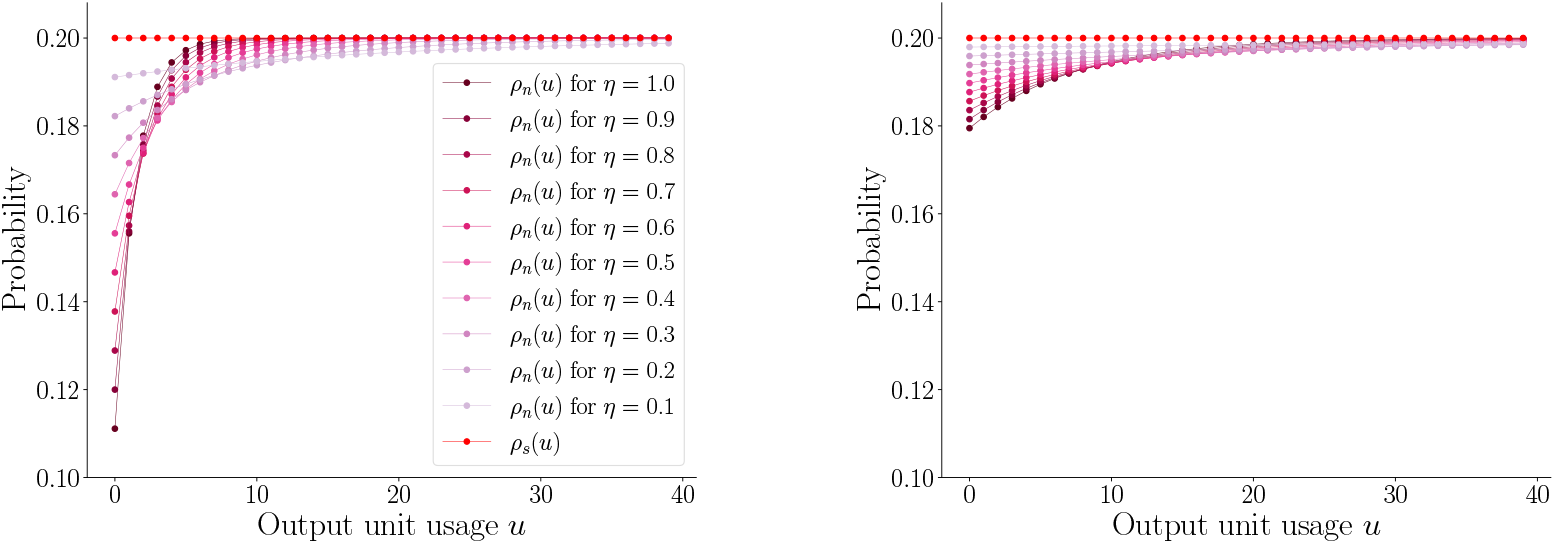
Probabilities of functional connections *ρ*_*n*_(*u*) and *ρ*_*s*_(*u*). Probability *ρ*_*l*_(*u*), *l* ∈ *n, s*, of a connection to be functional depending on output unit usage *u*. Pink shades: spurious-genuine connections (*ρ*_*n*_(*u*), Eq. (71)); red: connections to spurious units (*ρ*_*s*_(*u*), Eq. (42)). Left: *f*_in_ = 0.1, right: *f*_in_ = 0.025. Probability *ρ*_*s*_ of genuine-spurious connections to be functional is constant as a function of *u*. Probability *ρ*_*n*_ of spurious-genuine connections to be functional increases exponentially with *u*. (Other parameters: *c* = 0.2, *c*_*m*_ = 1.)

For a particular output unit usage *u*, the dendritic sums 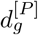 are distributed following

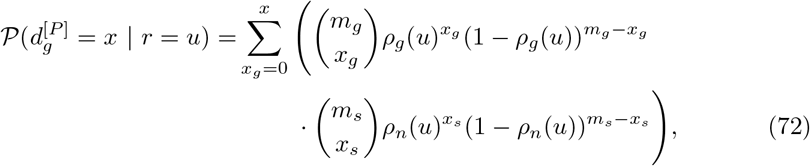

where *x*_*s*_ = *x* − *x*_*g*_. In total, the probability of the dendritic sum 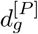 of a genuine output unit to have a particular value *x* can be described by

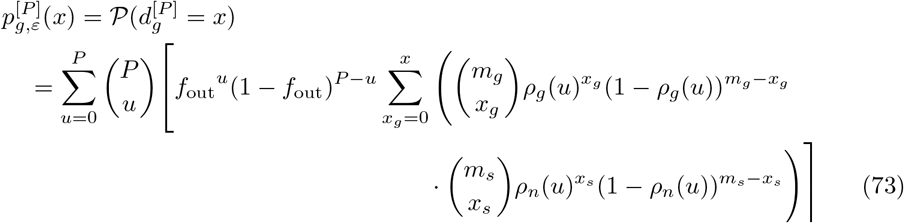

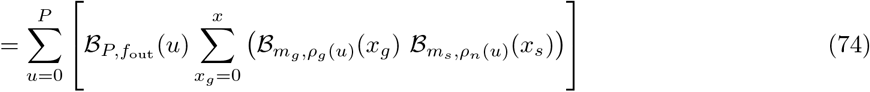

[69]. Compared to the distribution of the dendritic sums for the noise-less input pattern, this distribution is shifted to the left. Thus, it is closer to the distribution of the dendritic sums of the spurious units, the Hamming distance between the target output and the calculated output pattern is larger, and the capacity thus decreases (compare Fig 12 and S3 Fig to Fig 10).

**Fig 12.**
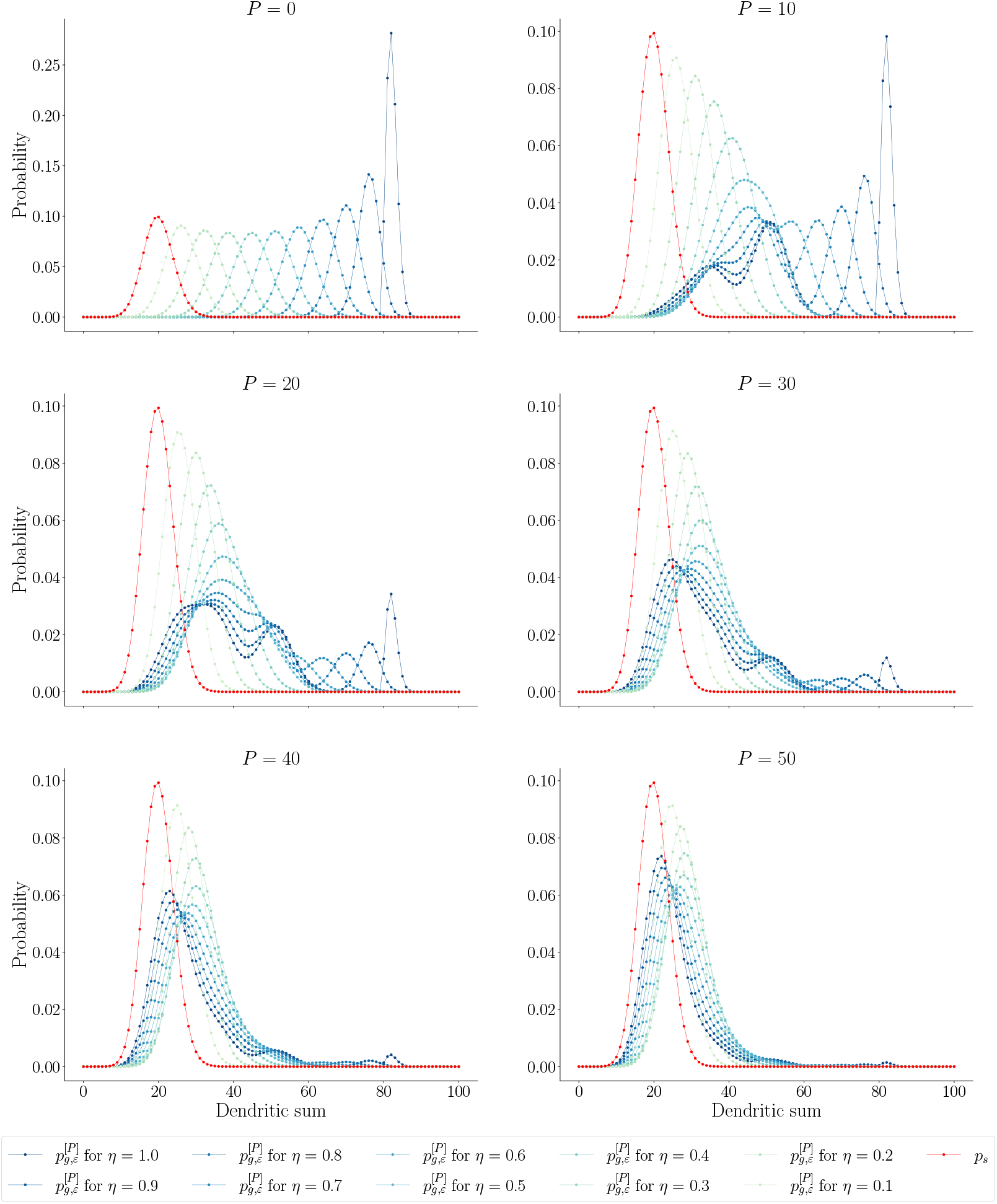
Distributions of dendritic sums with noise. Analytical probability mass functions of dendritic sums with noise on the input patterns during retrieval after storing *P* = 0, 10, 20, 30, 40, 50 patterns (Eq. (73) with (57) and (58); Eq. (34) with (42)). Spurious units in red; genuine units in blue shades, for several transition probabilities *η*. With an increasing number of subsequent patterns *P*, the distributions of the dendritic sums of genuine units approach the distribution of the dendritic sums of spurious units. Compared to the noise-less case (see Fig 10), the genuine distributions are closer to the spurious distribution. Other parameters: *N*_in_ = 1000, *f*_in_ = 0.1, *N*_out_ = 1000, *f*_out_ = 0.1, *c* = 0.2, *c*_*m*_ = 1, *ε* = 0.2. The same figure but with *ε* = 0.4 can be found in S3 Fig.

### Analytical calculation of the capacity of the network

In what follows, our aim is to analytically quantify the capacity of the network. The capacity is defined as the number of subsequent patterns *P* for which the signal quality has declined to the retrieval threshold. Therefore, we first discuss the signal quality as a function of *P*. Before we enter the details, we briefly recap basic quantities: The difference between a calculated output pattern **ŷ**^[*k*]^(*P*) given a particular input pattern **x**^[*k*]^ after learning *P* subsequent pattern pairs and the corresponding target output pattern **y**^[*k*]^ is measured in terms of the Hamming distance between the two, denoted by *H* **y**^[*k*]^, (**ŷ**^[*k*]^(*P*)). This Hamming distance is always averaged across a large number of patterns, and it lies between 0 and the average Hamming distance between two random patterns of length *N*_out_ and activation ratio *f*_out_, which is

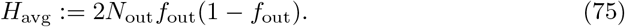

We define the memory signal quality *S*_*P*_ at *P* patterns as the difference between *H*_avg_ and the Hamming distance between the target output and the calculated output *H* (**y**^[*k*]^, **ŷ**^[*k*]^(*P*)) :

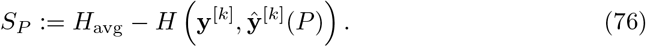

The signal quality can be expressed as a fraction *s*_*P*_ of *H*_avg_

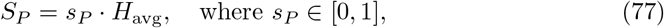

see Fig 13, green line. We call *s*_*P*_ the normalized signal quality. With *s*_*P*_, the Hamming distance reads as

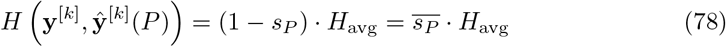

**Fig 13.**
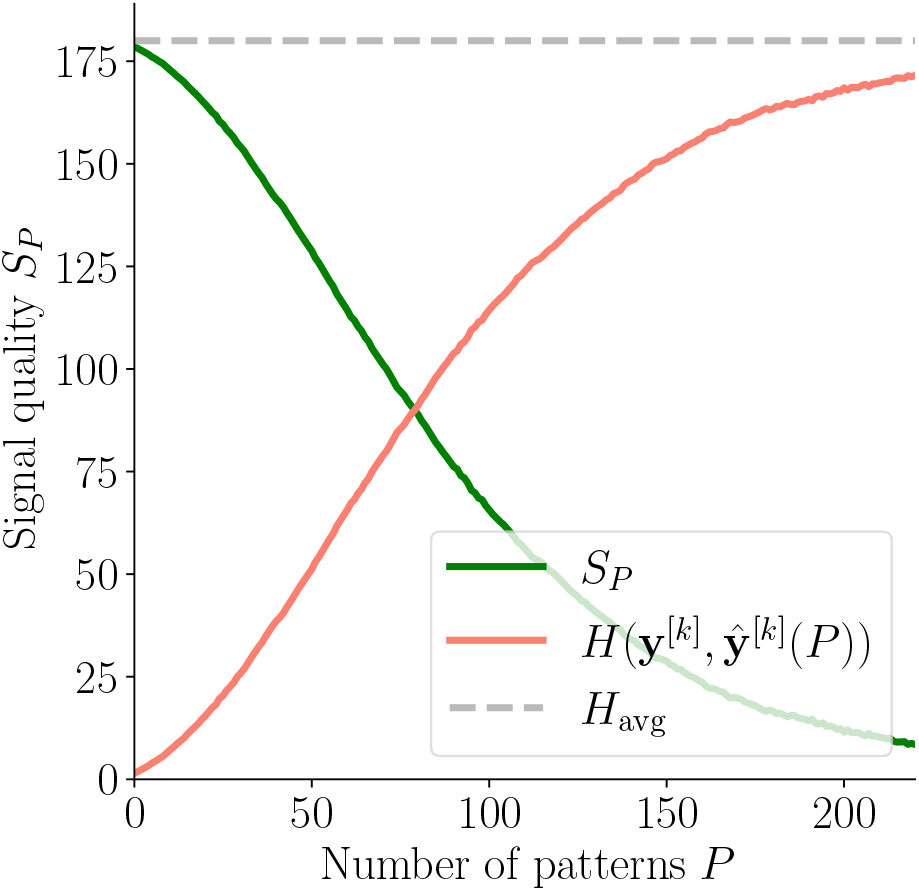
Hamming distance and signal quality. The Hamming distance *H* (**y**^[*k*]^, **ŷ**^[*k*]^(*P*)) between the target and the calculated output pattern (salmon line) increases with an increasing number of patterns *P* and approaches *H*_avg_ (grey line) for *P* → ∞. The signal quality *S*_*P*_ (green line) decreases with increasing *P* and approaches 0.

(see Fig 13, salmon line), where we introduce the additional symbol

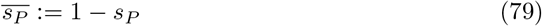

out of convenience for the derivations in this section.

### Analytical Hamming distance

In this section, we derive the expected Hamming distance 𝔼(*H* (**y**^[*k*]^, **ŷ**^[*k*]^(*P*))) between the target output **y**^[*k*]^ and the output **ŷ**^[*k*]^(*P*) calculated after learning *P* additional patterns.

### The activation threshold and balancing tails of the distributions

Since this Hamming distance depends on the activation threshold 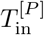, we first discuss the choice of 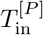, which is self-consistently defined such that the correct number of output units, *M*_out_, is activated:

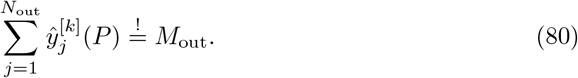

The activation of output unit *j* is calculated as

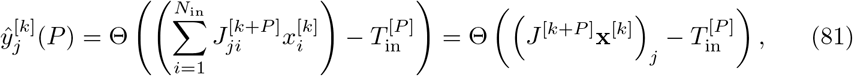

where 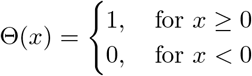 is the Heaviside step function. The number of active output units is hence

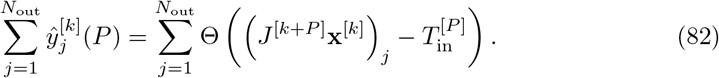

Since the weight matrix *J* ^[*k*+*P*]^ is updated in every learning step and hence depends on the number of subsequently learned patterns *P*, the activation threshold 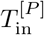 also depends on *P*. It thus has to be newly determined in every update step in order to obtain the given number of active output units *M*_out_.

The active output units can either be genuine, hence active in the target output pattern, or spurious, hence inactive in the target output pattern. The dendritic sum

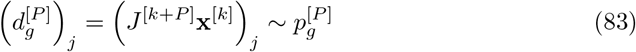

of a genuine output unit *j* follows the genuine distribution 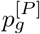 (Eq. (63)). A genuine output unit is activated if its dendritic sum is at least as large as the activation threshold 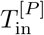, hence if

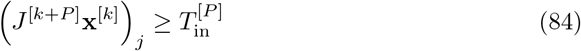

(blue area in Fig 14A). There is a total amount of *M*_out_ genuine output units. Thus, if we assume *M*_out_ ≫ 1, the number of genuine output units that are activated for a given threshold 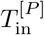 can be approximated by using the law of large numbers as

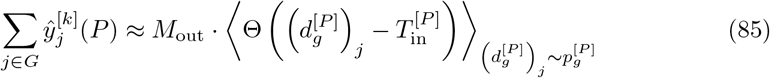

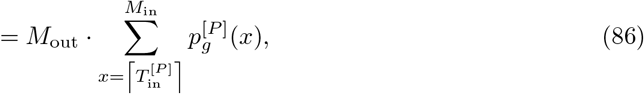

where *G* is the set of indices of genuine output units. Analogously, assuming *N*_out_ −*M*_out_ ≫1, the number of spurious units that are activated can be approximated as

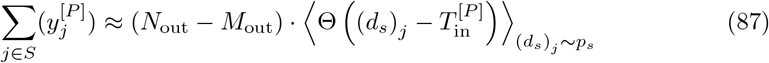

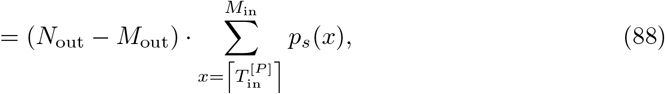

where (*d*_*s*_)_*j*_ denotes the dendritic sum of the *j*-th output unit, which is spurious, and *S* is the set of indices of spurious output units (cf. red area in Fig 14A). In order to activate the right amount of output units, the activation threshold 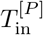 thus has to be chosen such that

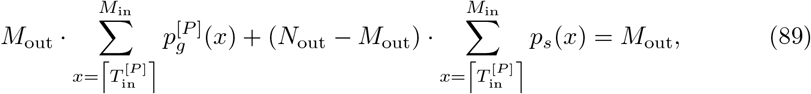

see Fig 14B. By basic computations, we have that

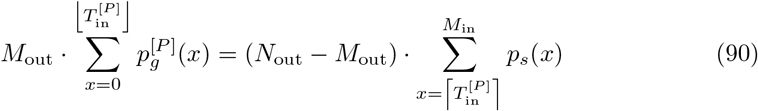

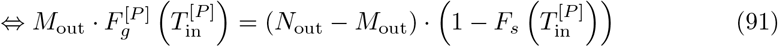

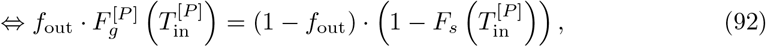

where 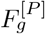 and *F*_*s*_ are the cumulative distribution functions (CDF) of 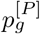 and *p*_*s*_, respectively (Fig 14C). We call the last equation the Balance Equation because it balances the false positive and the false negative errors.

**Fig 14.**
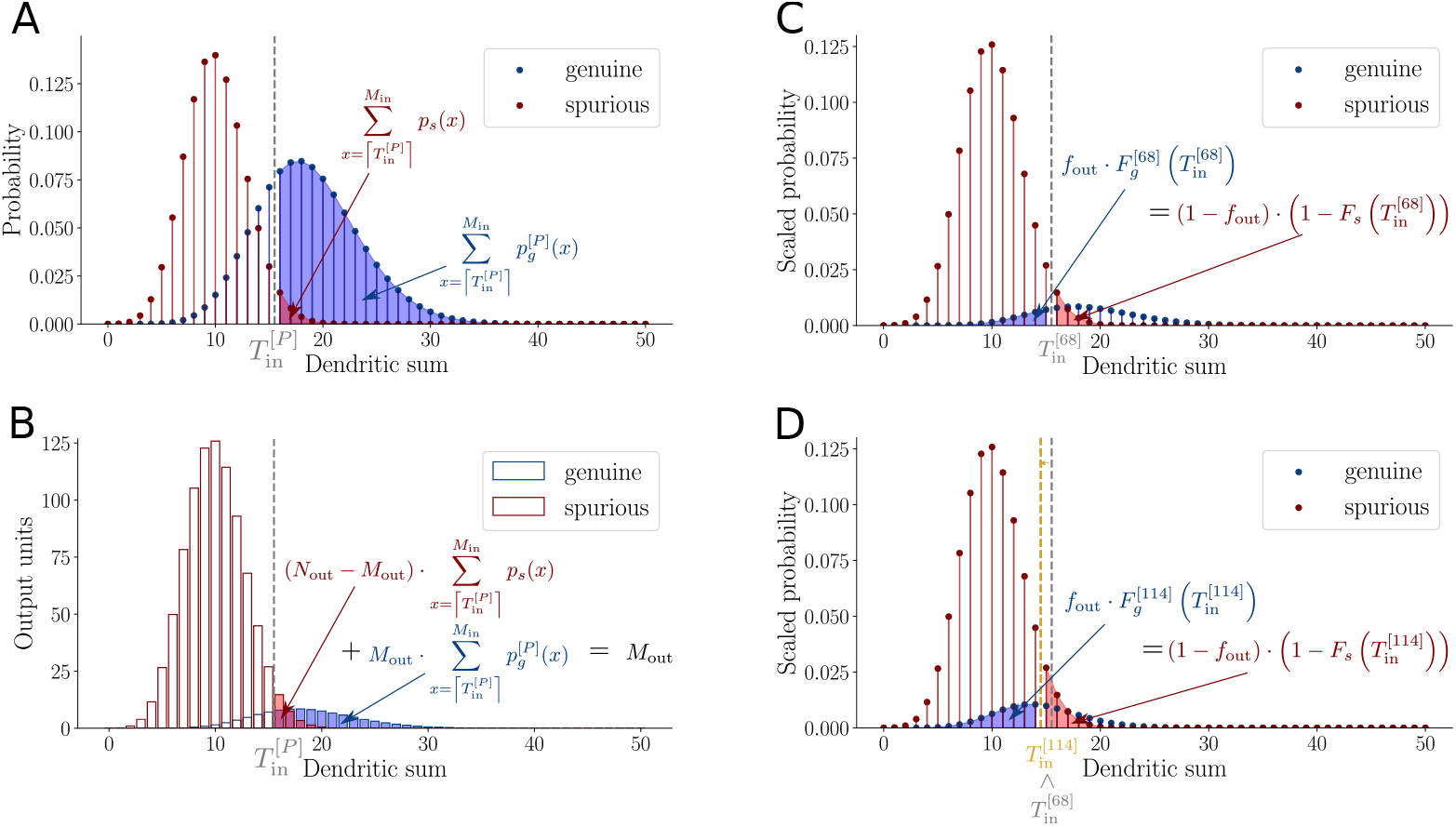
The activation threshold and balancing tails of the distributions. (**A**) Distributions of dendritic sums of genuine (blue) and spurious (red) output units. Output units with a dendritic sum larger than the activation threshold 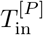 are activated. (**B**) Distributions of dendritic sums scaled by the number of units whose dendritic sums follow the respective distributions. The sum of the number of genuine (blue) and the number of spurious (red) units that have a dendritic sum larger than the threshold 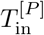 corresponds to the predefined number of active output units *M*_out_. (**C**) Distributions of dendritic sums of genuine (blue) and spurious (red) units scaled by *f*_out_ and 1 −*f*_out_, respectively. The mass of the part of the genuine distribution left of the activation threshold 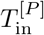 (blue area) must correspond to the mass of the part of the spurious distribution right of 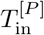 (red area). (**D**) Same as (**C**) but for larger number of patterns *P* = 114 instead of *P* = 68. The threshold 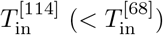 moves to the left in order to maintain the balance of the tails, which ensures the right amount of active output units.

With an increasing number of patterns *P*, the center of mass of the genuine distribution 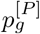 moves to the left, while the spurious distribution stays constant. For a fixed *T*_in_, the left-hand side of Eq. (92) would hence grow while the right-hand side would not, which would lead to an output activation ratio that is smaller than *f*_out_. In order to keep the Balance Equation fulfilled, the activation threshold 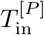 has to move to the left for increasing *P* (Fig 14C, *P* = 68, and 14D, *P* = 114).

### Deriving the Hamming distance from dendritic sums

The Hamming distance

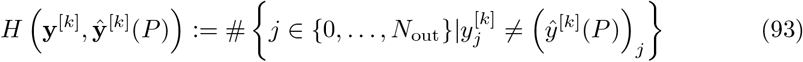

is the sum of the number of genuine units that are wrongly set to 0 and the number of spurious units that are wrongly set to 1 in the calculated output **ŷ**^[*k*]^(*P*):

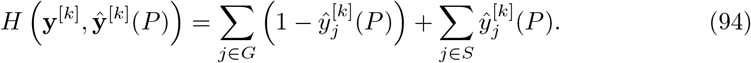

In terms of the distributions *p*_*s*_ and 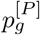, the expected Hamming distance reads as

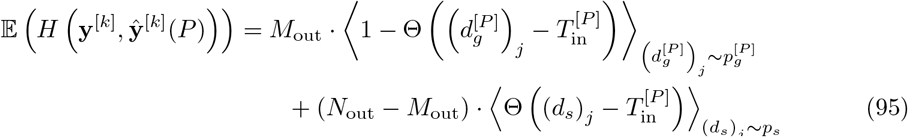

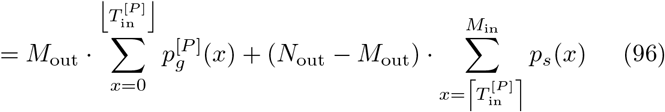

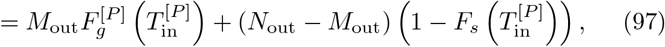

see Fig 15A. The Hamming distance is hence the sum of the part of the genuine distribution that lies below the activation threshold 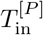 and the part of the spurious distribution that lies above 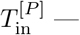 weighted by the corresponding number of units. Using the Balance Equation (92), we can express 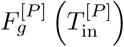 in terms of 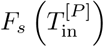, and the expected Hamming distance simplifies to

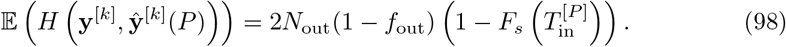

**Fig 15.**
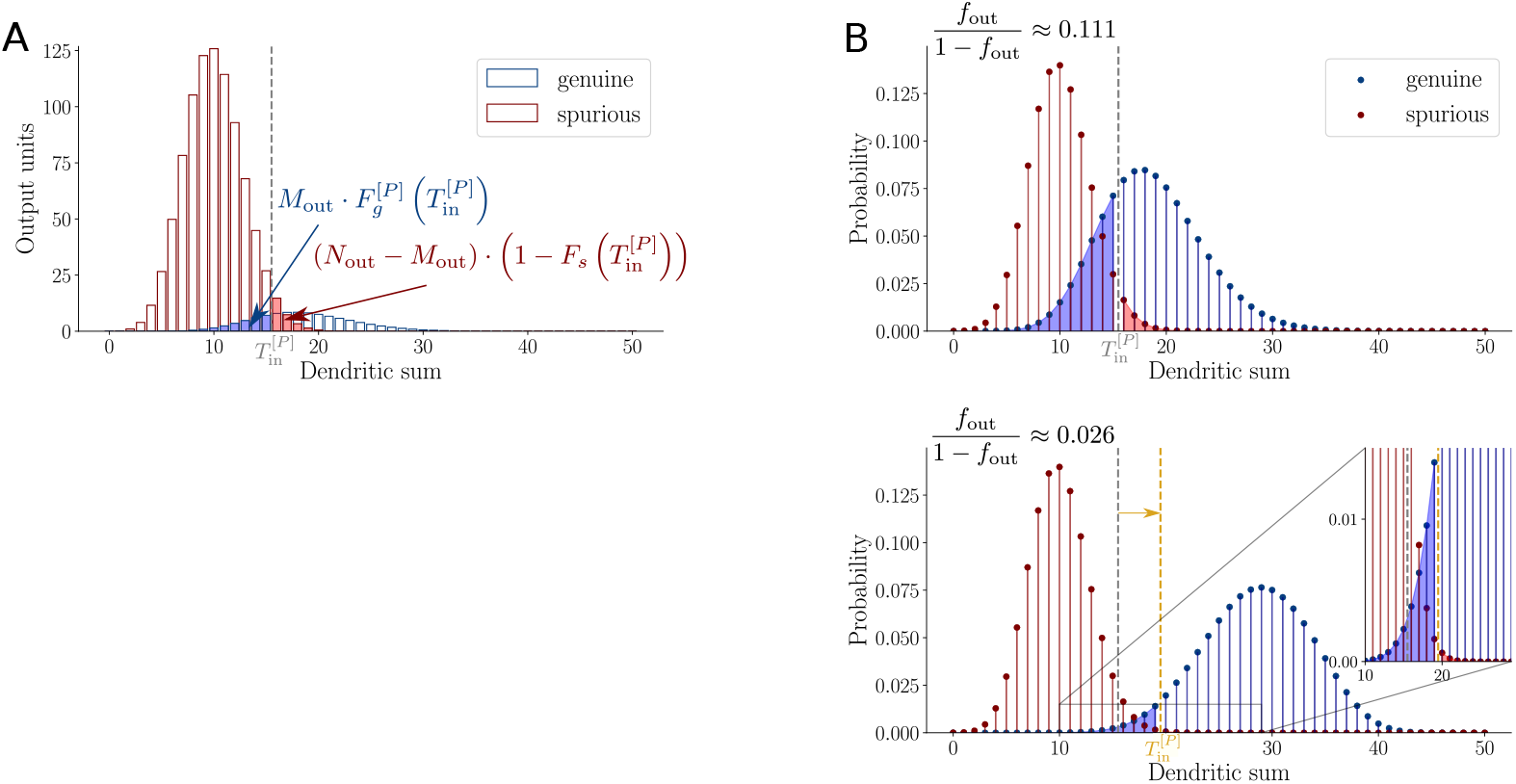
Deriving the Hamming distance from dendritic sums. (**A**) Hamming distance is the sum of falsely deactivated genuine units (blue area) and falsely activated spurious units (red area). (**B**) If *f*_out_ is changed, the threshold 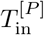 needs to be adapted such that the ratio of the red area to the blue area corresponds to the ratio 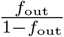. 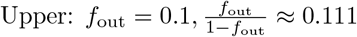,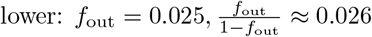 and hence the red area is much smaller compared to the blue area in the lower panel than in the upper panel. This is achieved by a larger 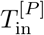, which can also affect the Hamming distance.

The dependence on the number of patterns *P* is now solely contained in the activation threshold 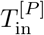 because *F*_*s*_ is constant in *P*. The Hamming distance increases with decreasing 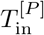 and, as discussed previously, 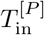 decreases with increasing *P*. Thus, the larger the number of subsequently learned patterns *P*, the larger the Hamming distance between target output and calculated output and, according to Eq. (76), the smaller the signal quality *S*_*P*_.

Note that the dependence of the Hamming distance on the output activation ratio *f*_out_ is less straightforward. On the one hand, if *f*_out_ is increased, the factor 1 − *f*_out_ in Eq. (98) yields a decrease in the Hamming distance. On the other hand, an increase of *f*_out_ also decreases the activation threshold 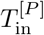 and thus increases the factor 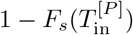 in Eq. (98), which yields an increase in the Hamming distance (see Fig 15B). It is not easy to determine analytically which of these opposing effects is stronger. As mentioned in the Results, we speculate that the combined effect on the Hamming distance is small in the parameter range that we investigated numerically.

### Signal quality after *P* subsequent patterns

In order to derive the signal quality *S*_*P*_ = *s*_*P*_ *H*_avg_ (see Eqs. (76) and (77)) after *P* additional learning steps, we combine Eqs. (78) and (98) and obtain

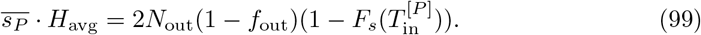

Inserting *H*_avg_ := 2*N*_out_*f*_out_(1 − *f*_out_) yields

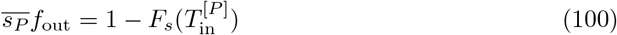

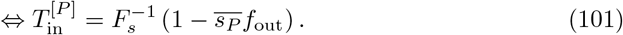

Using Eq. (100), we find that the Balance Equation (92) is fulfilled if and only if

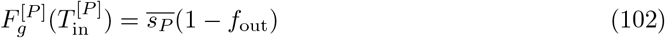

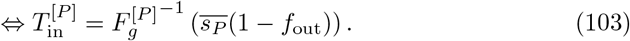

Fig 16 illustrates the two equations. We can equate the right-hand sides of Eqs. (101) and (103) to get rid of the explicit value of 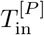 and obtain a single equation

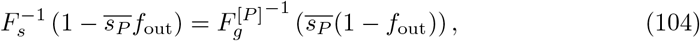

which we call the Signal Quality Equation. If this equation is solved for 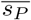, inserting its solution into Eq. (77) with 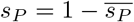 determines the signal quality *S*_*P*_. There is an explicit solution if 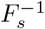 and 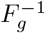 are easy to calculate. An analytical solution to this equation, which requires several approximation steps, is outlined in S2 Appendix. Alternatively, the Signal Quality Equation can be solved numerically (see next subsection).

**Fig 16.**
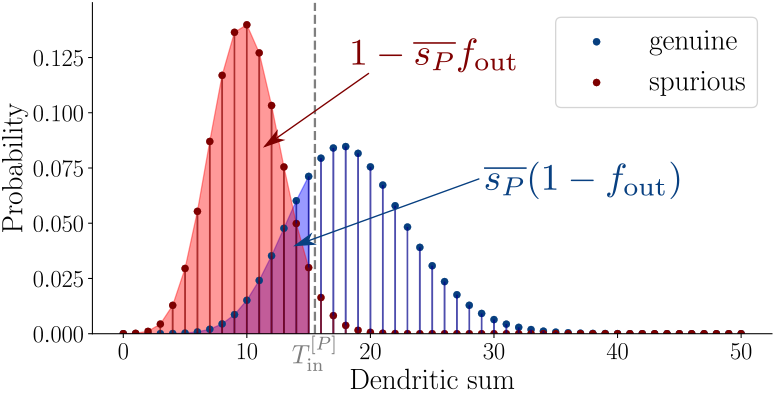
Signal Quality Equation. The activation threshold 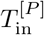 corresponds to the 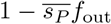-quantile of the spurious distribution and to the 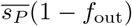-quantile of the genuine distribution.

Notably, since the CDF of a discrete probability distribution is a piecewise constant function, the CDFs in Eqs. (102) and (102) are not generally invertible. A strict inverse *F* ^*−*1^(*y*) such that *F* (*F* ^*−*1^(*y*)) = *y* does not always exist because the same CDF value *F* (*x*) can correspond to multiple arguments *x*. Being aware of this fact, we use the notation 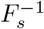 and 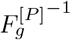 here anyway. Later in this section, the binomial distributions (see Eqs. (61) and (63)) are approximated by normal distributions, and their approximated CDFs become strictly increasing and continuous, and thus invertible.

### Numerical solution of the Signal Quality Equation

The Signal Quality Equation (104) could be solved numerically. As discussed above, the CDFs *F*_*s*_ and 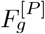 are not invertible. Thus, we approximate the discrete distributions 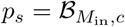 and 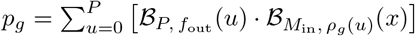 by continuous distributions

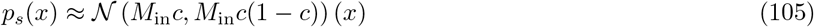

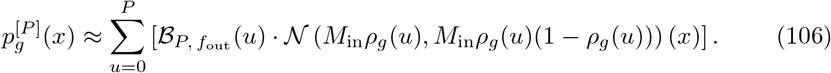

This is a good approximation for large enough *M*_in_ and sparse but not too sparse connectivity *c*. Specifically, we have to assume

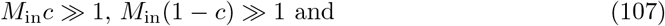

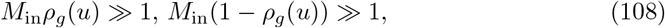

for all *u* ∈ {0, …, *P*}. These four assumptions reduce to the two assumptions

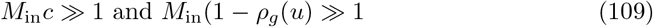

because *ρ*_*g*_(*u*) ≥ *c*. Then, the Signal Quality Equation (104) can be written as

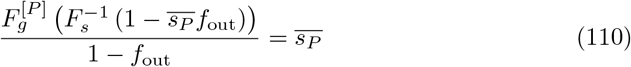

and easily solved numerically without any additional approximation steps. This semi-analytical strategy yields a better approximation of the results from network simulations (see Fig 17) than the signal quality based on a fully analytical description (see S2 Appendix).

**Fig 17.**
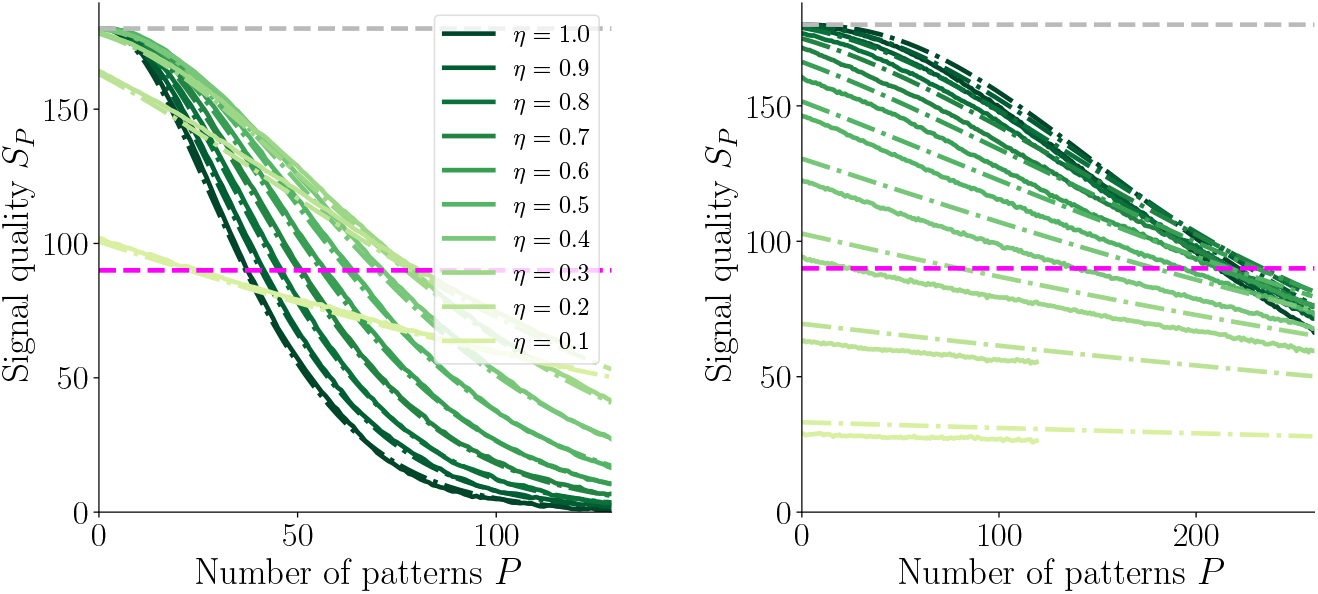
Comparison of signal quality obtained from network simulations and from a semi-analytical approximation. Signal quality decays with number *P* of subsequently learned patterns *P* for various values of transition probability *η*. Solid lines: numerical network simulations dash-dotted lines: semi-analytical estimate (as discussed in this subsection) of the signal quality; dashed grey line: maximal signal quality, which is the average Hamming distance between two random *f*_out_-sparse patterns; dashed blue line: retrieval threshold. Left: *f*_in_ = 0.1, right: *f*_in_ = 0.012. Other parameters: *N*_in_ = *N*_out_ = 1000, *f*_out_ = 0.1, *c*_*m*_ = 1, *c* = 0.2.

### Memory capacity of the network

Instead of calculating the signal quality at any *P* subsequent patterns (see previous section), which can then be used for finding the *P* for which the signal quality reaches the retrieval threshold, the network capacity can be determined by directly equating the signal quality to the retrieval threshold (expressed in terms of *s*_*P*_) and solving the resulting equation for *P*.

We are searching for the number of patterns *P* for which the signal quality as defined in Eq. (76) equals the retrieval threshold *T*_*S*_, which can be expressed as a fraction *t*_*S*_ of *H*_avg_:

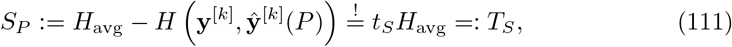

with *t*_*S*_ ∈ (0, 1). In analogy to 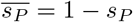, we define 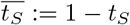 for convenience. Instead of calculating 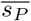 for a given *P*, we now use the same Signal Quality Equation (104) to find the number of patterns *P* for which 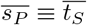 (which is equivalent to *s*_*P*_ ≡ *t*_*S*_) and 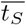 is given. We thus want to solve the Capacity Equation

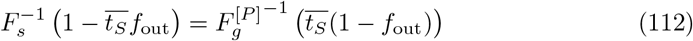

for *P*, which is the only unknown in this equation for a fixed set of network parameters. Notably, the Signal Quality Equation (104) and the Capacity Equation (112) are of the same form but, in the first, the number of patterns *P* is given and the fraction 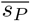, which determines the signal quality 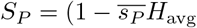, is unknown while, in the second, the retrieval ratio *t*_*S*_ is given and the number of patterns *P*, which corresponds to the capacity, is unknown.

### Approximation of 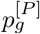 by a single binomial distribution

In the following, we will approximate the distributions of dendritic sums such that the inverses of their distribution functions *F*_*s*_ and 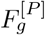 can be calculated. First, the linear combination of binomial PMFs in 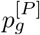 in Eq. (63) is approximated by a single binomial PMF 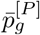 (note the overline of the symbol *p* in the approximation).

A simple way to do so is by choosing the binomial distribution 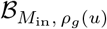 that provides the largest contribution because it has the largest weight. The largest weight is 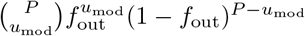 with *u*_mod_ = ⌊*f*_out_(*P* + 1)⌋. We hence obtain the approximation

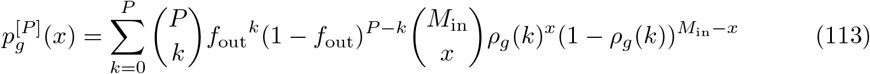

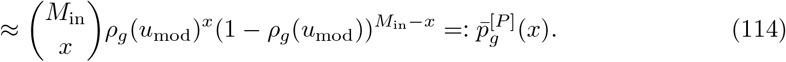

We are ultimately interested in a good approximation of the cumulative distribution function (see Capacity Equation (112) and Fig 16). We denote the cumulative distribution function of the approximation (114) as 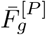 (note, again, the overline). Fig 18 shows a numerical comparison of the summed probabilities

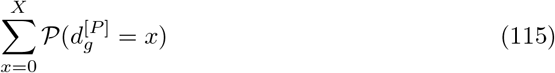

up to various values of *X* for the original distribution 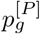 and the approximation 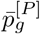 (Eq. (114)) in terms of the absolute difference between the two

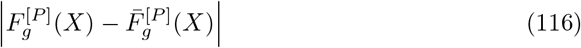

and of the relative error

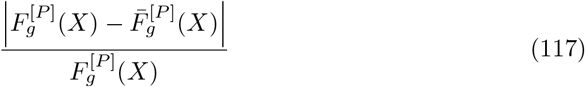

as functions of *P* for several values of *X* between *M*_in_*c* and *M*_in_*c*_*m*_. The non-continuities in the errors as functions of *P* occur due to the discrete jumps in *u*_mod_ = ⌊*f*_out_(*P* + 1) ⌋. The approximation by an individual binomial distribution is better for smaller transition probabilities *η* (compare first to second and third to fourth row in Fig 18). As discussed in S1 Appendix, a large *f*_in_ (above all if combined with a transition probability *η* that is close to one) leads to a more pronounced multimodality of the distribution for small *P* values which fades out more quickly with increasing *P* due to the steeper slope of *ρ*_*g*_(*u*) for small values of *u* (compare Fig 18 upper left to upper right). The more pronounced the multimodality, the less accurate the approximation by a single binomial distribution can be. The output activation ratio *f*_out_ has a different effect on the quality of the approximation than the input parameters. Increasing *f*_out_ increases *u*_mod_. A large *u*_mod_ allows for a low multimodality already for small values of *P* because the strongest weights are quickly in a range where *ρ*_*g*_(*k*) is not extremely steep anymore but approaching the asymptotic value *c* (compare Fig 18 upper left and upper center). We would like to emphasize that decreasing the morphological connectivity *c*_*m*_ strongly improves the match of 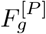 and 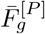 (compare Fig 18 upper left to lower right). Most figures in this manuscript show results for *c*_*m*_ = 1. In terms of the approximation of the distributions of dendritic sums by a binomial distribution, this is the worst-case scenario. In biological nervous systems, there is typically no all-to-all morphological connectivity observed. The analytical approximation derived in this section hence improves significantly for more realistic values of *c*_*m*_ ≪ 1.

**Fig 18.**
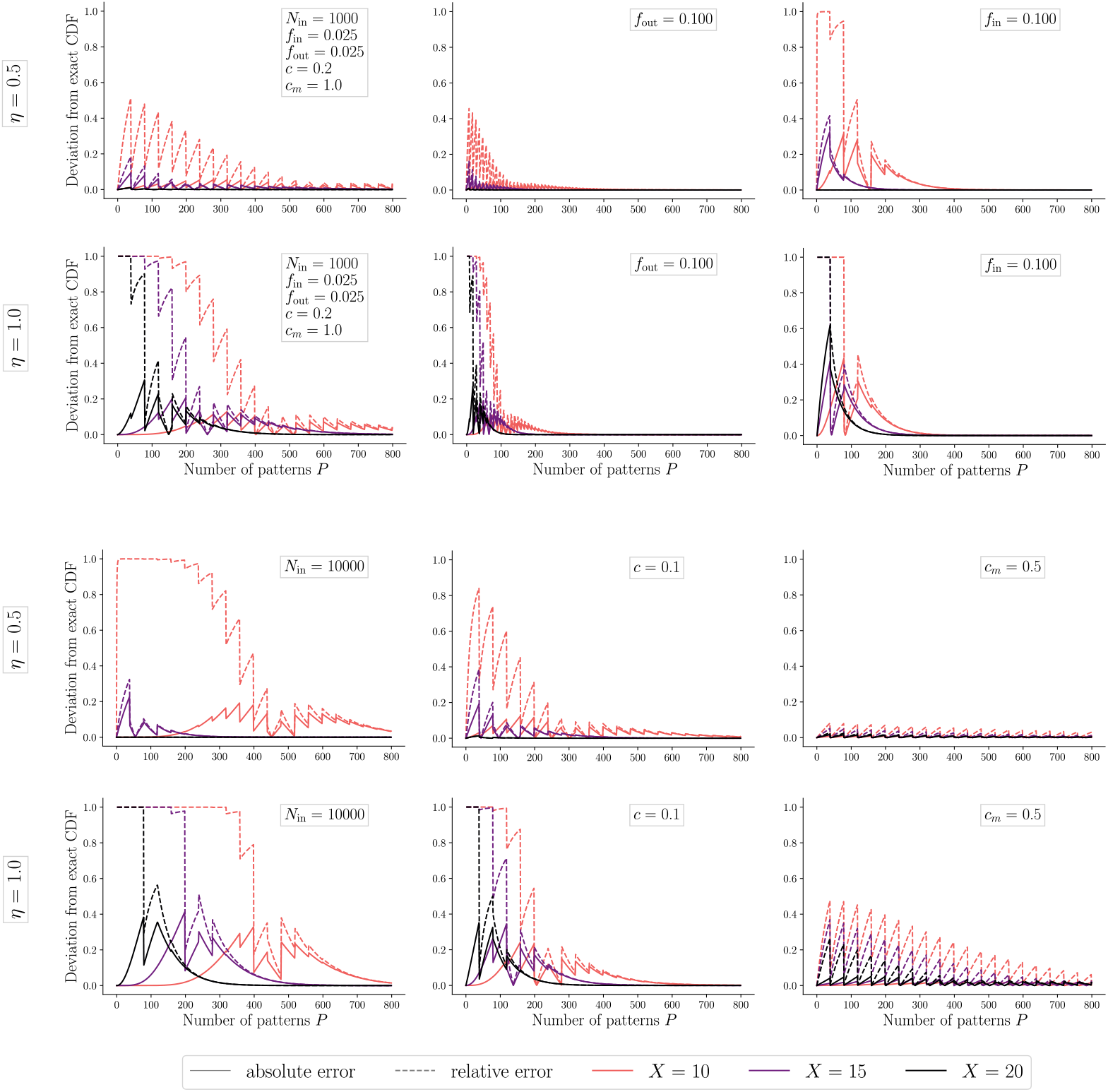
Error of approximation of 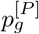 by a single binomial distribution. Absolute (solid lines, Eq. (116)) and relative (dashed lines, Eq. (117)) error of the approximation of 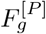 by the cumulative distribution function 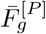 of a binomial distribution (Eq. (114)) for various parameter combinations. For three different values of *X* (argument of the CDF) between *cM*_in_ and *c*_*m*_*M*_in_. In general, the errors decrease with an increase of *X* and with an increase of the number of patterns *P*. Default parameters: *N*_in_ = 1000, *f*_in_ = *f*_out_ = 0.025, *c* = 0.2, *c*_*m*_ = 1 (upper left). First and third row: *η* = 0.5, second and fourth row: *η* = 1. Upper center: *f*_out_ = 0.1, upper right: *f*_in_ = 0.1, lower left: *N*_in_ = 10^4^, lower center: *c* = 0.1, lower right: *c*_*m*_ = 0.5.

### Approximation of binomial distributions by normal distributions

In what follows, we approximate the binomial distributions *p*_*s*_ (Eq. (61)) and 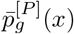 (Eq. (114)) by normal distributions to be able to solve the Capacity Equation (112) analytically. This assumption allows us to derive explicit expressions, which are excellent approximations for the regime of sparse input patterns but large input layer sizes and not too sparse connectivity. Similar to the assumptions for the numerical solution of the Signal Quality Equation, we assume

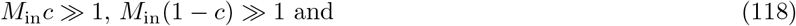

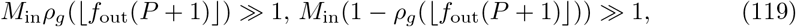

for all *P*, which can be reduced to the two assumptions

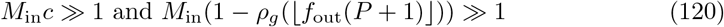

because *ρ*_*g*_(⌊*f*_out_(*P* + 1)⌋) ≥ *c*. (Note that *M*_in_(1 − *ρ*_*g*_(⌊*f*_out_(*P* + 1)⌋)) ≫ 1 is not true for *η* ≈ 1, *c*_*m*_ ≈ 1 and small *P*.)

We approximate the binomial PMFs *p*_*s*_ = ℬ (*M*_in_, *c*) and 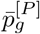, as derived in the previous subsection, by normal probability densities

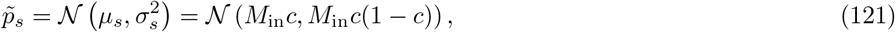

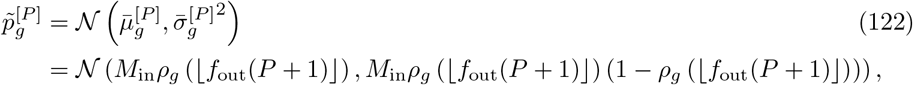

respectively, with cumulative distribution functions

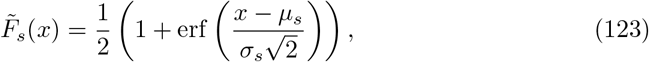

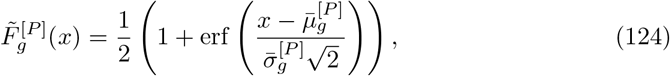

respectively. Note that, in the following, we will often ignore the floor function in the probability density 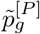 and use *ρ*_g_ (*f*_out_*P*) instead of *ρ*_*g*_ (⌊*f*_out_ (*P* + 1)⌋) because this simplifies some calculations and gives rise to a smooth dependence of the signal quality on *P*. This implies that the capacity can take any non-negative real value and not only integer values that can be interpreted as a number of patterns.

The quantile functions in the Capacity Equation (112) are hence approximated by

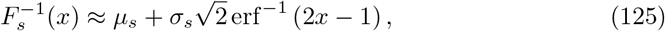

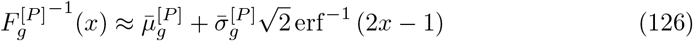

and we obtain

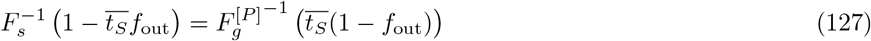

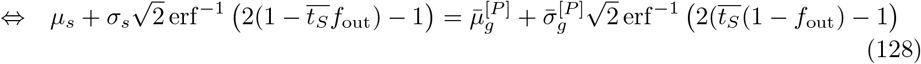

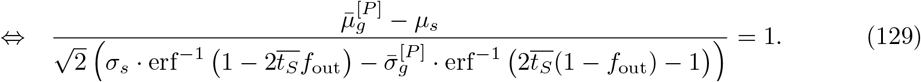

We define the function

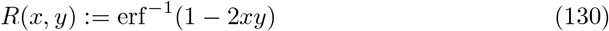

and introduce the abbreviations

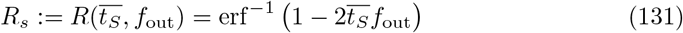

and

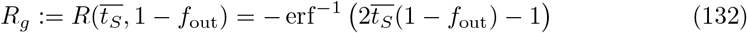

for the weights of the standard deviations in Eq. (129) (Fig 19). Then we have

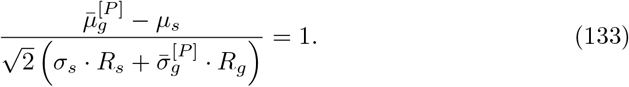

**Fig 19.**
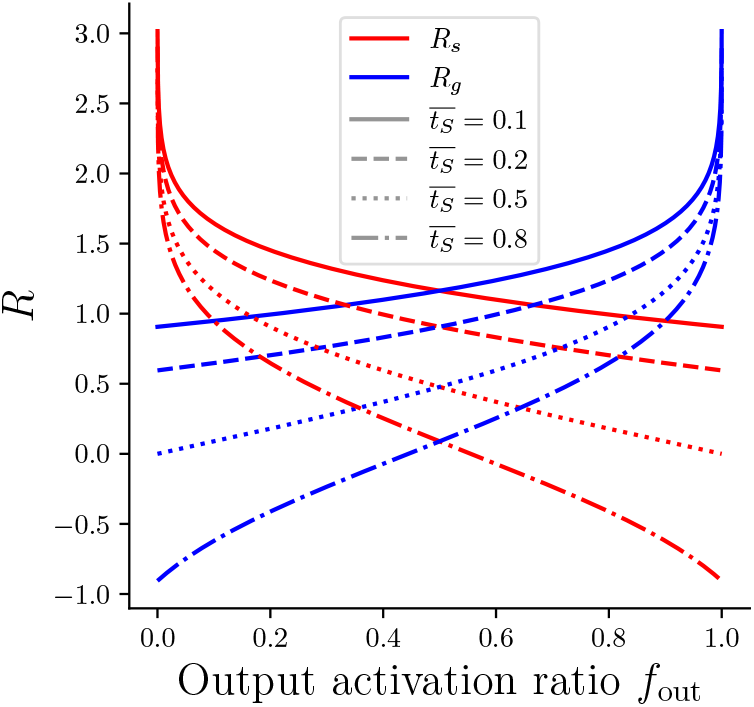
Weights of standard deviations. *R*_*g*_ and *R*_*s*_ as functions of *f*_out_ for 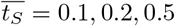 and 0.8. *R*_*g*_ is *R*_*s*_ mirrored at *f*_out_ = 0.5. While *R*_*s*_ is decreasing as a function of *f*_out_, *R*_*g*_ is hence increasing as a function of *f*_out_. Both decrease with increasing 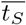. For 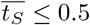, *R*_*g*_ and *R*_*s*_ are non-negative. For *f*_out_ ≤ 0.5, we have *R*_*s*_ ≥ 0 and, for *f*_out_ ≥ 0.5, we have *R*_*g*_ ≥ 0.

Note that, for fixed 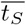, the weights’ dependence on *f*_out_ is symmetric:

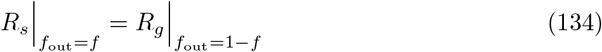

For *f*_out_ *<* 0.5, the prefactor of the spurious standard deviation, *R*_*s*_, is larger than the prefactor of the genuine distribution, *R*_*g*_. For 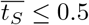, both *R*_*g*_ and *R*_*s*_ are non-negative for all *f*_out_.

The left hand side of Eq. (133) can be interpreted as a generalized version of the sensitivity index

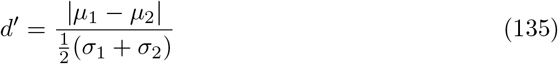

which can be used to quantify the discriminability of two distributions with means *µ*_1_ and *µ*_2_ and standard deviations *σ*_1_ and *σ*_2_. In our case, the additional weighting of the standard deviations by *R*_*g*_ and *R*_*s*_ takes into account the dissimilarly weighted contributions of the false positive and false negative activations due to the unequal amount of genuine units *M*_out_ and spurious units *N*_out_ − *M*_out_ reflected in Eq. (92).

### Solving the Capacity Equation

With the approximations discussed in the previous subsection, the Capacity Equation (112) is equivalent to Eq. (133). Now, we solve Eq. (133) for *P* with the means and standard deviations from the approximate distributions in Eq. (121) and (123):

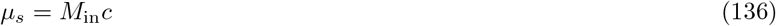

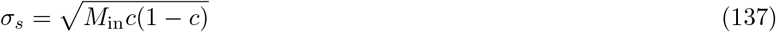

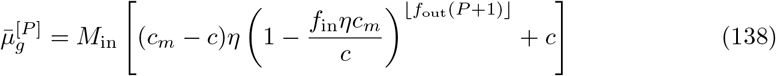

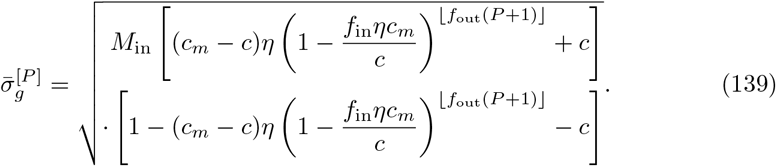

We obtain

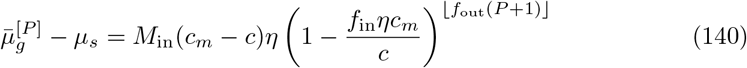

and

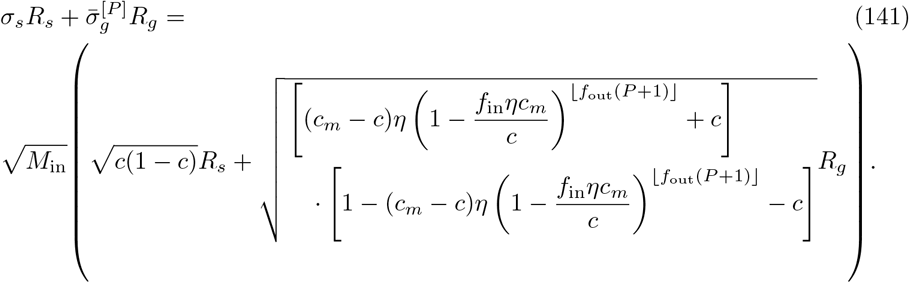

Using the abbreviation 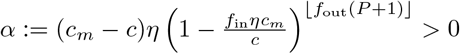, Eq.(133) becomes

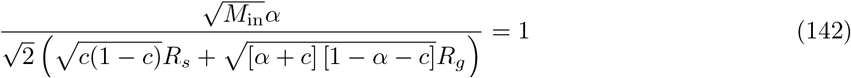

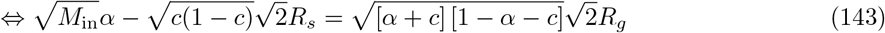

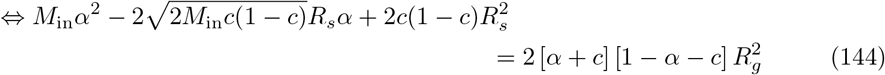

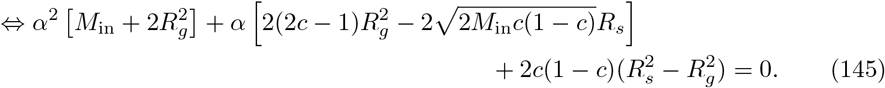

Solving the quadratic equation gives

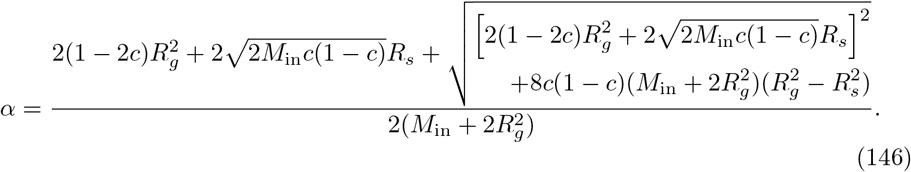

For convenience, we introduce the abbreviations

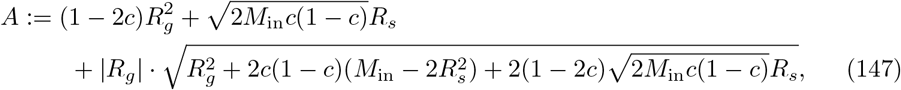

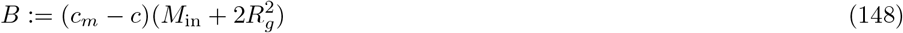

and have

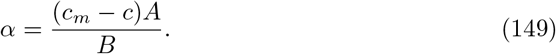

The leading coefficient of the quadratic equation (145) in *α* is positive. For increasing *P*, the term 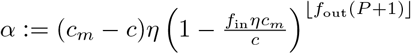 is decreasing. This means that, by increasing *P*, we are approaching zero from the right and we are hence interested in the larger solution of the equation. Therefore, we have to choose the positive sign in Eq. (146). Since *c*_*m*_ *> c, η >* 0 and 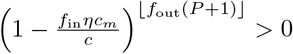, *α* must be positive. Using the positive sign in front of the square root ensures a positive *α*. Note that the expression under the second square root could become negative if *R*_*s*_ becomes too large compared to *R*_*g*_. For *M*_in_ ≫ 1, this is only the case for very small *f*_out_ values combined with a small 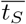. These cases cannot be treated analytically in this way.

Next, we insert 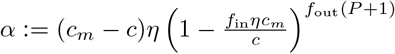 and approximate ⌊*f*_out_(*P* + 1)⌋ ≈ *f*_out_*P* to obtain

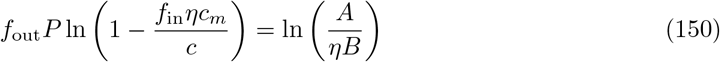

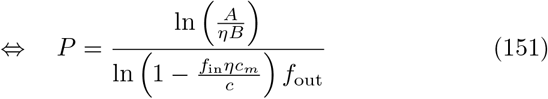

Since *P* is a number of patterns, we round the obtained expression to the nearest integer and obtain the capacity of the network

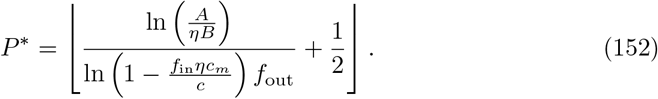

If *R*_*g*_ and *R*_*s*_ are known, this expression allows us to directly calculate the capacity of the network.

### Approximation of the error function

At this point, the constants *R*_*s*_ and *R*_*g*_ (see Eqs. (131) and (132)) could be approximated with the help of any invertible approximation of the error function. We choose the highly accurate global Padé approximation

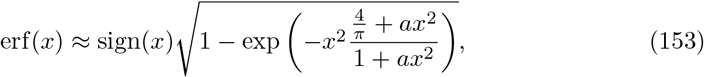

with 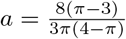 [70], see Fig 20, which allows for the approximation

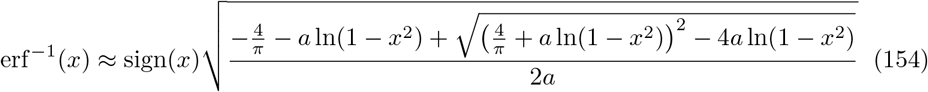

to use for *R*_*s*_ and *R*_*g*_ in Eq. (133). All analytical approximations of the network capacity shown in figures are calculated with this approximation unless stated otherwise.

**Fig 20.**
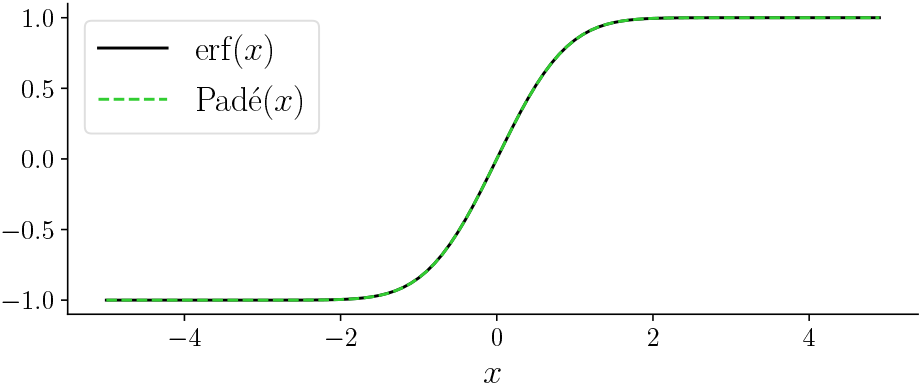
Approximation of the error function by a global Padé approximation. The maximal absolute error is 0.00033, the maximal relative error is 0.00035.

### Optimal transition probability and maximal capacity

From Eq. (152), we can derive the optimal transition probability *η*_opt_ and therewith the maximal capacity 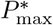 for a fixed set of network parameters.

We assume 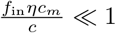 and use the Taylor series of ln(1 − *x*) at *x* = 0

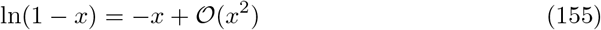

to approximate

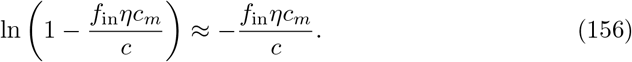

The capacity as a function of *η* then reads as

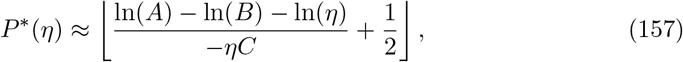

with *A* and *B* as defined in Eqs. (147) and (148) and

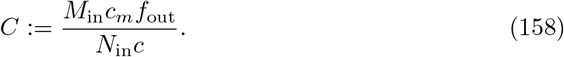

In the following, we will ignore the rounding of *P*^∗^ to an integer value in (157) and allow the capacity to take any (non-negative) real value. We then have

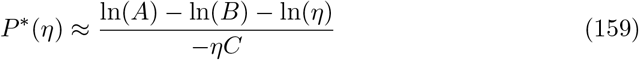

as also stated in the Results in Eq. (19). Next, we calculate the optimal transition probability *η*_opt_ by differentiating *P*\* with respect to *η*

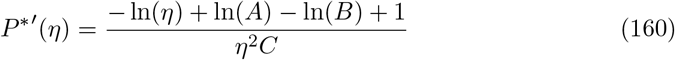

and solving

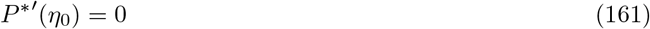

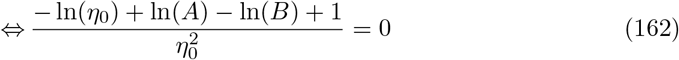

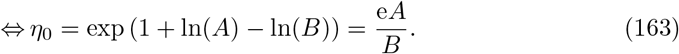

This is a global maximum of *P*\*(*η*) because

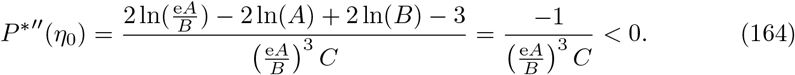

The optimal transition probability must be bounded to (0, 1]. We can assume that 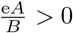 (otherwise there is no real solution for (152)). If 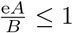, we have 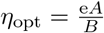. If 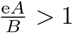, the capacity as a function of the transition probability *P*\*(*η*) is monotonically increasing in the range *η* ∈ (0, 1] that we are interested in because we have shown that *P*\* has a global maximum at e*A/B >* 1. We thus have

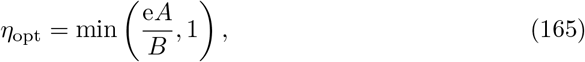

as also displayed in Eq. (20) in the Results. Fig 21 shows *η*_opt_ as a function of the input and output activation ratios *f*_in_ and *f*_out_.

**Fig 21.**
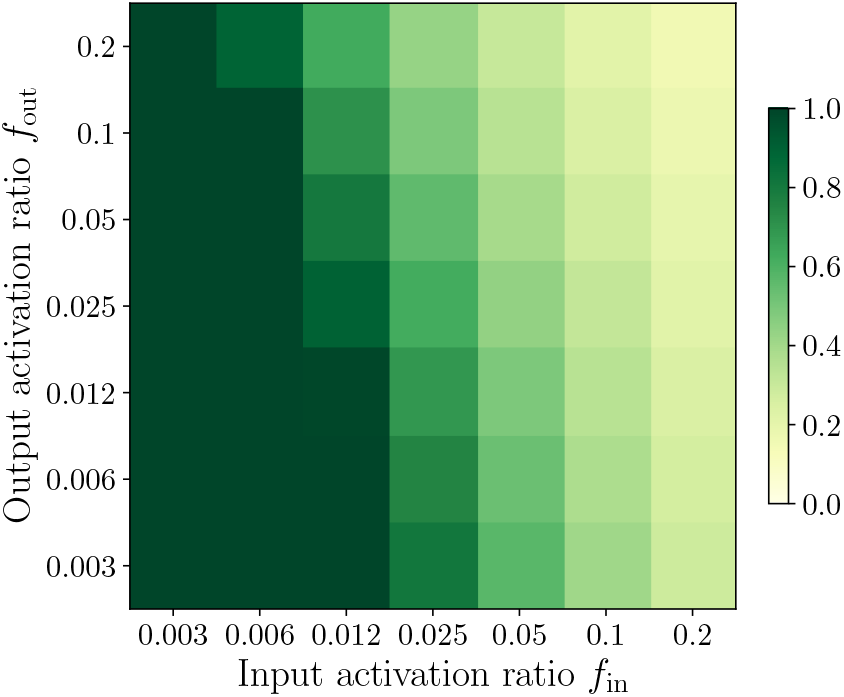
Analytical optimal transition probability. Optimal transition probability *η*_opt_ (Eq. (165)) as a function of *f*_in_ and *f*_out_. *η*_opt_ decreases both as a function of *f*_in_ and as a function of *f*_out_ but the slope is larger for *f*_in_. In Fig 3E in the Results, the corresponding optimal transition probability *η*_opt_ obtained from numerical simulations is shown. (Other parameters: *N*_in_ = 1000, *c* = 0.2, *c*_*m*_ = 1, *t*_*S*_ = 0.5.)

The maximal capacity that can be obtained for a given set of network parameters by optimizing the transition probability is then defined as

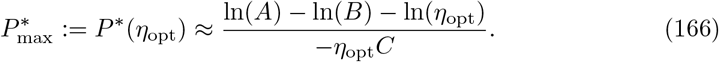

Comparisons of the analytical approximation of the optimal transition probability *η*_opt_ (Eq. (165)) and the maximal capacity 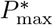 (Eq. (166)) with results obtained from numerical network simulations are presented in Fig 3B,C and Fig 5.

### Dependence on input parameters

Since only *C* = *M*_in_*c*_*m*_*f*_out_*/*(*N*_in_*c*) depends on *N*_in_ while *A* and *B* do not (see Eqs. (147)–(158)), the capacity is proportional to the input layer size *N*_in_ (Fig 5B). Further, we note that the optimal transition probability *η*_opt_ (see Eq. (165)) does not depend on the input layer size *N*_in_, but it does depend on the number of active input units *M*_in_ = *f*_in_*N*_in_ (Fig 5A).

We further observe that the maximal capacity as a function of the number of active input units *M*_in_ has a maximum that is independent of *N*_in_. In the range where *η*_opt_ = e*A/B*, the maximal capacity as a function of *M*_in_

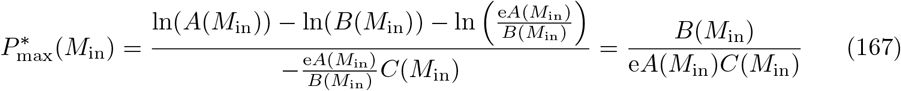

is strictly monotonically decreasing: For *M*_1_ *< M*_2_, we have

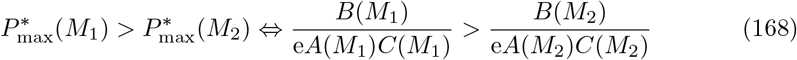

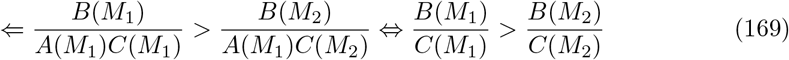

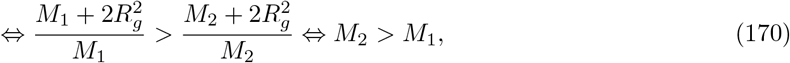

where we used that *A*(*M*_1_) *< A*(*M*_2_) because *A* is monotonically increasing as a function of *M*_in_ (due to *R*_*s*_ *>* 0 for *f*_out_ *<* 0.5) and that *A* has to be positive in order to yield a real valued number for the capacity in Eq. (152). Thus, if there is a maximum of 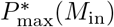, it must occur within the range of *M*_in_ values that have an optimal transition probability of 1 (see Fig 22A-C). In order to find this maximum, we set *η* = 1 in Eq. (159), and then we set the derivative (with respect to *M*_in_) of the maximal capacity

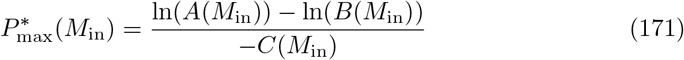

to zero:

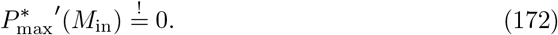

**Fig 22.**
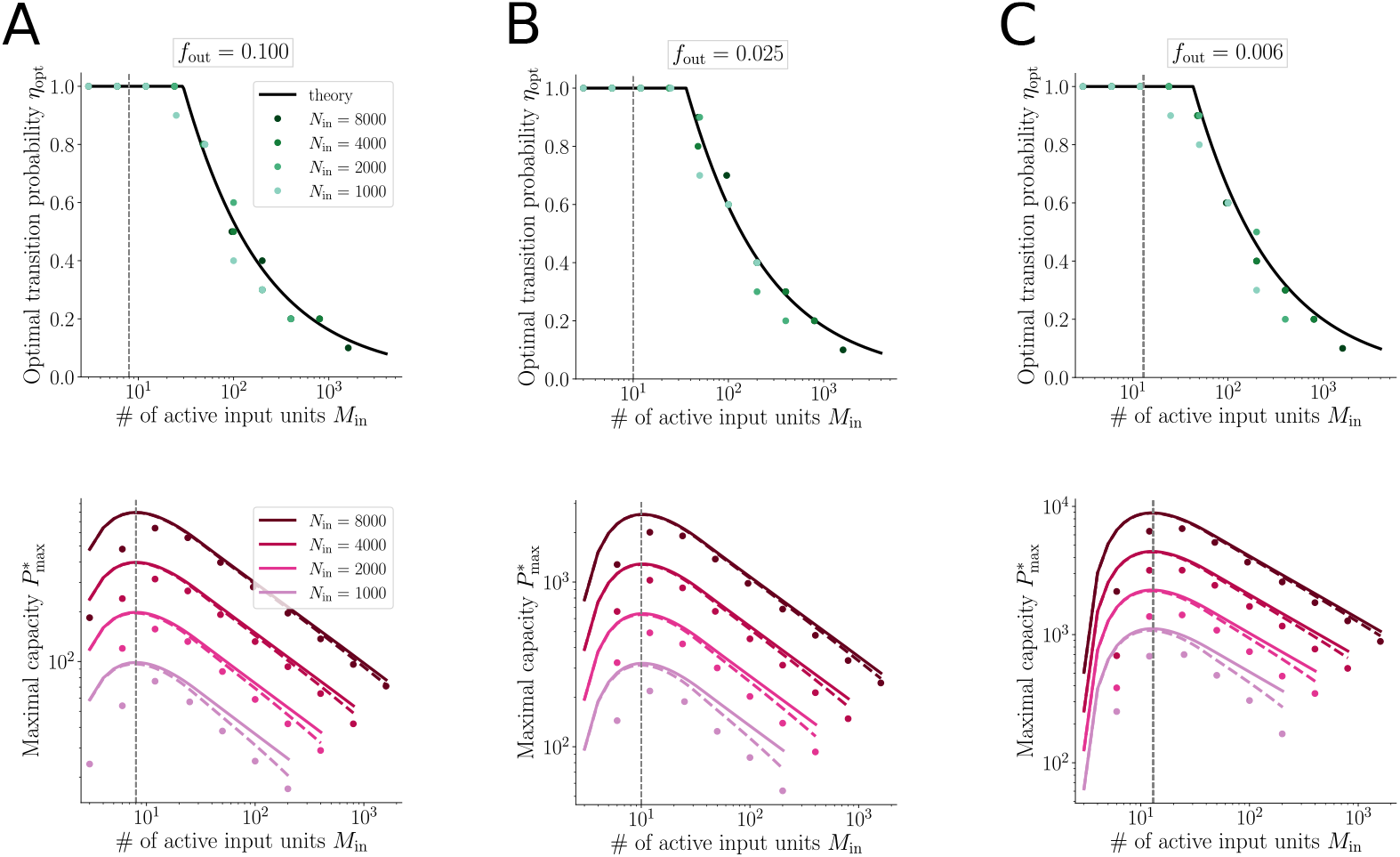
Optimal transition probability and maximal capacity for various *N*_in_. Dependence of optimal transition probability (top) and maximal capacity (bottom) on number of active input units (*M*_in_ = *f*_in_*N*_in_) for various input layer sizes (*N*_in_ ∈ {1000, 2000, 4000, 8000}). Dots: numerical simulations; solid lines: optimal *η* obtained from Eq. (165) and maximal capacity calculated with Eq. (159); dashed lines: maximal capacity calculated with Eq. (152), i.e. without the Taylor approximation (156). In (**A**), *f*_out_ = 0.1, in (**B**), *f*_out_ = 0.025 and in (**C**), *f*_out_ = 0.006. Other parameters: *N*_out_ = 1000, *c* = 0.2, *c*_*m*_ = 1, *t*_*S*_ = 0.1.

This equation can be solved numerically for fixed values of *c, c*_*m*_, *f*_out_, and 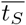. For example, for *c* = 0.2, *c*_*m*_ = 1, *f*_out_ = 0.006 and 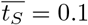, we obtain an optimal number of active input units *M*_in_ = 13, which yields a maximal capacity *P*_max_(13) ≈ 1.11172 ·*N*_in_ (see Fig 22C).

The fit between the theoretical approximation and the actual capacity obtained in numerical simulations improves for increasing input layer size *N*_in_, increasing input activation ratio *f*_in_, and increasing output activation ratio *f*_out_ (see comparison in Table 1). As discussed before, we need *M*_in_ ·*c* ≫ 1 and *M*_in_ · (1 − *ρ*_*g*_(*u*_mod_)) ≫ 1 in order to approximate the distributions of the dendritic sums by Gaussian distributions. A large *f*_out_ reduces the multimodality of the genuine distribution and hence also allows for a better approximation.

### Extension to noisy input patterns during retrieval

In this section, we extend the derivation of the memory capacity and therewith the derivation of the maximal memory capacity and the optimal transition probability to the case of noisy input patterns during retrieval. We assume a noise level *ε* ∈ (0, 1). See earlier sections of the Methods for details on the distributions of dendritic sums with noisy cues; some of the expressions stated there are repeated in what follows in order to simplify the argument.

### Mean and standard deviation

First, we approximate the distribution of dendritic sums of genuine output units

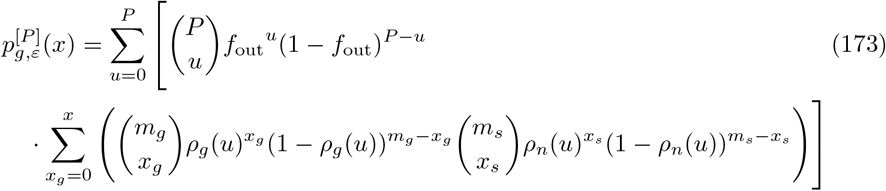

by the renormalized maximum summand (*u* = *u*_mod_) and have

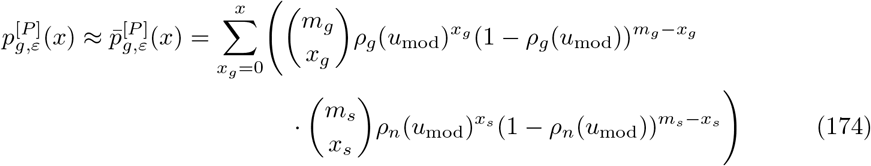

with *u*_mod_ = ⌊(*P* + 1)*f*_out_⌋.

We use again the abbreviation 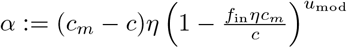. As suggested by [69], the mean and standard deviation of 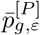 are approximated as

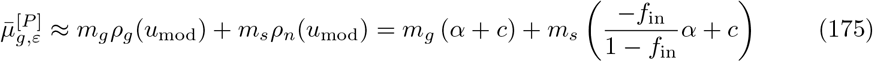

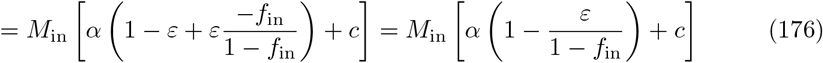

and

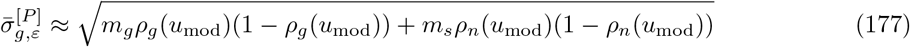

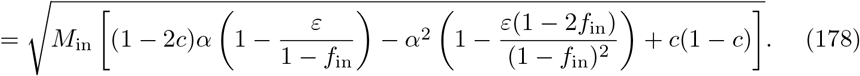

### Memory capacity of the network

As for the case without noise, we solve Eq. (133) for *P* to obtain the capacity of the network. With 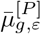 and 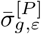, I have

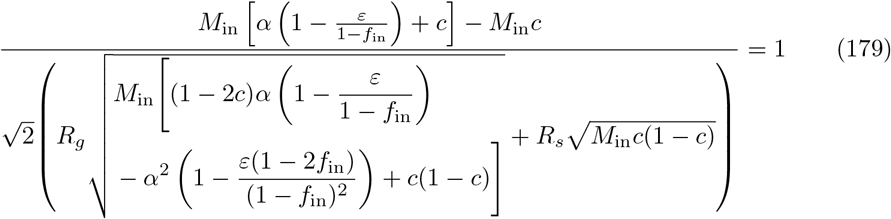

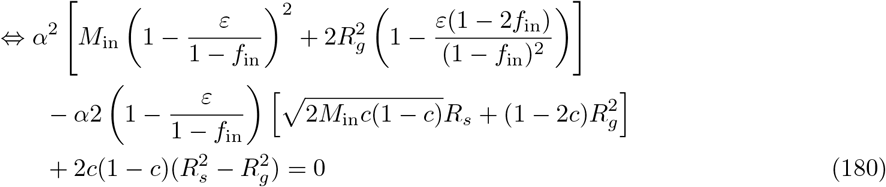

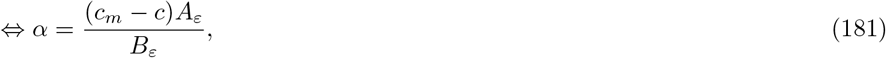

with the abbreviations

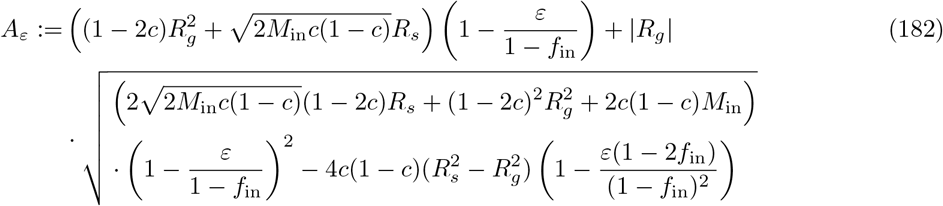

and

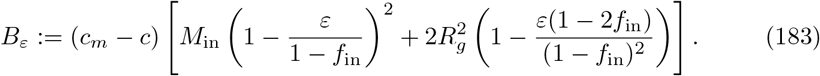

These results generalize the expressions for *A* and *B* in Eqs. (147) and (148), which were derived for *ε* = 0. Expanding ln(1 − *x*), we obtain the capacity

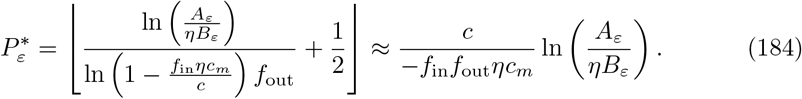

### Optimal transition probability and maximal capacity

The optimal transition probability *η*_opt_ and hence the maximal capacity 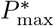 of the network can be calculated in the same way as without noise.The only difference lies in the expressions *A*_*ε*_ and *B*_*ε*_ that were previously called *A* and *B*. Expression *C* = *M*_in_*c*_*m*_*f*_out_*/*(*N*_in_*c*) remains the same as in Eq. (158). We have

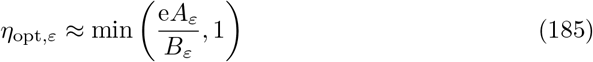

and

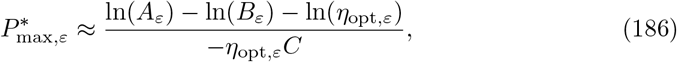

as also stated in Eqs. (22) and (25) in the Results.In the range where *η*_opt,*ε*_ *<* 1, we thus can express the maximal capacity as

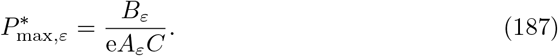

We can assume that 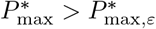, where 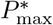 is the maximal capacity for *ε* = 0 defined in Eq. (159), because the additional noise on the input patterns during retrieval naturally has a detrimental effect on the capacity. In the range of parameters where *η*_opt_ *<* 1 (see Eq. (167)) and *η*_opt,*ε*_ *<* 1, we thus have

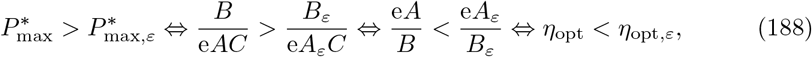

since *C >* 0 and we can assume *A/B >* 0 and *A*_*ε*_*/B*_*ε*_ *>* 0 in order to obtain a real-valued capacity. The optimal transition probability hence increases with added noise compared to the noise-less case by a factor that will be further approximated in the next section. A comparison of 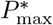 and 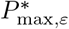 as well as of *η*_opt_ and *η*_opt,*ε*_ is shown in Fig 6D,E and in S4 Fig.

### Approximation of 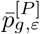 by a single binomial distribution

In order to find a relationship between the capacity 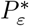 with noise and the capacity *P*\* without noise that is easier to interpret, we can further approximate 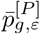 by a single binomial distribution 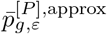 with the same mean

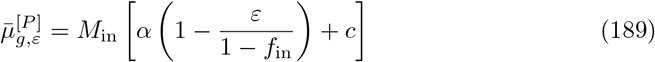

as in Eq. (175). If 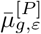 is interpreted as the mean of a binomial distribution 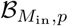, its success probability is 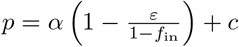 and the corresponding effective standard deviation has to be

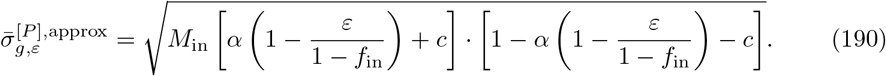

This approximation is exact if the two variances

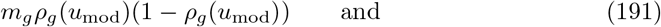

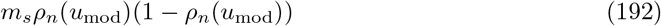

that contribute to 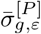 (Eq. (177)) are the same. Inserting 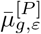 and 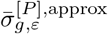 in Eq. (133) and using the abbreviation 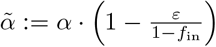 yields the equation

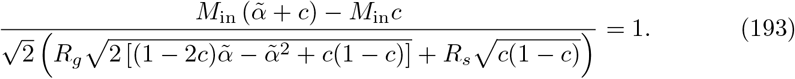

This equation in 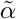 is identical with Eq. (142) in *α* and thus has the solution

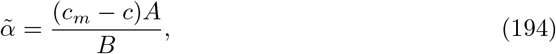

with *A* and *B* being exactly the same as without noisy input patterns (Eqs. (147) and (148)). From this we obtain the capacity

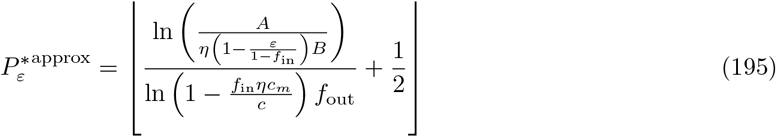

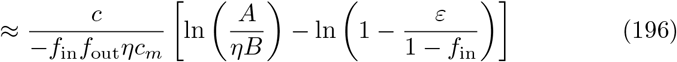

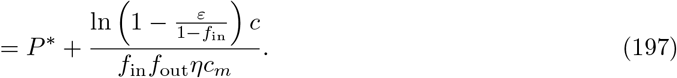

The optimal transition probability with noise during retrieval *η*_opt,*ε*_ can thus be approximated as a multiple of the optimal transition probability without noise *η*_opt_ by

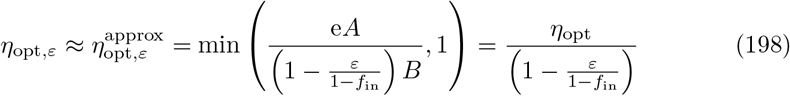

and the maximal capacity 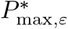 by

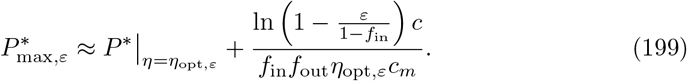

For *ε* ≪ 1 and *f*_in_ ≪ 1, we further obtain the approximation

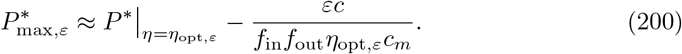

See Fig 6D,E and S4 Fig for these approximations of *η*_opt,*ε*_ and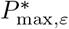.

### Network simulations

In this section, we describe the network simulations performed to obtain the numerical results presented in this paper. First, we report how the main parameters were sampled. Then, we explain the simulation paradigm in detail. Last, we clarify how we average across results for different patterns before briefly broaching the issue of enforcing the output activation ratio.

All simulations were performed in python 3.9. All code written in support of this publication will be made publicly available at https://itbgit.biologie.hu-berlin.de/auer/sparseness_plasticity_lifetime before publication.

### Parameter sampling

Unless stated otherwise we perform all simulations for transition probabilities

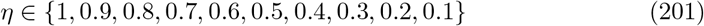

and define the value for which the capacity is the largest as the optimal transition probability. For the reference network size of *N*_in_ = *N*_out_ = 1000, the activation ratios are typically sampled as

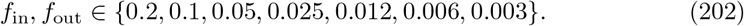

For larger network sizes, smaller values *f*_in_ and *f*_out_ are added to achieve minimal *M*_in_ and *M*_out_ values comparable to those of the small reference network.

### The simulation paradigm

We now describe how patterns are sampled and how the weight matrix is initialized as well as how a simulation step is executed and how many steps are carried out in total. As also explained in the Results, we aim at quantifying the memory capacity of the network by tracking the signal quality, which depends on the Hamming distance between a calculated output vector and the corresponding target output vector. The capacity is defined as the number of subsequently learned patterns for which the signal quality reaches a specific value. In each simulation step the association between one pattern pair is learned, and in order to calculate the capacity we need a number of simulation steps that is larger than the capacity itself.

### Input and output patterns

Input patterns are generated as vectors **x** of length *N*_in_ by randomly choosing *M*_in_ entries to be one while the others are zero. Analogously, output patterns are random vectors **y** of length *N*_out_ with *M*_out_ entries equal to one. One random input pattern **x** together with one random output pattern **y** constitute a pattern pair (**x, y**). Note that we do not explicitly exclude the possibility of generating the same vector twice as an input or as an output pattern, but for *M*_in_, *M*_out_ ≫ 1 and *N*_in_ ≫*M*_in_, *N*_out_ ≫*M*_out_, this is very unlikely.

If noisy input patterns are used as retrieval cues, they are generated based on the noise-less input patterns in the following way: Each originally active input unit is deactivated with probability *ε*. We count the number of deactivated units and randomly choose the same number of originally inactive input units to be activated. The number of active input units *M*_in_ is thus maintained.

### Initialization of the weight matrix

First, we generate a matrix 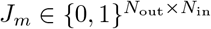 that represents the morphological connections with connectivity *c*_*m*_. In every row of *J*_*m*_, we set *c*_*m*_*N*_in_ randomly chosen entries to one, and the other entries are zero. The binary matrix *J*_*m*_ acts as a mask on the actual weight matrix *J* in every update step such that only the entries in *J* that correspond to a one in *J*_*m*_ can be altered. The weight matrix *J* of size *N*_out_ × *N*_in_ is initialized such that per row *cN*_in_ entries of those that are morphologically available (i.e., have a value of one in *J*_*m*_) are set to one, the rest is set to zero.

### One simulation step

Each simulation step *k* corresponds to one pattern pair (**x**^[*k*]^, **y**^[*k*]^) being learned. In one simulation step *k*, the weights of the matrix in the previous step, *J* ^[*k−*1]^, are first updated according to Hebbian learning and then updated according to the homeostatic mechanism (described in the Results).This corresponds to transforming *J* ^[*k−*1]^ into *J* ^[*k*]^.

Then, for a fixed *J* ^[*k*]^, we evaluate the output of the network for all input patterns of pairs 0, …, *k* that have been learned up to this point; the input patterns are presented to the network again one after the other with plasticity off, and the corresponding output patterns are calculated. By **ŷ**^[*i*]^(*j*) we denote the output of the *i*-th input pattern calculated after learning *j* subsequent pattern pairs. To summarize, the outputs

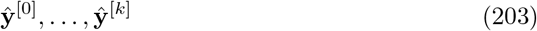

for each of the input patterns

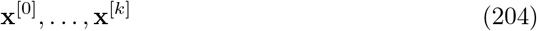

are calculated and compared to the original target outputs

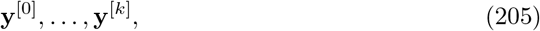

respectively. The *k* + 1 Hamming distances

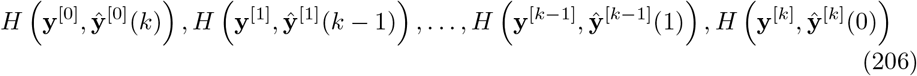

between target and calculated output are stored. Note that these Hamming distances that are stored after one particular simulation step belong to different numbers of additional pattern pairs that are stored after the pattern that is tested. With respect to pattern 0, the maximum number *k* of additional pattern pairs have been stored, while with respect to pattern *k*, no additional pattern pair has been stored.

### Number of simulation steps

The average Hamming distance between two random *f*_out_-sparse vectors of length *N*_out_ is

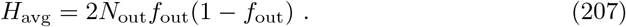

To define a threshold *T*_*S*_ for the successful retrieval of an output pattern, we use a certain fraction of *H*_avg_,

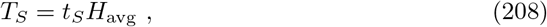

where *t*_*S*_ ∈ (0, 1). If not stated otherwise, the retrieval threshold is chosen as

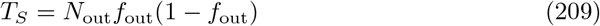

and hence, *t*_*S*_ = 0.5.

The degradation of the calculated output vector is quantified by means of the signal quality *S*, which is defined as the difference between the average Hamming distance of two random vectors, *H*_avg_, and *H*(**y, ŷ**):

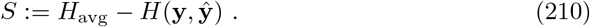

For a successful retrieval of the calculated output, we then require *S > t*_*S*_*H*_avg_. This translates to the assumption that an output pattern can be recovered if the Hamming distance *H*(**y, ŷ**) between the target output **y** and the calculated output **ŷ** is smaller than the threshold (1 − *t*_*S*_)*H*_avg_. In this case, the output, which is typically degraded by subsequent storage of other input-output pairs, is still close enough to the original output pattern. To be more specific, let us consider the example of input-pattern pair 0 and *k* subsequent learning steps. The signal quality is then

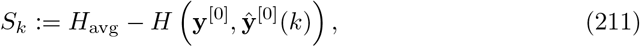

which is based on the Hamming distance (*H* **y**^[0]^, **ŷ**^[0]^(*k*)) of the target output **y**^[0]^ of the very first pattern pair and its output **ŷ**^[0]^(*k*) calculated with weight matrix *J* ^[*k*]^.

The maximum number of simulation steps for a particular parameter combination was determined by the observed degradation of calculated output patterns. Once the signal quality for the first time is less than 0.8 · *H*_avg_ (or equivalently, when the Hamming distance is above 0.2 · *H*_avg_), 200 additional patterns are learned before the simulation is stopped.

We denote the total number of patterns that have been learned by the network at the end of a simulation by *K*.

### Averaging

The signal quality that is shown in various figures in this manuscript is always averaged across many patterns. Recall that the number of subsequently learned patterns *P* is understood as relative to the pattern number *k* ∈ {0, …, *K*}. For each pattern (**x**^[*k*]^, **y**^[*k*]^) that the network learns, we compute the Hamming distance of the target output and the calculated output after *P* = 0, 1, 2, … patterns additionally learned after the *k*-th pattern, hence with weight matrix *J* ^[*k*+*P*]^. For a fixed *P* (always relative to a particular pattern), we average across all patterns for which we have learned at least *P* subsequent patterns:

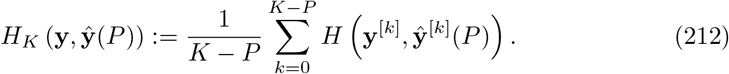

All simulation data shown in this manuscript are obtained by averaging across at least *N*_avg_ = 200 patterns. This means, if the largest *P* shown in a figure is *N*_*P*_, simulations ran for at least *K* = *N*_*P*_ + *N*_avg_ steps.

Whenever we investigate the distributions of dendritic sums for a particular number of subsequent patterns *P*, two kinds of averaging are done: First, we pool the dendritic sums of all genuine output units and of all spurious output units of one pattern after learning *P* subsequent patterns, respectively. In addition, we pool a certain number of patterns. A comparison of distributions of dendritic sums obtained from numerical simulations and from theory (Eqs. (61) and (63)) is shown in S5 Fig.

### A note on the output sparseness

During retrieval, the activation of the output units is calculated as

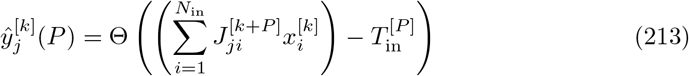

where Θ is the Heaviside step function. Note that, due to the discrete nature of the dendritic sums of the output units, it is unlikely that an activation threshold 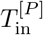 exists that allows to activate exactly *M*_out_ output units. Most of the times, there is a group of units with identical dendritic sums that are exactly at the critical value. If the threshold is chosen larger than these dendritic sums, too few units are activated, if it is chosen smaller, too many units are activated. Since we want to keep *M*_out_ fixed, we have to find a way to enforce the exact output sparseness. Hence, we typically randomly choose the right number of these units to be activated and the rest to be inactive. On average, this has a slightly deteriorative effect on the capacity of the network because spurious units might be chosen to be activated even though some genuine units have the same dendritic sum. In S5 Appendix, we discuss in more detail for which parameters this might have significant effects on the capacity of the network and how the enforcement of the output sparseness level could be handled differently in order to avoid this problem.

## Discussion

We studied the effect of population sparseness on the optimal learning speed in an associative network. We quantified the storage capacity of a heteroassociative feedforward network, where the capacity is based on a threshold of Hamming distance between the target patterns and the calculated output patterns.Furthermore, we determined the transition probability that maximizes the capacity (see Eqs. (19)-(21)).Analytical derivations and numerical simulations show that the optimal transition probability increases with the sparseness of both the input and the output patterns, but the impact of the input sparseness is stronger (see Fig 3B,C,F).The maximum capacity is largest for the lowest *output* activation ratio, whereas it depends non-monotonically on the *input* activation ratio (Fig 3B-D).We found that the maximum capacity is extensive with respect to the input layer size but that the optimal transition probability is independent of the network size (Fig 5).As already hinted at by [71] in a gradient descent framework, we confirm that not the input activation ratio but the actual number of active input units is the crucial parameter that determines the optimal transition probability — independent of the network size (Fig 5A).The number of active input units that yields the largest possible capacity is also independent of the network size (Fig 5). All results hold for recall with perfect input patterns as well as with noisy input patterns. As expected, the maximal capacity decreases as a function of the noise level but, interestingly, the optimal transition probability increases (Fig 6).In contrast to previous works based on single neurons [15] or biologically implausible algorithms such as gradient descent [64, 71], which is non-local, our in-depth numerical and analytical analysis based on Hebbian and homeostatic learning presents a substantial advancement as it reveals the distinct effects of input and output sparseness as well as network size and input noise on the optimal learning speed while relying on a very simple network and pattern structure.

### Palimpsest memory and homeostatic plasticity

The architecture of the network used in this manuscript is similar to the Willshaw network [28] in that it is a feedforward one-layer network that learns associations between binary input and output patterns in a Hebbian way.In contrast to a classical Willshaw network in which all memories disappear when the capacity is reached, our network realizes a palimpsest memory system in which older memories are erased by newer memories due to a homeostasis mechanism (see e.g. 3A). This degradation of older memories allows for an online learning regime where an in principle infinite sequence of patterns can be learned — new patterns always gradually replacing older ones [35, 37]. The signal quality of a memory pattern hence does not only passively decay over time due to the interference between patterns, but also actively due to the deactivation or depression of synaptic connections. The underlying heterosynaptic plasticity, which is characterized by an input-unspecific change of synaptic efficacies [72, 73], has been found, e.g., in the hippocampus and the neocortex [74, 75]. Heterosynaptic plasticity can provide an effective homeostatic constraint on Hebbian plasticity [76, 77]. In the presented work, a combination of Hebbian homosynaptic and homeostatic heterosynaptic learning was used. Hebbian plasticity increases connection weights with a predefined probability if both the pre- and the postsynaptic units are active, and homeostatic plasticity decreases weights with another predefined probability if only the postsynaptic unit is active (details in Methods).The update probabilities, which can be interpreted as learning rates, are balanced such that the total number of functional connections per output unit remains constant. This special heterosynaptic plasticity rule is inspired by experimental evidence of a constant total summed synaptic weight on a dendritic branch [66, 78].

In general, variations of the implemented homeostatic plasticity are possible. For example, in contrast to our approach, the total number of functional connections per input unit could be conserved to realize a presynaptic homeostasis by depressing connections from active input neurons to inactive output neurons (as suggested by [35], e.g., reported by [79]). Alternatively, the number of functional connections could only be conserved across the entire network, i.e. not be normalized per (input or output) unit; such a normalization could be achieved by randomly silencing connections that are not involved in a given association of patterns [36]. Numerical simulations based on either of these two variations suggest that the qualitative dependence of the optimal learning speed on the sparseness of the patterns is the same as for the learning rule analyzed in detail in this paper (see S4 Appendix). Finally, a decrease in synaptic efficacies could occur due to synaptic aging where connections that have not been involved in a pre- and postsynaptic co-activity for a long time might be more likely to be depressed than those that have been strengthened more recently [80].

### Input sparseness vs output sparseness

In our heteroassociative network, both a change in the input and the output sparseness value affect the optimal learning speed and the corresponding maximum memory capacity (Figs 3, 5 and 6, Eqs. (19)-(21), details in Methods).The optimal transition probability monotonically decreases with an increase in the input activation ratio as well as with an increase in the output activation ratio but the gradient is steeper for the input activation ratio (see Figs 3E, 6B and 21). In order to learn fast, sparseness in the input representations is thus more beneficial than sparseness in the output representations. As known from the literature [28, 32, 35], the capacity of an associative network benefits from sparse representations. The same is of course true for the maximum capacity, which is obtained by learning with the optimal transition probability for each combination of parameters. While enforcing sparser output patterns decreases the overlap in the output space and hence decreases the likelihood of retrieval errors up to extreme sparseness values of only one active output unit, using too sparse input patterns can be detrimental to the capacity. If the input representations contain a too small number of active units *M*_in_, it is difficult for the signal, which scales as *M*_in_, to overcome the noise, which scales as 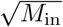 (Figs 3A,B and 5B). In other words, a very small number of active input units cannot reliably drive the respective output units. If *M*_in_ is smaller than some optimal value, the initial signal quality approaches the retrieval threshold, and if *M*_in_ becomes very small, the capacity can be zero because the initial signal quality lies below the threshold [36, 37]. This collapse of the maximal capacity for too sparse input patterns only occurs in the range of *M*_in_-values where the optimal transition probability has already reached its maximal value 1 (Fig 5A, details in Methods). Note that, while the assumption of only binary synaptic weights combined with a transition probability is a simplification of the state transitions occurring in real synapses, assuming a bounded number of discrete states is realistic [81, 82]. It hence makes sense to assume that the learning speed in terms of the amount of potentiation experienced by synapses combined with a probability of potentiation is bound to reach an upper limit.

### Choice of the activation threshold and optimized capacity

The central purpose of this work is to rigorously establish the relationship between the population sparseness of the patterns and the optimal learning speed of the network. In this spirit, for a given output activation ratio, the setup needs to enable a meaningful comparison of target and calculated output patterns in order to quantify the capacity and therewith the optimal transition probability as functions of the activation ratio.

Thus, it is of crucial interest to enforce a particular number of active output units *M*_out_, which is implemented by choosing the activation threshold of the McCulloch-Pitts output neurons accordingly (Eq. (1)).The maximum capacity calculated in this framework (Eq. (21), details in Methods) represents the highest capacity obtained under the given constraints using the optimal transition probability. In other approaches, the beneficial effect of sparseness on the capacity has been studied extensively in different settings. For example, [33] could increase the capacity by using an optimized, covariance-based learning rule. Moreover, [83] suggested the use of a numerically optimized activation threshold in a setup that is very similar to ours. In this manuscript, such aspects have been excluded, in an attempt to keep the model as simple as possible to allow for a detailed analysis of the effect of sparseness on the learning speed. The behavior of the capacity that is reported in the current work is always focused on its relation to the optimal transition probability.

### Role of the output size and retrieval criterion

The analytical description of the memory capacity relies on the distributions of dendritic sums which are based on the synaptic connections of single output units (Eqs. (8) and (9)). For the Hamming distance (Eq. (12)) and thus also for the signal quality *S*_*P*_ (Eq. (5), also see Methods), the number of output units *N*_out_ has to be taken into account as the probabilities of output units being in the wrong state are scaled by the respective number of units. We defined the memory capacity as the number of subsequent patterns that can be learned until the signal quality *S*_*P*_ reaches the retrieval threshold *T*_*S*_ (Eq. (6) and Eq. (15)). The retrieval threshold *T*_*S*_ = *t*_*S*_*H*_avg_, with a fixed *t*_*S*_ ∈ (0, 1), is chosen as a fraction of *H*_avg_ = 2*N*_out_*f*_out_(1 − *f*_out_), the average Hamming distance between two random *f*_out_-sparse patterns of size *N*_out_ (Eq. (7)). Hence, the number of wrong output units that are accepted for the pattern to be classified as retrievable depends on *N*_out_ and *f*_out_. This dependence accounts for the fact that a fixed number of wrongly activated units has a more severe destructive effect if the total number of active output units is small. For example, if in a pattern with *M*_out_ = 100 active output units nine are wrong, the pattern is still very similar to the original one; if *M*_out_ = 10, nine wrong units are detrimental. The capacity is obtained by comparing the signal quality *S*_*P*_ to the retrieval threshold *T*_*S*_ or, equivalently, by comparing the normalized signal quality *S*_*P*_ */H*_avg_ to the retrieval ratio *t*_*S*_. Since the normalized signal quality *S*_*P*_ */H*_avg_ does not depend on *N*_out_ (derived in Methods) but only on the distributions of the dendritic sums and *f*_out_ (see Eqs. (77) and (79) as well as S2 Appendix), the capacity does not depend on *N*_out_ either.

As an alternative, which has not been explored in this manuscript, the retrieval threshold could be defined independent of *H*_avg_ and thus independent of *N*_out_, e.g., implementing the retrieval criterion that a maximum of two units may be wrongly activated or that at least ten units need to be correctly activated. In such alternative cases, the capacity could depend on the output network size *N*_out_.

Furthermore, the output size as well as the input size in combination with the output and input activation ratios, respectively, pose an upper limit to the representational capacity, i.e., the maximal number of distinct patterns. If every input pattern and every output pattern should occur maximally once, the number of distinct input-output pattern pairs is bounded by min 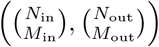.

In general, however, even if the size of the output layer is negligible for the storage capacity under specific circumstances, it plays an important role in a potential additional readout layer, e.g. a classification of the output representations. In [84], it has been shown that an increase in the number of output units increases the performance of a readout unit that performs a binary classification.

## Fast learning

In the context of fast learning, several distinct but related notions are used. Here, it seems useful to explain and compare the concepts of transition probability, learning rate, and convergence time. [64] and [15] are the only two references that discuss the impact of sparseness on learning speed. They build their arguments on analyzing learning rate and convergence, while we focus on the transition probability.

A high transition probability refers to the likelihood that a synaptic weight is increased during learning, while a large learning rate refers to the magnitude of synaptic weight updates (per time unit) during learning. These two are closely related as they both affect changes in weights, and the average weight change in one learning step is proportional to the product of the learning rate and the transition probability. In the present work, connections can only be silent (0) or functional (1). Although the update steps from 0 to 1 or from 1 to 0 could be interpreted as a fixed learning rate, the learning speed is variable and represented by the transition probability *η*.

Another possibility to vary the learning speed would be to use a fixed transition probability while varying the learning rate, e.g., by weight changes that are continuous and within the interval between 0 and 1. We believe that such an alternative model may well yield qualitatively similar results for the effect of sparseness on the optimal learning speed. It would also be interesting to analyze more complex and more realistic synapse models in this context, but all this is beyond the scope of the present work.

In contrast, the term “convergence” and its magnitude do not refer to how many or by how much weights are altered, but it is a system-level outcome. It refers to how quickly a neural network stabilizes or reaches its learning objective, e.g., a minimum in a gradient-descent learning algorithm. While a high learning rate and a high transition probability can both speed up weight updates, fast convergence is achieved by balancing these factors to ensure that the network efficiently settles into an optimal or stable solution. A high learning rate can accelerate updates towards a solution, but it does not necessarily relate to fast convergence. On the contrary, if the update steps are too large, they risk overshooting or instability. [64] argue that sparse representations yield faster convergence because the learning rate can be increased without risking instabilities. In our model, convergence is not evaluated because every pattern is only presented once to do one weight update step, and output patterns are always calculated from the input in one step. It would be interesting to investigate the effect of sparseness on transition probability (and/or learning rate) in combination with convergence speed in a network model that allows for both. Building on the results of this manuscript and [64]’s reasoning, we believe that high sparseness would yield a high optimal transition probability (or learning rate) while converging fast because sparseness allows for large update steps without instability.

## Optimal sparseness values

In terms of the capacity obtained with an optimized transition probability, our study predicts an optimal small but not too small input activation ratio *f*_in_ and an optimal output activation ratio *f*_out_ = 1*/N*_out_ (Fig 3). The optimal *f*_in_ depends on the input size *N*_in_ while the optimal number of active input units *M*_in_ does not (Fig 5). Nevertheless, the optimal *M*_in_ cannot be interpreted as a quantitative prediction of the optimal number of active units because it depends on the retrieval threshold *T*_*S*_ (see Methods).If *T*_*S*_ is increased, i.e., if the retrieval criterion becomes more strict, the optimal *M*_in_ increases (and the corresponding maximum capacity decreases).

Further, the quantification of the optimal learning speed in our model is solely focused on optimizing the capacity of the network.In reality, the brain might favor a trade-off of various properties instead of optimization for one purpose [71]. Even though sparseness can be advantageous, e.g., in terms of energy efficiency [85], storage capacity [86] and downstream classification [84], it might be beneficial to avoid extreme sparseness because it can reduce the representational capacity [71, 87], the information capacity per pattern [88, 89] and the generalization capacity [71, 90] and it is less robust to noise or damage of neuronal structures [71, 87]. Moreover, sparse connectivity, which is also ubiquitous in the brain, seems to yield similar favorable effects as population sparseness and could be optimized in combination with sparse activity [71, 91].

In general, it is thus not reasonable to predict *one* optimal activation ratio or *one* optimal number of active neurons. Optimal (and actual) values are probably different for different brain regions, cell types, tasks, etc. First steps towards more concrete optimal sparseness values could combine experimentally measured values, obtained with modern sophisticated techniques such as Neuropixels recordings, and computational models that take into account the specific properties and presumed role(s) of particular brain regions. [2]

## Supporting information

**S1 Fig. Comparison of optimal transition probability** *η*_**opt**_ **and maximal capacity** 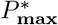 **for different retrieval thresholds** *T*_*S*_.

**S2 Fig. Comparison of optimal transition probability** *η*_**opt**_ **and maximal capacity** 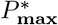 **for different morphological connectivity levels** *c*_*m*_.

**S3 Fig. Comparison of optimal transition probability** *η*_**opt**_ **and maximal capacity** 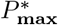 **for different functional connectivity levels** *c*.

**S4 Fig. Comparison of optimal transition probability** *η*_**opt**,*ε*_ **and maximal capacity** 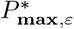 **for different input noise levels** *ε* **during retrieval**.

**S5 Fig. Comparison of distributions of dendritic sums obtained from network simulations and from theory**.

**S6 Fig. Distributions of dendritic sums with noise (***ε* = 0.4).

**S1 Appendix. Analysis of shape of distributions**.

**S2 Appendix. Analytical calculation of the signal quality.**

**S3 Appendix. Monotonicity of capacity for small** *f*_**in**_.

**S4 Appendix. Comparison to other versions of homeostasis.**

**S5 Appendix. Can the output activation ratio be enforced?**

## Acknowledgments

We thank Natalie Schieferstein for valuable comments on the manuscript.

## Notes

### Competing Interest Statement

The authors have declared no competing interest.

### Summary of Updates

Supporting information has been added. Figure S6 revised.

### Funding

- Deutsche Forschungsgemeinschaft: 327654276 SFB 1315

