## Supporting information for "Population sparseness determines strength of Hebbian plasticity for maximal memory lifetime in associative networks"

**S1 Fig.**

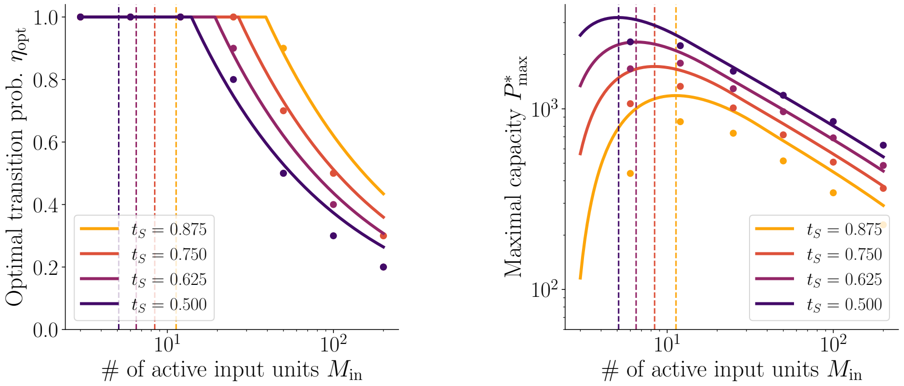


**Fig. S1. Comparison of optimal transition probability $\eta_{\mathbf{opt}}$ and maximal capacity $P^*_{\mathbf{max}}$ for different retrieval thresholds $T_S = t_S H_{\mathbf{avg}}$.** Left: The optimal transition probability $\eta_{\mathrm{opt}}$ increases with increasing $t_S$. Right: The maximal capacity decreases with increasing $t_S$. The number of active input units $M_{\mathrm{in}}$ that yields the largest capacity increases with increasing retrieval ratio $t_S$ (vertical dashed lines). Solid lines show theoretical results obtained from Eq. (20) and Eq. (21), and dots show numerical results. Further parameter values:
$N_{\mathrm{in}} = N_{\mathrm{out}} = 1000, f_{\mathrm{out}} = 0.006, c = 0.2, c_m = 1, N_{\mathrm{avg}} = 200.$

**S2 Fig.**

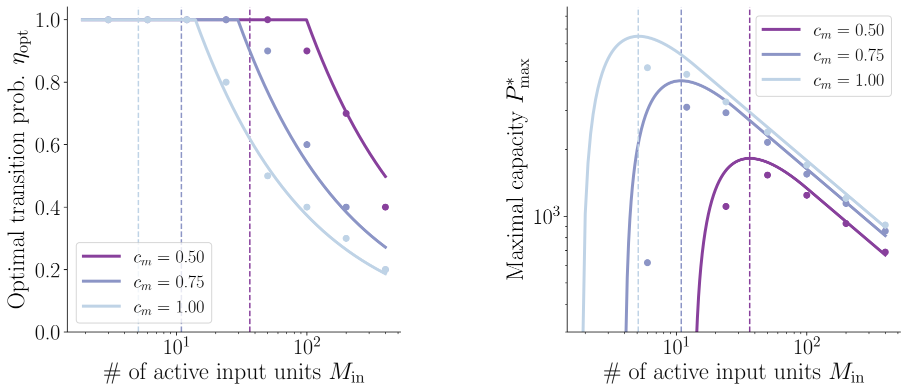


**Fig. S2. Comparison of optimal transition probability $\eta_{\mathrm{opt}}$ and maximal capacity $P^*_{\mathrm{max}}$ for different morphological connectivity levels $c_m$.** Left: The optimal transition probability $\eta_{\mathrm{opt}}$ decreases with increasing $c_m$. Right: The maximal capacity increases with increasing $c_m$. The number of active input units $M_{\mathrm{in}}$ that yields the largest capacity decreases with increasing morphological connectivity $c_m$ (vertical dashed lines). Solid lines show theoretical results obtained from Eq. (20) and Eq. (21), and dots show numerical results. Further parameter values:
$N_{\mathrm{in}} = N_{\mathrm{out}} = 2000, f_{\mathrm{out}} = 0.006, c = 0.2, t_S = 0.5, N_{\mathrm{avg}} = 200.$

**S3 Fig.**

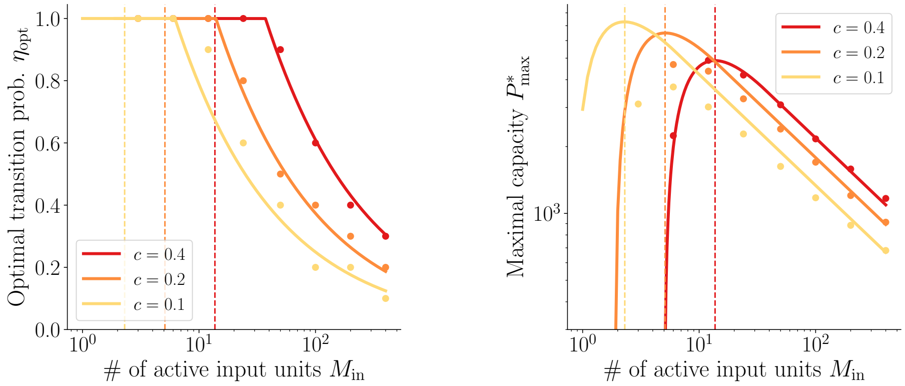


**Fig. S3. Comparison of optimal transition probability $\eta_{\mathbf{opt}}$ and maximal capacity $P^*_{\mathbf{max}}$ for different functional connectivity levels $c$.** Left: The optimal transition probability $\eta_{\mathrm{opt}}$ increases with increasing $c$. Right: The maximal capacity increases with increasing $c$ for large enough $M_{\mathrm{in}}$. The number of active input units $M_{\mathrm{in}}$ that yields the largest capacity increases with increasing functional connectivity $c$ (vertical dashed lines). Solid lines show theoretical results obtained from Eq. (20) and Eq. (21), and dots show numerical results. For small $c$, the analytical approximation strongly overestimates the maximal capacity, especially if only a small number of input units is active because the approximation of the binomial distributions by normal distributions is unsatisfactory for small $M_{\mathrm{in}}c$. Further parameter values: $N_{\mathrm{in}} = N_{\mathrm{out}} = 2000, f_{\mathrm{out}} = 0.006, c_m = 1, t_S = 0.5, N_{\mathrm{avg}} = 200$.

**S4 Fig.**

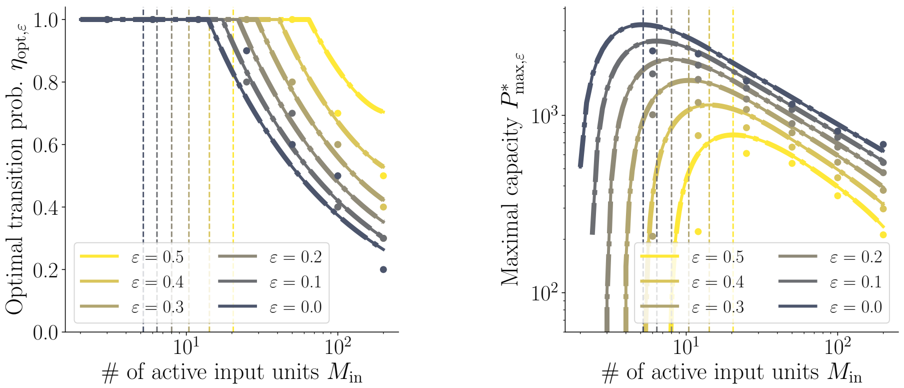


**Fig. S4. Comparison of optimal transition probability $\eta_{\text{opt}}$ and maximal capacity $P^*_{\text{max}}$ for different input noise levels $\varepsilon$ during retrieval.** Left: The optimal transition probability $\eta_{\text{opt},\varepsilon}$ increases with increasing $\varepsilon$. Right: The maximal capacity decreases with increasing $\varepsilon$. The number of active input units $M_{\text{in}}$ that yields the largest capacity increases with increasing noise level $\varepsilon$ (vertical dashed lines). Solid lines show theoretical results obtained from Eq. (20) and Eq. (21), dash-dotted lines show an approximation for small $\varepsilon$ (Eqs. (22) and (26)) and dots show numerical results. Further parameter values:
$N_{\text{in}} = N_{\text{out}} = 1000, f_{\text{out}} = 0.006, c = 0.2, c_m = 1, t_S = 0.5, N_{\text{avg}} = 200$.

#### S5 Fig.

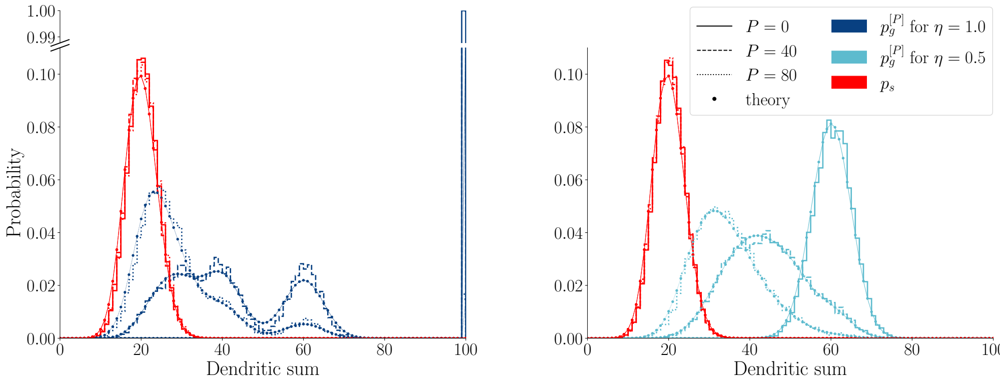


**Fig. S5. Comparison of distributions of dendritic sums obtained from network simulations and from theory.** Histograms show distributions of dendritic sums estimated from numerical simulations, pooled across 200 genuine units and 1800 spurious units and 50 patterns. Different line styles represent different numbers of subsequent patterns ($P = 0$ solid lines, $P = 40$ dashed lines, $P = 80$ dotted lines). Theoretical approximations of the distributions are shown as dots (connected by lines in the respective line styles). Left: $\eta = 1$, right: $\eta = 0.5$. Further parameters: $N_{\text{in}} = 1000, N_{\text{out}} = 2000, f_{\text{in}} = f_{\text{out}} = 0.1, c = 0.2, c_m = 1$.

### S6 Fig.

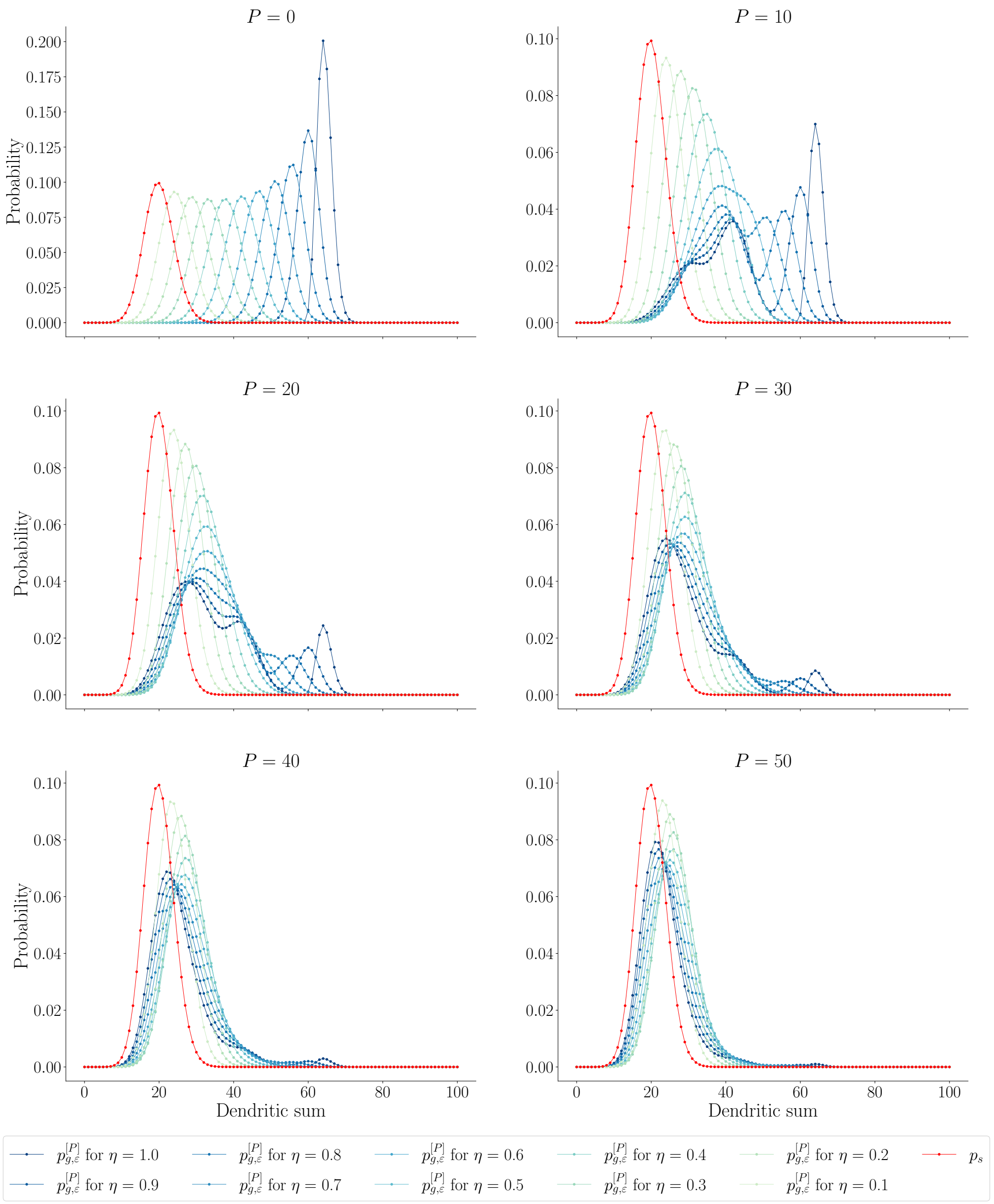


**Fig. S6. Distributions of dendritic sums with noise ($\varepsilon = 0.4$).** Same as Fig 12 but with $\varepsilon = 0.4$. Analytical probability mass functions of the distributions of dendritic sums with noise on the input patterns during retrieval after storing $P = 0, 10, 20, 30, 40, 50$ patterns. Spurious units in red; genuine units in blues, for several transition probabilities $\eta$. Other parameters: $N_{\text{in}} = 1000, f_{\text{in}} = f_{\text{out}} = 0.1, c = 0.2, c_m = 1$.

**S1 Appendix.**

### Analysis of shape of distributions

The distributions of the dendritic sums are not necessarily simple binomial distributions. For many parameter combinations, the distributions are multi-modal and hence the means and standard deviations alone are not useful to characterize the distributions. In the following, we first discuss the multimodality of the distributions using an example. Further, we investigate in which cases the distributions are similar to binomial distributions and in which cases they are not, and we analyze the shape of the distributions in the latter case in more detail. We suggested an approximation using a single binomial distribution, which was necessary to calculate the capacity of the network. This Appendix supports the derivations in the Methods by arguing why the approximation by a single binomial distribution could be satisfactory.

#### Location of bumps — an example

To better understand the location and magnitude of bumps of the distribution of net inputs to a genuine unit, $p_g^{[P]}$, let us first consider one specific example (see Fig S7) with parameters:

$$N_{\text{in}} = 1000, f_{\text{in}} = f_{\text{out}} = 0.1, \eta = 0.8, c = 0.2, c_m = 1, P = 10. \quad (1)$$

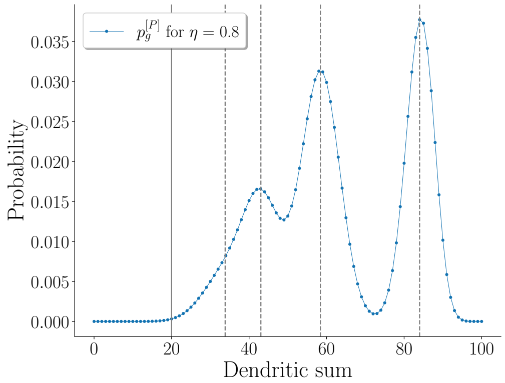


**Fig. S7. Bumps in distribution of dendritic sums for a genuine output unit.** Vertical dashed grey lines correspond to the locations of the peaks of the first four bumps. The center of the right-most peak is at $\rho_g(0)M_{\text{in}} = 84$, the second at $\rho_g(1)M_{\text{in}} = 58.4$, etc. For large values of $u$, the value of $\rho_g(u)$ asymptotically approaches $c$, and then there would be only a single bump at $cM_{\text{in}} = 20$ (solid vertical line). Parameters: $f_{\text{in}} = 0.1, f_{\text{out}} = 0.1, N_{\text{in}} = 1000, c = 0.2, c_m = 1, \eta = 0.8, P = 10$.

For a small $P$, such as $P = 10$ here, a large group of genuine output units are active only during the initial storage of a pattern (here also called the $k$-th pattern) and not in any of the other ten patterns. In fact, the probability that an output unit is active in no subsequent pattern is $\mathcal{B}_{10,0.1}(0) \approx 0.35$, which is the weight of the rightmost binomial distribution $\mathcal{B}_{M_{\text{in}},\rho_g(0)}$. To understand the location of its peak, let us recall that when the $k$-th pattern was learned, the silent genuine-genuine connections became functional with the transition probability $\eta = 0.8$. In addition, $cM_{\text{in}}$ genuine-genuine connections per output unit were already functional due to the initial random connectivity of $cN_{\text{in}}$ connections per output unit. No connections to genuine units were silenced. In total, the genuine units hence receive on average a net input of slightly more than $M_{\text{in}} \cdot \eta = 100 \cdot 0.8 = 80$, namely

$$\rho_g(0) \cdot M_{\text{in}} = (c + (c_m - c)\eta)M_{\text{in}} = (0.2 + (1 - 0.2) \cdot 0.8) \cdot 100 = 84. \tag{2}$$

This is the location of the right-most peak of the distribution.

Some genuine output units are active once across all additional $P$ patterns, say in pattern $k + \kappa$. This decreases their dendritic sum because some of the functional genuine-genuine connections (of the $k$-th pattern) are silenced in order to counteract the connections that became functional for pattern $k + \kappa$ and maintain the functional connectivity level $c$. Note that, nevertheless, a few previously silent genuine-genuine connections of the $k$-th pattern might be made functional if there is an overlap between the active input units of the $k$-th pattern and the active input units of the other pattern $k + \kappa$ for which the respective output unit is active. Overall, the average number of connections that remain functional and receive input, thus contributing to the dendritic sum, is

$$\rho_g(1) \cdot M_{\text{in}} = \left(0.2 + (1 - 0.2) \cdot 0.8 \cdot \left(1 - \frac{0.1 \cdot 0.8 \cdot 1}{0.2}\right)\right) \cdot 100 = 58.4, \tag{3}$$

which is the location of the second peak from the right of the distribution of the dendritic sums in Fig S7. In our example, an output unit is active $f_{\text{out}} \cdot P = 0.1 \cdot 10 = 1$ time on average. The mass that belongs to the second bump from the right is $\mathcal{B}_{10,\,0.1}(1) \approx 0.39$, which is the largest weight because $\mathcal{B}_{10,\,0.1}(1) > B_{10,\,0.1}(u)$ holds, for any $u \neq 1$.

The third peak from the right corresponds to $\rho_g(2)M_{\text{in}} = 43.04$ with a weight of $\mathcal{B}_{10,\,0.1}(2) \approx 0.19$. The remaining peaks are not discriminable and are merging into this third bump.

The width of the peaks can be described by the standard deviation of $\mathcal{B}_{M_{\text{in}},\rho_g(u)}$:

$$\sqrt{M_{\text{in}}\,\rho_g(u)\,(1 - \rho_g(u))}, \tag{4}$$

where $u$ is the output unit usage. Typically, the width first increases with increasing $u$ and then decreases again, but at some point the distance between the peaks becomes smaller than their width and the oscillatory structure fades out. This will be discussed in more detail in the next section.

#### Multimodality of distributions

For spurious units, the probability of a functional connection $\rho_s(u) = c$ is constant as a function of the output unit usage $u$. The distribution of dendritic sums $p_s$ is thus a binomial distribution and always has a single peak (see Eq. (61) in main text). For genuine units, $\rho_g(u)$ is not constant. It decays exponentially with the output unit usage $u$, also approaching the functional connectivity $c$ (see Fig 9). The distribution of dendritic sums $p_g^{[P]}$ is thus a sum of several different binomial probability mass functions (PMFs) $\mathcal{B}_{M_{\text{in}},\rho_g(u)}$, which are weighted by prefactors $\mathcal{B}_{P,f_{\text{out}}}(u)$ (see Eq. (63) in main text).

The multimodality of the distribution of the dendritic sums of genuine units depends on two aspects, which are discussed in this section:

- First, multimodality depends on the weights $\mathcal{B}_{P,f_{\text{out}}}(u)$ in Eq. (63) (black curves in Fig S8). Important is how many $u$ values have a large weight because each large weight adds a binomial distribution $\mathcal{B}_{M_{\text{in}},\rho_g(u)}$ in Eq. (63) to the total distribution of dendritic sums. The more binomials $\mathcal{B}_{M_{\text{in}},\rho_g(u)}$ in Eq. (63) have a large weight $\mathcal{B}_{P,f_{\text{out}}}(u)$, the more distinct bumps could occur in the total distribution.

- Second, multimodality depends on how different the distributions $\mathcal{B}_{M_{\text{in}},\rho_g(u)}$ are, which is determined by the $\rho_g(u)$-values (blue curves in Fig S8). It is therefore important how different the $\rho_g(u)$-values are from each other, i.e., how steep the slope of the blue curves in Fig S8 is, in particular in the range of $u$ values for which $\mathcal{B}_{P,f_{\text{out}}}(u)$ is large (black curves in Fig S8). For example, the more different $\rho_g(u)$ is from $\rho_g(u+1)$, the more the contributions from $\mathcal{B}_{M_{\text{in}},\rho_g(u)}$ and $\mathcal{B}_{M_{\text{in}},\rho_g(u+1)}$ appear as distinct bumps in the total distribution. How well separated the bumps appear is also determined by the widths of the two binomials.

As discussed in the previous subsection for the example in Fig S7, the values of $\rho_g$ determine the locations of the peaks. The first bump (centered at $M_{\text{in}}\rho_g(0)$) represents the probability of the connection being functional if the output unit has never been active across all $P$ additional patterns, the second bump (centered at $M_{\text{in}}\rho_g(1)$) represents the probability of the connection being functional if the output unit has been active once across the pattern set, etc.

Different values $\rho_g(u)$, for $u \in \{1, \dots, P\}$, lead to different contributions to the distribution of dendritic sums. The contributions depend on the corresponding values of $\mathcal{P}(r = u) = \mathcal{B}_{P,\, f_{\text{out}}}(u)$ (cf. Eq. (63)). In general, since only values of $u$ that are less than or equal to $P$ give a probability $\mathcal{B}_{P,\, f_{\text{out}}}(u) > 0$, the number of $\rho_g(u)$-values that contribute to the overall distribution at all is small if $P$ is small. Fig S8 shows the PMF $\mathcal{B}_{P,\, f_{\text{out}}}(u)$ for $f_{\text{out}} = 0.1$ (black, solid line) and the corresponding $\rho_g(u)$ for various transition probabilities $\eta$.

The PMF $\mathcal{B}_{P,\, f_{\text{out}}}$ depends on $P$ and $f_{\text{out}}$ but — for $f_{\text{out}}$ small enough — mostly on the product $P \cdot f_{\text{out}}$. If $f_{\text{out}}$ is decreased by a factor 10 (Fig S8, dark grey dashed line) or 100 (Fig S8, light grey dotted line) and $P$ is increased accordingly by a factor 10 or 100, the PMFs look very similar even though their standard deviations are not exactly the same.

If the contribution of several $\rho_g(u)$-values is large and if these values are not too similar to each other for different $u$, each of them will create a separate bump in the distribution of the dendritic sums. For smaller $u$, the values of $\rho_g(u)$ are more different from each other than for larger $u$. They asymptotically approach $c$ and hence become more and more similar with increasing $u$ (cf. Fig 9). The range of $u$-values for which $\mathcal{B}_{P,\, f_{\text{out}}}(u)$ is large moves to the right with increasing $P$ (Fig S8). For small $P$, several $\rho_g(u)$-values that are very different from each other determine the distribution; this gives rise to clearly distinct bumps. Increasing $P$ leads to the strongly contributing $\rho_g(u)$-values being increasingly similar and making the distribution loose its multi-modality.

The larger $\eta$, the larger the first value $\rho_g(0)$ and hence the more different the first few values are from each other and from the asymptotic value $c$. The exponential decay of $\rho_g(u)$ has a steeper slope the larger $\eta$ is (cf. Fig 9 and Fig S8). If $\eta$ is small we have $\rho_g(0) \gtrsim c$, and thus the decay of $\rho_g(u)$ with increasing $u$ is gentle; and in the distribution of dendritic sum no visually distinct bumps occur. In this case, this distribution resembles a binomial distribution. In contrast, large $\eta$-values can lead to distributions that are less similar to a binomial distribution (at least for small $P$).

How different the $\rho_g(u)$-values must be such that they appear as separate bumps in the distribution and are not perceived as one binomial-like bump depends on the distance between two subsequent peaks compared to the width of the bumps (cf. Fig S9). If the width of the bumps is much larger than the distance between the peaks, they merge into one joint bump. The width of a single bump at $M_{\text{in}}\rho_g(u)$ can be determined by the standard deviation of the corresponding binomial distribution that describes the probability of the connection being functional given a specific output unit

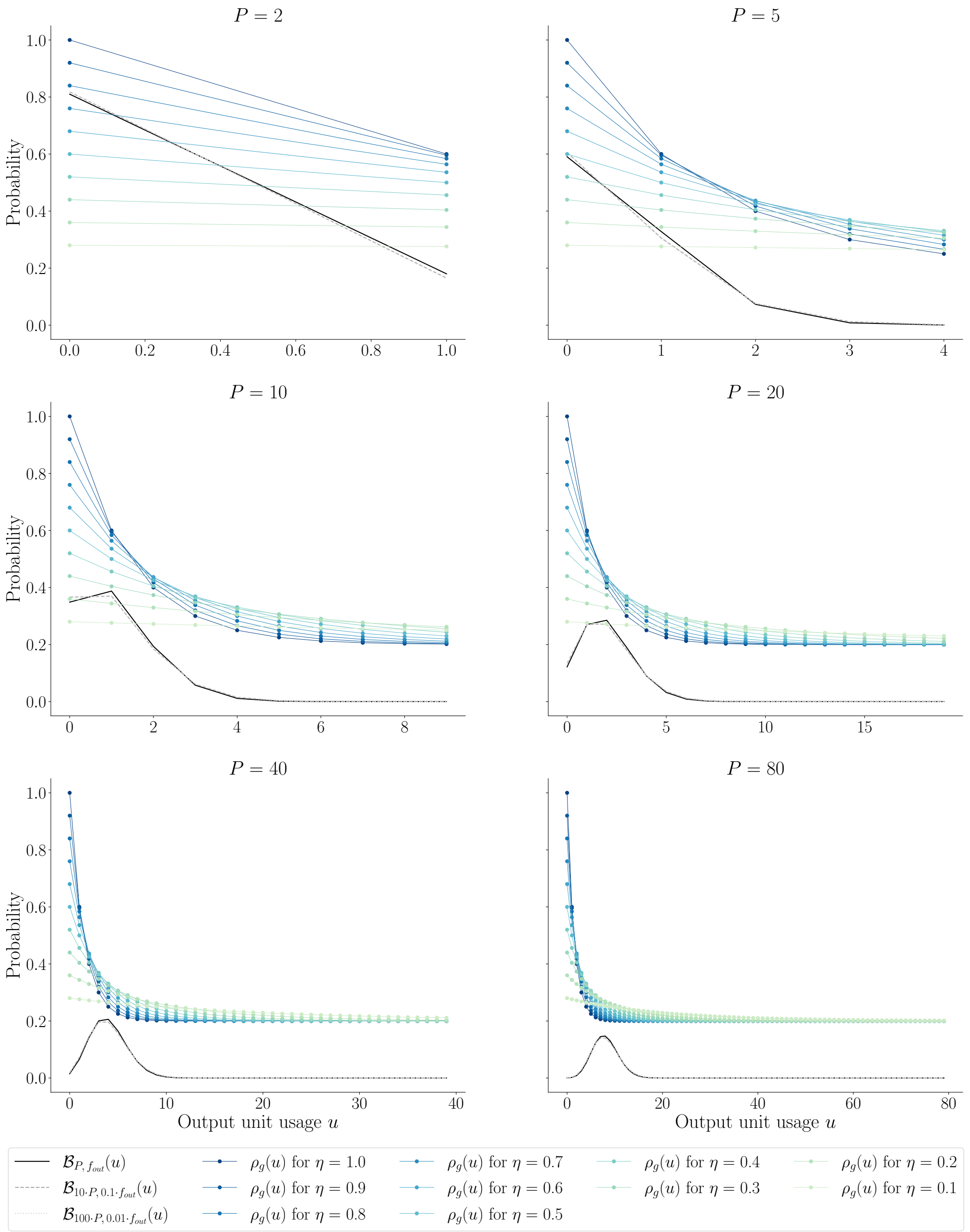


**Fig. S8. Probability of functional connection $\rho_g(u)$ and corresponding weight $\mathcal{B}_{P,\,f_{\mathbf{out}}}(u)$.** The probability $\mathcal{B}_{P,\,f_{\text{out}}}(u)$ that an output unit is on $u$ times across $P(=2,5,10,20,40,80)$ patterns and the corresponding $\rho_g(u)$-values for various $\eta$-values. Black, solid: $f_{\text{out}} = 0.1$; dark grey, dashed: $f_{\text{out}} = 0.01$ and given $P$-value increased by factor 10; light grey, dotted: $f_{\text{out}} = 0.001$ and given $P$-value increased by factor 100. The values of $u$ for which the weight $\mathcal{B}_{P,f_{\text{out}}}(u)$ is large increase with increasing $P$. (Other parameters: $N_{\text{in}} = 1000, f_{\text{in}} = 0.1, f_{\text{out}} = 0.1, c = 0.2, c_m = 1$.)

usage $u$ (cf. Eq. (57)):

$$\sqrt{M_{\text{in}}\rho_g(u)(1-\rho_g(u))} \tag{5}$$
$$= \sqrt{N_{\text{in}} f_{\text{in}} \left[(c_m - c)\eta\left(1 - \frac{f_{\text{in}}\eta c_m}{c}\right)^u + c\right]\left[1 - (c_m - c)\eta\left(1 - \frac{f_{\text{in}}\eta c_m}{c}\right)^u - c\right]}.$$

The distance between the peaks $u$ and $u+1$ is given by

$$
\begin{aligned}
M_{\text{in}}\left|\rho_g(u+1)-\rho_g(u)\right| &= M_{\text{in}}\left(\rho_g(u)-\rho_g(u+1)\right) && (6)\\
&= M_{\text{in}}\left(\rho_g(u)\left(1-1+\frac{f_{\text{in}}\eta c_m}{c}\right)-f_{\text{in}}\eta c_m\right) = N_{\text{in}} f_{in}^2 \eta c_m \left(\rho_g(u)\frac{1}{c}-1\right) && (7)\\
&= N_{\text{in}} f_{\text{in}}^2 \eta \frac{c_m}{c}\left((c_m-c)\eta\left(1-\frac{f_{\text{in}}\eta c_m}{c}\right)^u + c - c\right) && (8)\\
&= N_{\text{in}} f_{\text{in}}^2 \eta^2 \frac{c_m(c_m-c)}{c}\left(1-\frac{f_{\text{in}}\eta c_m}{c}\right)^u . && (9)
\end{aligned}
$$

We define distributions for which the ratio of the distance between the peaks and the standard deviation

$$
\frac{M_{\text{in}}\left|\rho_g(u+1)-\rho_g(u)\right|}{\sqrt{M_{\text{in}}\rho_g(u)(1-\rho_g(u))}} \tag{10}
$$

is smaller than 1 for all $u \geq 0$ as *sufficiently binomial-like.* For $u = 0$ and the special case $\eta = 1$, the ratio of the distance between the peaks and the standard deviation is infinite because the standard deviation is zero since the distribution reduces to $\mathcal{P}(d_g = M_{\text{in}}) = 1$. It can be observed that this ratio decreases with increasing $u$. For $0 < \eta < 1$, we can thus investigate for which parameters the ratio is smaller than 1 at $u = 0$ and imply that it will also be smaller than 1 for any $u > 0$. We have

$$
\begin{aligned}
\frac{M_{\text{in}}\left|\rho_g(1)-\rho_g(0)\right|}{\sqrt{M_{\text{in}}\rho_g(0)(1-\rho_g(0))}} &= \frac{N_{\text{in}} f_{\text{in}}^2 \eta^2 \frac{c_m(c_m-c)}{c}}{\sqrt{N_{\text{in}} f_{\text{in}}\left((c_m-c)\eta+c\right)\left(1-(c_m-c)\eta-c\right)}} && (11)\\
&= \frac{\sqrt{N_{\text{in}} f_{\text{in}}} f_{\text{in}} \eta^2 c_m (c_m-c)}{c\sqrt{\left((c_m-c)\eta+c\right)\left(1-(c_m-c)\eta-c\right)}} && (12)
\end{aligned}
$$

and hence

$$
\begin{aligned}
&\frac{M_{\text{in}}\left|\rho_g(1)-\rho_g(0)\right|}{\sqrt{M_{\text{in}}\rho_g(0)(1-\rho_g(0))}} \ll 1 && (13)\\
\Leftrightarrow\ & N_{\text{in}} f_{\text{in}}^3 \ll \frac{c^2\left((c_m-c)\eta+c\right)\left(1-(c_m-c)\eta-c\right)}{\eta^4 c_m^2 (c_m-c)^2}. && (14)
\end{aligned}
$$

Making further biologically plausible assumptions, namely $c_m = 2c$ and $c \ll 1$, we arrive at the simpler condition

$$
N_{\text{in}} f_{\text{in}}^3 \eta^4 c \ll \frac{1}{4}. \tag{15}
$$

To summarize, the width of the distribution $\mathcal{B}_{P,f_{\text{out}}}$, which determines the number of $\rho_g(u)$-values with a significant contribution, in combination with the normalized distance between the single peaks determine the multimodality of the distribution $p_l^{[P]}$.

In order to analytically derive the capacity of the network in the Methods, it is necessary to approximate the weighted sum of binomial PMFs by a single binomial PMF. This approximation is only justified if the original distribution is *sufficiently binomial-like.* Based on Eq. (15), we find that such an approximation is best for small $N_{\text{in}}$ and, in particular, for small $f_{\text{in}}$ and small $\eta$ as well as small $c$.

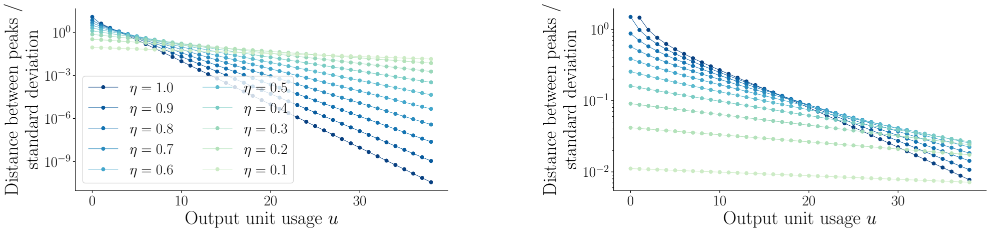


**Fig. S9. Distance between peaks of the bumps divided by the standard deviation of the bumps.** Distance between peaks of the bumps (Eq. (6)) divided by the standard deviation of the bumps (Eq. (5)) for various $\eta$ values for genuine units decreases with output unit usage $u$. Left: $f_{\text{in}} = 0.1$, right: $f_{\text{in}} = 0.025$. The ratio decreases with increasing output unit usage $u$ and with decreasing transition probability $\eta$. (It is $\infty$ for $u = 0$ and $\eta = 1$.) Further parameter values: $N_{\text{in}} = 1000, c = 0.2, c_m = 1$.

**S2 Appendix.**

### Analytical calculation of the signal quality

In order to analytically describe the signal quality $S_P = (1 - \overline{s_P})H_{\text{avg}}$ for a number of subsequent patterns $P$, we have to solve the Signal Quality Equation

$$F_s^{-1}\left(1 - \overline{s_P} f_{\text{out}}\right) = {F_g^{[P]}}^{-1}\left(\overline{s_P}(1 - f_{\text{out}})\right) \tag{16}$$

for $\overline{s_P}$. If we approximate both $p_s$ and $p_g^{[P]}$ by a single normal distribution as outlined in the Methods, the Signal Quality Equation can be replaced by

$$\frac{\bar{\mu}_g^{[P]} - \mu_s}{\sqrt{2}\left(\sigma_s \cdot R_s + \bar{\sigma}_g^{[P]} \cdot R_g\right)} = 1, \tag{17}$$

where

$$R_s := \text{erf}^{-1}\left(1 - 2\overline{s_P} f_{\text{out}}\right), \tag{18}$$

$$R_g := \text{erf}^{-1}\left(2\overline{s_P}(1 - f_{\text{out}}) - 1\right). \tag{19}$$

Note that Eq. (17) is analytically solvable for $\overline{s_P}$ only if $f_{\text{out}} = 0.5$. In this case, we get[1]

$$s_P = \text{erf}\left(\frac{\bar{\mu}_g^{[P]} - \mu_s}{\sqrt{2}\left(\sigma_s + \bar{\sigma}_g^{[P]}\right)}\right) \tag{20}$$

and, thus,

$$S_P = \text{erf}\left(\frac{\bar{\mu}_g^{[P]} - \mu_s}{\sqrt{2}\left(\sigma_s + \bar{\sigma}_g^{[P]}\right)}\right) H_{\text{avg}}, \tag{21}$$

while for other $f_{\text{out}} \neq 0.5$, we need additional assumptions to solve Eq. (17).

---

[1]

$$\frac{\bar{\mu}_g^{[P]} - \mu_s}{\sqrt{2}\left(\sigma_s \cdot \text{erf}^{-1}\left(1 - \overline{s_P}\right) - \bar{\sigma}_g^{[P]} \cdot \text{erf}^{-1}\left(\overline{s_P} - 1\right)\right)} = 1$$

$$\Leftrightarrow \frac{\bar{\mu}_g^{[P]} - \mu_s}{\sqrt{2}\left(\sigma_s \cdot \text{erf}^{-1}\left(1 - \overline{s_P}\right) + \bar{\sigma}_g^{[P]} \cdot \text{erf}^{-1}\left(1 - \overline{s_P}\right)\right)} = 1$$

$$\Leftrightarrow \frac{\bar{\mu}_g^{[P]} - \mu_s}{\sqrt{2}\left(\sigma_s + \bar{\sigma}_g^{[P]}\right)} = \text{erf}^{-1}\left(1 - \overline{s_P}\right)$$

$$\Leftrightarrow 1 - \text{erf}\left(\frac{\bar{\mu}_g^{[P]} - \mu_s}{\sqrt{2}\left(\sigma_s + \bar{\sigma}_g^{[P]}\right)}\right) = \overline{s_P}$$

#### Approximation of $\bar{\sigma}_g^{[P]}$ by $\sigma_s$

For $f_{\text{out}} \neq 0.5$, Eq. (17) is not solvable. Comparing the variances of the genuine and the spurious distributions

$$\bar{\sigma}_g^{[P]^2} = M_{\text{in}} \left[ (c_m - c)\eta \left( 1 - \frac{f_{\text{in}}\eta c_m}{c} \right)^{\lfloor f_{\text{out}}(P+1) \rfloor} + c \right] \tag{22}$$

$$\cdot \left[ 1 - (c_m - c)\eta \left( 1 - \frac{f_{\text{in}}\eta c_m}{c} \right)^{\lfloor f_{\text{out}}(P+1) \rfloor} - c \right], \tag{23}$$

$${\sigma_s}^2 = M_{\text{in}} c(1 - c), \tag{24}$$

respectively, yields

$$\bar{\sigma}_g^{[P]^2} - \sigma_s^2 = M_{\text{in}}(c_m - c)\eta \left( 1 - \frac{f_{\text{in}}\eta c_m}{c} \right)^{\lfloor f_{\text{out}}(P+1) \rfloor} \tag{25}$$

$$\cdot \left[ 1 - 2c - (c_m - c)\eta \left( 1 - \frac{f_{\text{in}}\eta c_m}{c} \right)^{\lfloor f_{\text{out}}(P+1) \rfloor} \right] \tag{26}$$

For $P \to \infty$, we have $\bar{\sigma}_g^{[P]} \to \sigma_s$ (Fig S10). The case of large $P$ is particularly interesting for calculating the memory lifetime. We thus assume that, when $S_P$ reaches the retrieval threshold, $P$ is large enough for the assumption $\bar{\sigma}_g^{[P]} \approx \sigma_s$ to be reasonable. Eq. (17) now reads as

$$\frac{\bar{\mu}_g^{[P]} - \mu_s}{\sqrt{2}\sigma_s} = R_s + R_g. \tag{27}$$

Note that for large $\eta$ and small $P$, this assumption is less accurate (e.g., $\bar{\sigma}_g^{[0]} \to 0$ for $\eta \to 1$, while $\sigma_s$ does not depend on $\eta$) (Fig S10).

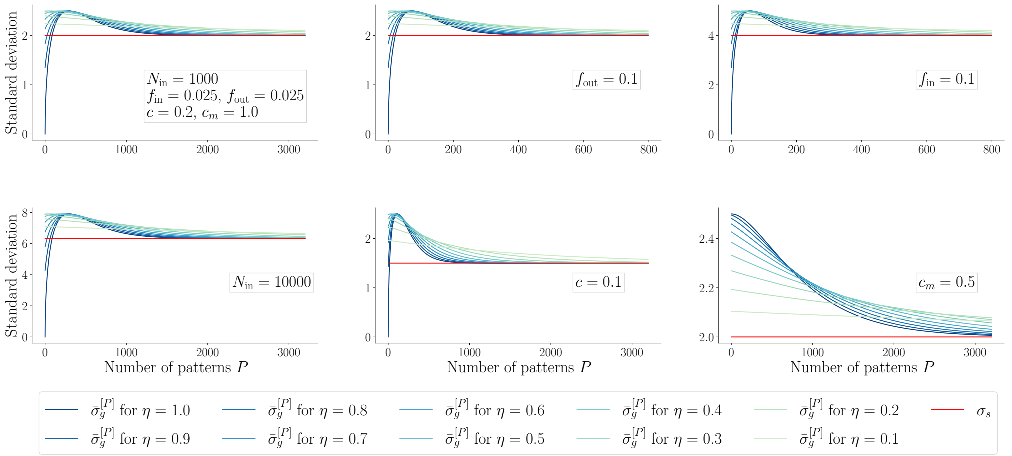


**Fig. S10. Comparison of $\bar{\sigma}_g^{[P]}$ and $\sigma_s$.** Comparison of the standard deviations of the genuine (blue lines) and the spurious (red lines) distributions of dendritic sums for various parameter combinations. Top left: default parameters $N_{\text{in}} = 1000, f_{\text{in}} = 0.025, f_{\text{out}} = 0.025, c = 0.2, c_m = 1$; top center: $f_{\text{out}} = 0.1$; top right: $f_{\text{in}} = 0.1$; bottom left: $N_{\text{in}} = 10000$; bottom center: $c = 0.1$; bottom right: $c_m = 0.5$.

#### Approximation of the error function

In order to solve Eq. (27) for $\overline{s_P}$, we now approximate the error function by

$$\mathrm{erf}(x) \approx \tanh\left(\frac{x\pi}{\sqrt{6}}\right) \tag{28}$$

and use $\mathrm{arctanh}(x) = \frac{1}{2}\ln\left(\frac{1+x}{1-x}\right)$. This yields[2]

$$\frac{\bar{\mu}_g^{[P]} - \mu_s}{\sqrt{2}\sigma_s} = \frac{\sqrt{6}}{\pi}\left[\mathrm{arctanh}\left(1 - 2\overline{s_P} f_{\mathrm{out}}\right) - \mathrm{arctanh}\left(2\overline{s_P}(1 - f_{\mathrm{out}}) - 1\right)\right] \tag{29}$$

$$\Leftrightarrow \overline{s_P} = \frac{-1 + \sqrt{1 + 4 f_{\mathrm{out}}(1 - f_{\mathrm{out}})\left(\exp\left(\frac{\pi}{\sqrt{3}}\frac{\bar{\mu}_g^{[P]} - \mu_s}{\sigma_s}\right) - 1\right)}}{2 f_{\mathrm{out}}(1 - f_{\mathrm{out}})\left(\exp\left(\frac{\pi}{\sqrt{3}}\frac{\bar{\mu}_g^{[P]} - \mu_s}{\sigma_s}\right) - 1\right)}. \tag{30}$$

Note that the more accurate approximation of the error function employed in the Methods (Eq. (153) in the main text) cannot be used here because it does not allow us to solve Eq. (27) for $\overline{s_P}$.

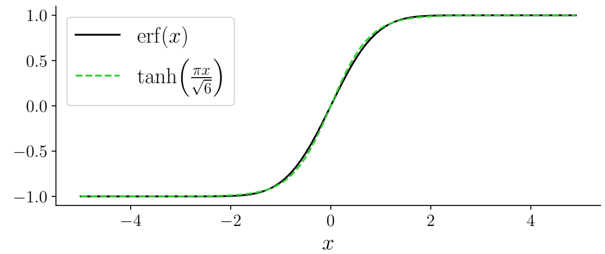


**Fig. S11. Approximation of error function by a hyperbolic tangent.** The maximal absolute error is 0.0453, the maximal relative error is 0.1366.

The signal quality $S_P = s_P H_{\mathrm{avg}} = (1 - \overline{s_P}) H_{\mathrm{avg}}$ with $\overline{s_P}$ from Eq. (30) as a function of the number of patterns $P$ is shown in Fig S12) (colored dashed lines). The analytical

[2]

$$\begin{aligned}
\frac{\bar{\mu}_g^{[P]} - \mu_s}{\sqrt{2}\sigma_s} &= \frac{\sqrt{6}}{\pi}\left[\mathrm{arctanh}\left(1 - 2\overline{s_P} f_{\mathrm{out}}\right) - \mathrm{arctanh}\left(2\overline{s_P}(1 - f_{\mathrm{out}}) - 1\right)\right] \\
\Leftrightarrow \frac{\bar{\mu}_g^{[P]} - \mu_s}{\sqrt{2}\sigma_s} &= \frac{\sqrt{6}}{2\pi}\left[\ln\left(\frac{2 - 2\overline{s_P} f_{\mathrm{out}}}{2\overline{s_P} f_{\mathrm{out}}}\right) - \ln\left(\frac{2\overline{s_P}(1 - f_{\mathrm{out}})}{2 - 2\overline{s_P}(1 - f_{\mathrm{out}})}\right)\right] \\
\Leftrightarrow \frac{\bar{\mu}_g^{[P]} - \mu_s}{\sqrt{2}\sigma_s} &= \frac{\sqrt{6}}{2\pi}\left[\ln\left(\frac{(1 - \overline{s_P} f_{\mathrm{out}})(1 - \overline{s_P}(1 - f_{\mathrm{out}}))}{\overline{s_P} f_{\mathrm{out}} \overline{s_P}(1 - f_{\mathrm{out}})}\right)\right] \\
\Leftrightarrow \exp\left(\frac{\pi}{\sqrt{3}}\frac{\bar{\mu}_g^{[P]} - \mu_s}{\sigma_s}\right) &= \frac{1 - \overline{s_P} + \overline{s_P}^2 f_{\mathrm{out}}(1 - f_{\mathrm{out}})}{\overline{s_P}^2 f_{\mathrm{out}}(1 - f_{\mathrm{out}})} \\
\Leftrightarrow 0 &= \overline{s_P}^2 \cdot f_{\mathrm{out}}(1 - f_{\mathrm{out}})\left(\exp\left(\frac{\pi}{\sqrt{3}}\frac{\bar{\mu}_g^{[P]} - \mu_s}{\sigma_s}\right) - 1\right) + \overline{s_P} - 1 \\
\Leftrightarrow \overline{s_P} &= \frac{-1 + \sqrt{1 + 4 f_{\mathrm{out}}(1 - f_{\mathrm{out}})\left(\exp\left(\frac{\pi}{\sqrt{3}}\frac{\bar{\mu}_g^{[P]} - \mu_s}{\sigma_s}\right) - 1\right)}}{2 f_{\mathrm{out}}(1 - f_{\mathrm{out}})\left(\exp\left(\frac{\pi}{\sqrt{3}}\frac{\bar{\mu}_g^{[P]} - \mu_s}{\sigma_s}\right) - 1\right)}
\end{aligned}$$

As $\bar{\mu}_g^{[P]} > \mu_s$, the denominator is positive. The numerator also has to be positive to achieve $\overline{s_P} \in [0, 1]$. Since the term inside the square root is greater than one, the solution of the quadratic equation for $\overline{s_P}$ with the positive sign before the square root yields a valid $\overline{s_P}$. Using the negative sign before the square root in the numerator, which would of course also yield a solution of the second to last equation, would give a negative numerator and by that a negative $\overline{s_P}$ and should thus be neglected.

estimate captures the trend of the lines and, most importantly, their relation to each other if different transition probabilities are compared. It matches the results of network simulations in the sense that the signal quality $S_P$ for larger transition probability $\eta$ is initially higher but decays more quickly with the number of patterns $P$. This yields different optimal transition probabilities depending on the input and output activation ratios $f_{\text{in}}$ and $f_{\text{out}}$. The semi-analytical strategy presented in the Methods yields a better approximation of the results from network simulations than the signal quality obtained from Eq. (30) (compare Fig 17 to Fig S12). This benefit of the semi-analytical approximation compared to the fully analytical approximation is particularly prominent for large $\eta$, which is due to the fact that both the approximation of $p_g^{[P]}$ by a single binomial distribution and the approximation of $\bar{\sigma}_g^{[P]}$ by $\sigma_s$ used in the derivation of Eq. (30) (see Methods) are less accurate for large $\eta$ than for small $\eta$.

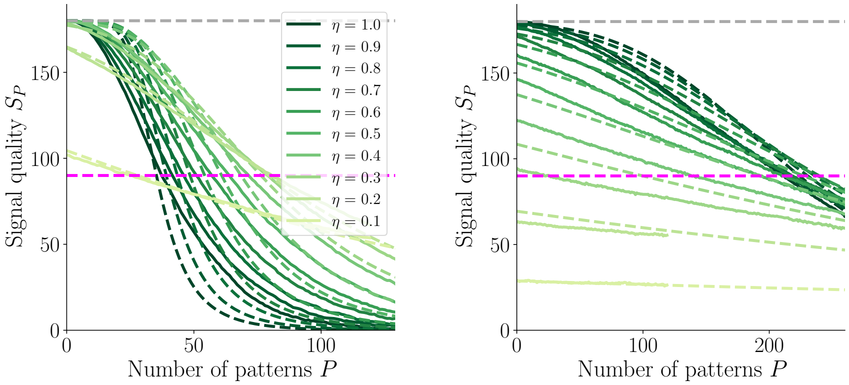


**Fig. S12. Comparison of signal quality obtained from network simulations and from analytical approximation.** Signal quality $S_P$ decays with number $P$ of subsequently learned patterns for various values of transition probability $\eta$. Solid lines: numerical network simulations (see Methods); dashed lines: analytical estimate (based on Eq. (30)); dashed grey line: maximal signal quality, which is the average Hamming distance between two random $f_{\text{out}}$-sparse patterns; dashed magenta line: retrieval threshold. Left: $f_{\text{in}} = 0.1$, right: $f_{\text{in}} = 0.012$. Other parameters: $N_{\text{in}} = N_{\text{out}} = 1000, f_{\text{out}} = 0.1, c_m = 1, c = 0.2$.

#### Signal quality with noisy input patterns

In this subsection, we evaluate the change of the signal quality when we apply noise to input patterns during retrieval (see Methods). The case without noise was already approximated in Eq. (30). If we include noise in this equation, the only term that changes there is the mean of the genuine distribution, which can be approximated by

$$\bar{\mu}_{g,\varepsilon}^{[P]} \approx m_g \rho_g(\lfloor f_{\text{out}}(P+1) \rfloor) + m_s \rho_n(\lfloor f_{\text{out}}(P+1) \rfloor) \tag{31}$$

$$= m_g \left( (c_m - c)\eta \left( 1 - \frac{f_{\text{in}} \eta c_m}{c} \right)^{\lfloor f_{\text{out}}(P+1) \rfloor} + c \right) \tag{32}$$

$$+ m_s \left( \frac{-f_{\text{in}} \eta (c_m - c)}{1 - f_{\text{in}}} \left( 1 - \frac{f_{\text{in}} \eta c_m}{c} \right)^{\lfloor f_{\text{out}}(P+1) \rfloor} + c \right). \tag{33}$$

The additional noise shifts the signal quality to lower values (compare Fig 6A top to Fig 6A bottom). Fig S13 compares the initial signal quality as a function of the transition probability $\eta$ for several noise levels $\epsilon$ and, in addition, it shows the impact of other parameters on the initial signal quality.

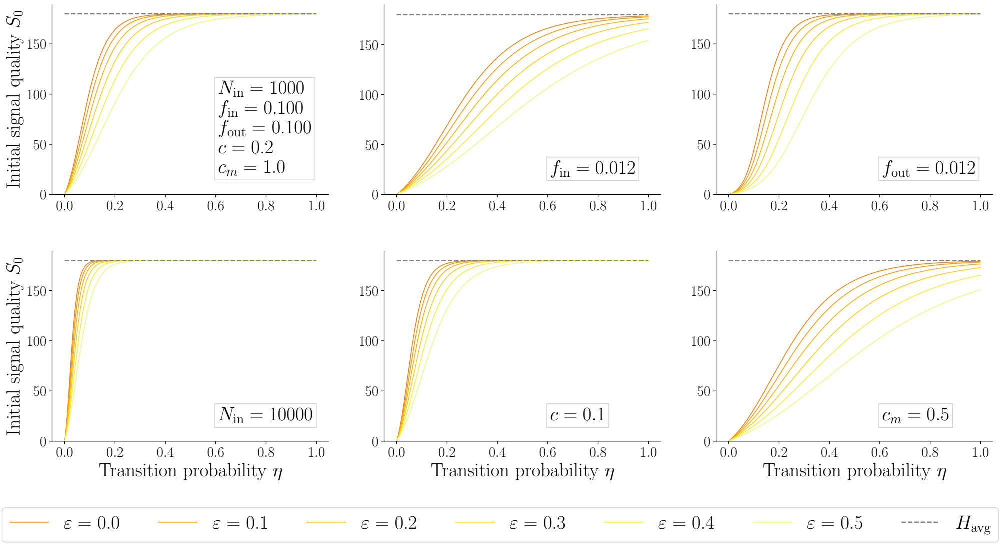


**Fig. S13. Effect of noise on the initial signal quality.** Comparison of initial signal quality based on Eq. (30) with $\mu_{g,\varepsilon}$ (Eq. (31)) for $P = 0$ as a function of the transition probability $\eta$ for various noise levels $\varepsilon$. The initial signal quality is monotonically increasing as a function of the transition probability $\eta$. It approaches its maximal value $H_{\text{avg}}$ more slowly if the noise level $\varepsilon$ is increased. Default parameters (top left): $N_{\text{in}} = N_{\text{out}} = 1000, f_{\text{in}} = f_{\text{out}} = 0.1, c = 0.2, c_m = 1$. Top row: center – $f_{\text{in}} = 0.012$, right – $f_{\text{out}} = 0.012$. Bottom row: left – $N_{\text{in}} = 10^4$, center – $c = 0.1$, right – $c_m = 0.5$.

Given the signal quality without noise $S_P$ (e.g. from numerical simulations), the corresponding signal quality with noisy input patterns during retrieval $S_{P,\varepsilon}$ can be estimated as

$$S_{P,\varepsilon} \approx S_P \cdot \frac{s_{P,\varepsilon} H_{\text{avg}}}{s_P H_{\text{avg}}} = S_P \cdot \frac{1 - \overline{s_{P,\varepsilon}}}{1 - \overline{s_P}}. \tag{34}$$

The factor that relates the signal quality without noise $S_P$ to the signal quality with noise $S_{P,\varepsilon}$ is thus given by

$$\frac{S_{P,\varepsilon}}{S_P} = \frac{1 - \frac{-1+\sqrt{1+4f_{\text{out}}(1-f_{\text{out}})\left(\exp\left(\frac{\pi}{\sqrt{3}}\frac{\bar{\mu}_{g,\varepsilon}^{[P]}-\mu_s}{\sigma_s}\right)-1\right)}}{2f_{\text{out}}(1-f_{\text{out}})\left(\exp\left(\frac{\pi}{\sqrt{3}}\frac{\bar{\mu}_{g,\varepsilon}^{[P]}-\mu_s}{\sigma_s}\right)-1\right)}}{1 - \frac{-1+\sqrt{1+4f_{\text{out}}(1-f_{\text{out}})\left(\exp\left(\frac{\pi}{\sqrt{3}}\frac{\bar{\mu}_{g}^{[P]}-\mu_s}{\sigma_s}\right)-1\right)}}{2f_{\text{out}}(1-f_{\text{out}})\left(\exp\left(\frac{\pi}{\sqrt{3}}\frac{\bar{\mu}_{g}^{[P]}-\mu_s}{\sigma_s}\right)-1\right)}}. \tag{35}$$

Naturally, this factor is smaller for larger $\varepsilon$. For small transition probabilities, it is almost constant as a function of $P$ (see solid lines with slope close to zero in Fig S14); however, for large transition probabilities, the factor is a non-monotonous function of $P$ (dotted lines in Fig S14), and thus the maximal capacity and the optimal transition probability could exhibit a complex dependence on noise. Note that this analytical factor $S_{P,\varepsilon}/S_P$ is independent of $H_{\text{avg}}$ and thus independent of $N_{\text{out}}$.

Fig S15 compares the signal quality $S_{P,\varepsilon}$ obtained from numerical simulations with noise $\varepsilon = 0.1$ (solid lines) to its approximation where the numerically obtained signal quality $S_P$ without noise is scaled by the analytical factor (dotted lines; see Eq. (35)).

The factor Eq. (35) allows for a good approximation of the signal quality with noise based on the signal quality without noise.

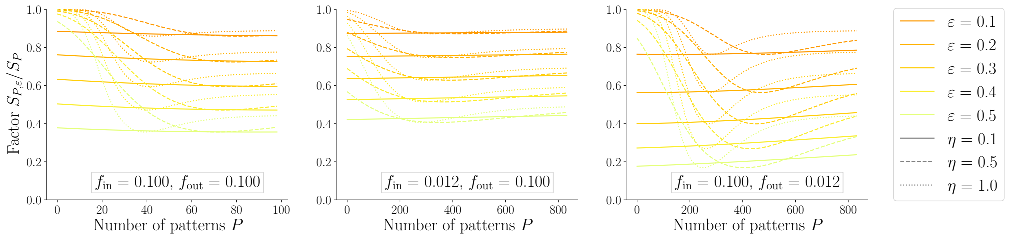


**Fig. S14. Factor relating $S_P$ to $S_{P,\varepsilon}$.** Non-constant factor $S_{P,\varepsilon}/S_P$ that relates the signal quality without noise to the signal quality with noise level $\varepsilon$ (cf. Eq. (35)) as a function of the number of patterns $P$ for various noise levels $\varepsilon = 0.1, 0.2, 0.3, 0.4, 0.5$ (colors) and various transition probabilities $\eta = 0.1, 0.5, 1.0$ (solid, dashed and dotted). Default parameters (left): $N_{\text{in}} = N_{\text{out}} = 1000, f_{\text{in}} = f_{\text{out}} = 0.1, c = 0.2, c_m = 1$. Center: $f_{\text{in}} = 0.012$. Right: $f_{\text{out}} = 0.012$.

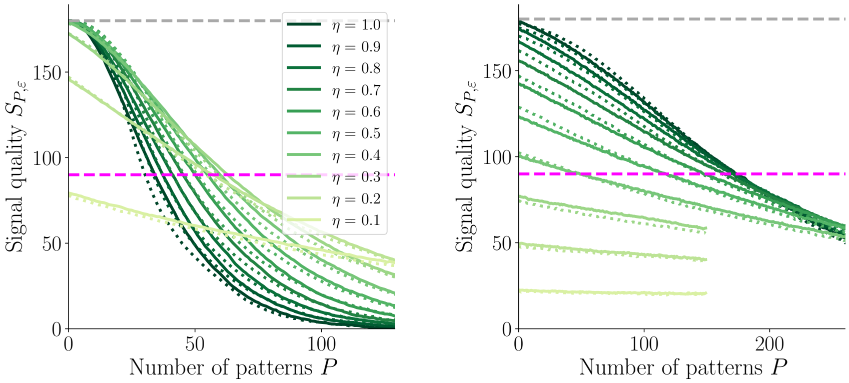


**Fig. S15. Approximation of $S_{P,\varepsilon}$ by scaling $S_P$.** Signal quality decays with number of subsequently learned patterns $P$ for various values of transition probability $\eta$. Noise on input pattern during retrieval: $\varepsilon = 0.2$. The approximation of the signal quality $S_{P,\varepsilon}$ by Eq. (34), where $S_P$ (without noise, $\varepsilon = 0$) is obtained from numerical simulations, fits the signal quality $S_{P,\varepsilon}$ obtained directly from numerical simulations with noise well. Solid lines - numerical simulations for $S_{P,\varepsilon}$ (see Methods); dotted lines - approximation of the signal quality $S_{P,\varepsilon}$ by Eq. (34) (with the factor Eq. (35)), where $S_P$ (without noise, $\varepsilon = 0$) is obtained from numerical simulations (see Methods); dashed grey line - maximal signal quality which is the average Hamming distance between two random $f_{\text{out}}$-sparse patterns; dashed blue line - retrieval threshold. Left: $f_{\text{in}} = 0.1$, right: $f_{\text{in}} = 0.012$. (Other parameters: $N_{\text{in}} = N_{\text{out}} = 1000, f_{\text{out}} = 0.1, c_m = 1, c = 0.2$.)

**S3 Appendix.**

### Monotonicity of capacity for small $f_{\text{in}}$

In this section, we show that the capacity as a function of the input activation ratio $f_{\text{in}}$ is monotonically increasing for $f_{\text{in}}$ close to zero. The generalized sensitivity index

$$d'_R = \frac{\bar{\mu}_g^{[P]} - \mu_s}{\sqrt{2}\left(\sigma_s R_s + \bar{\sigma}_g^{[P]} R_g\right)} \tag{36}$$

(introduced in Eq. (133)) is decreasing with the number of patterns $P$. The memory capacity of the network is the number of patterns $P$ for which the equation

$$d'_R = 1 \tag{37}$$

is fulfilled. In the following, we show that, for fixed $P$ and small $f_{\text{in}}$, $d'_R$ is monotonically increasing as a function of $f_{\text{in}}$. Hence, the smaller $f_{\text{in}}$, the smaller the $P$ for which Eq. (37) is fulfilled and the smaller the capacity.

For small values of $f_{\text{in}}$, we use the Taylor expansion of $\rho_g(u_{\text{mod}})$, where $u_{\text{mod}} = \lfloor f_{\text{out}}(P+1) \rfloor$, as a function of $f_{\text{in}}$ at $f_{\text{in}} = 0$

$$\begin{aligned} \rho_g(u_{\text{mod}}) &= (c_m - c)\eta \left(1 - f_{\text{in}} \frac{\eta c_m}{c}\right)^{u_{\text{mod}}} + c && (38) \\ &= (c_m - c)\eta \left(1 - f_{\text{in}} \frac{\eta c_m u_{\text{mod}}}{c} + \mathcal{O}(f_{\text{in}}^2)\right) + c && (39) \end{aligned}$$

and obtain the Taylor expanded versions of Eqs. (136)-(139) (see Methods)

$$\begin{aligned} \mu_s &= f_{\text{in}} N_{\text{in}} c && (40) \\ \sigma_s &= \sqrt{f_{\text{in}} N_{\text{in}} c(1-c)} && (41) \\ \bar{\mu}_g^{[P]} &= f_{\text{in}} N_{\text{in}} \left[(c_m - c)\eta \left(1 - f_{\text{in}} \frac{\eta c_m u_{\text{mod}}}{c} + \mathcal{O}(f_{\text{in}}^2)\right) + c\right] && (42) \\ \bar{\sigma}_g^{[P]} &= \sqrt{\begin{aligned} & f_{\text{in}} N_{\text{in}} \left[(c_m - c)\eta \left(1 - f_{\text{in}} \frac{\eta c_m u_{\text{mod}}}{c} + \mathcal{O}(f_{\text{in}}^2)\right) + c\right] \\ & \cdot \left[1 - (c_m - c)\eta \left(1 - f_{\text{in}} \frac{\eta c_m u_{\text{mod}}}{c} + \mathcal{O}(f_{\text{in}}^2)\right) - c\right]. \end{aligned}} && (43) \end{aligned}$$

In order to show that $d'_R$ is monotonically increasing with $f_{\text{in}}$, we show that

$$f_1 < f_2 \Rightarrow d'_R(f_1) \leq d'_R(f_2). \tag{44}$$

We have

$$\begin{aligned} & \frac{\bar{\mu}_g^{[P]} - \mu_s}{\sqrt{2}\left(\sigma_s R_s + \bar{\sigma}_g^{[P]} R_g\right)}(f_1) \leq \frac{\bar{\mu}_g^{[P]} - \mu_s}{\sqrt{2}\left(\sigma_s R_s + \bar{\sigma}_g^{[P]} R_g\right)}(f_2) && (45) \\ & \Leftrightarrow \sqrt{f_1}\left(1 - f_1 \frac{\eta c_m u_{\text{mod}}}{c} + \mathcal{O}(f_1^2)\right) \cdot \left[R_s \sqrt{c(1-c)} + R_g \sqrt{a} \sqrt{1 + f_2 \frac{b}{a} + \mathcal{O}(f_2^2)}\right] && (46) \\ & \leq \sqrt{f_2}\left(1 - f_2 \frac{\eta c_m u_{\text{mod}}}{c} + \mathcal{O}(f_2^2)\right) \cdot \left[R_s \sqrt{c(1-c)} + R_g \sqrt{a} \sqrt{1 + f_1 \frac{b}{a} + \mathcal{O}(f_1^2)}\right] \end{aligned}$$

with $a := c(1-c) + (c_m - c)\eta(1 - 2c - (c_m - c)\eta)$ and $b := (c_m - c)\eta \frac{\eta c_m u_{\text{mod}}}{c}(2(c_m - c)\eta - 1 + 2c)$. Note that $a > 0$ only if $\eta$ and $c_m$ are not both 1. We exclude the case $c_m = 1$, $\eta = 1$ here. However, evaluating Eq. (152) for small $f_{\text{in}}$ values suggests that the monotonicity of the capacity as a function of $f_{\text{in}}$ is maintained also for $c_m = 1$, $\eta = 1$.

We know that $\bar{\mu}_g^{[P]} - \mu_s \geq 0$ and since we are interested in $d'_R \geq 0$, we can further assume that $\sigma_s R_s + \bar{\sigma}_g^{[P]} R_g \geq 0$. Both sides of the inequality are thus positive and we can square them to obtain

$$f_1 \left(1 - 2f_1 \frac{\eta c_m u_{\text{mod}}}{c} + \mathcal{O}(f_1^2)\right) \tag{47}$$

$$\cdot \left[R_s^2 c(1-c) + 2R_s\sqrt{c(1-c)}R_g\sqrt{a}\sqrt{1 + f_2\frac{b}{a} + \mathcal{O}(f_2^2)} + R_g^2 a\left(1 + f_2\frac{b}{a} + \mathcal{O}(f_2^2)\right)\right]$$

$$\leq f_2 \left(1 - 2f_2 \frac{\eta c_m u_{\text{mod}}}{c} + \mathcal{O}(f_2^2)\right) \tag{48}$$

$$\cdot \left[R_s^2 c(1-c) + 2R_s\sqrt{c(1-c)}R_g\sqrt{a}\sqrt{1 + f_1\frac{b}{a} + \mathcal{O}(f_1^2)} + R_g^2 a\left(1 + f_1\frac{b}{a} + \mathcal{O}(f_2^2)\right)\right].$$

We now use the Taylor expansion

$$\sqrt{1 + x\alpha} = 1 + x\frac{\alpha}{2} + \mathcal{O}(x^2) \tag{49}$$

at $x = 0$ and obtain

$$f_1 \left(1 - 2f_1 \frac{\eta c_m u_{\text{mod}}}{c} + \mathcal{O}(f_1^2)\right) \tag{50}$$

$$\cdot \left[\left(R_s\sqrt{c(1-c)} + R_g\sqrt{a}\right)^2 + f_2\frac{b}{\sqrt{a}}R_g\left(R_g\sqrt{a} + R_s\sqrt{c(1-c)}\right) + \mathcal{O}(f_2^2)\right]$$

$$\leq f_2 \left(1 - 2f_2 \frac{\eta c_m u_{\text{mod}}}{c} + \mathcal{O}(f_2^2)\right) \tag{51}$$

$$\cdot \left[\left(R_s\sqrt{c(1-c)} + R_g\sqrt{a}\right)^2 + f_1\frac{b}{\sqrt{a}}R_g\left(R_g\sqrt{a} + R_s\sqrt{c(1-c)}\right) + \mathcal{O}(f_1^2)\right].$$

Keeping only the terms up to the first order yields

$$f_1\left(R_s\sqrt{c(1-c)} + R_g\sqrt{a}\right)^2 \leq f_2\left(R_s\sqrt{c(1-c)} + R_g\sqrt{a}\right)^2. \tag{52}$$

Since $\left(R_s\sqrt{c(1-c)} + R_g\sqrt{a}\right)^2 \geq 0$, this inequality is true for $f_1 < f_2$ and we have thus shown that the capacity is a monotonically increasing function of $f_{\text{in}}$ for values of $f_{\text{in}}$ close to zero.

**S4 Appendix.**

### Comparison to other versions of homeostasis

The homeostatic mechanism in this manuscript normalizes the number of functional connections per output unit, i.e., the in-degree, by silencing connections that originate from an inactive input unit. In the following, we compare this method to two other versions of synaptic homeostasis.

Instead of normalizing the in-degree, the number of functional connections per input unit could be preserved. This version simply inverses the roles of input and output units in the homeostasis step: Among the genuine-spurious connections, we randomly silence as many as necessary to maintain the connectivity $c$ per input unit. In this case, we also normalize the morphological connectivity $c_m$ per input unit and not per output unit. Fig S16A and B show the maximal capacity $P^*_{\text{max}}$ and the optimal transition probability $\eta_{\text{opt}}$ obtained from numerical simulations with this method.

Alternatively, the functional connectivity $c$ (and the morphological connectivity $c_m$) could only be maintained as an average across the whole network. In this instance, the appropriate number of randomly chosen spurious-spurious connections are silenced. The maximal capacity $P^*_{\text{max}}$ and the optimal transition probability $\eta_{\text{opt}}$ for random deactivation are depicted in Fig S16C and D.

In both cases, the results remain qualitatively unchanged compared to the results presented in the manuscript: The maximal capacity $P^*_{\text{max}}$ decreases as a function of the output activation ratio $f_{\text{out}}$ but depends non-monotonically on the input activation ratio $f_{\text{in}}$. The optimal transition probability $\eta_{\text{opt}}$ decreases with both $f_{\text{in}}$ and $f_{\text{out}}$ but the effect due to $f_{\text{in}}$ is stronger (compare Fig S16 to Fig 3D,E).

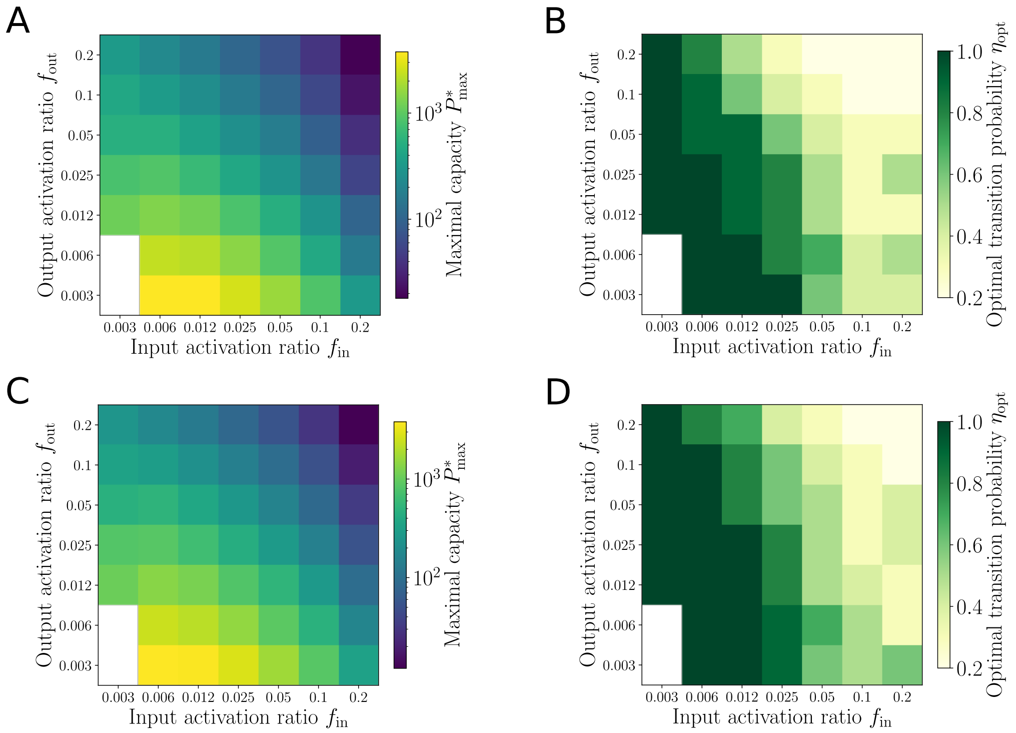


**Fig. S16. Maximal capacity and optimal transition probability with other versions of homeostasis.** The effect of $f_{\text{in}}$ and $f_{\text{out}}$ on the memory capacity $P^*_{\max}$ and on the optimal transition $\eta_{\text{opt}}$ is qualitatively preserved with other homeostasis mechanisms. In (**A**)-(**B**), the functional connectivity $c$ is preserved per input unit. In (**C**)-(**D**), $c$ is maintained across the whole network. In both (**A**)-(**B**) and (**C**)-(**D**), the memory capacity $P^*_{\max}$ monotonically decreases with $f_{\text{out}}$ whereas $P^*_{\max}$ depends non-monotonically on $f_{\text{in}}$. The optimal transition probability $\eta_{\text{opt}}$ decreases with increasing $f_{\text{in}}$ and $f_{\text{out}}$. Compare (**A**) and (**C**) to Fig 3D as well as (**B**) and (**D**) to Fig 3E. In (**A**)-(**D**), $N_{\text{in}} = N_{\text{out}} = 1000, c = 0.2, c_m = 1, t_S = 0.5, N_{\text{avg}} = 200$.

**S5 Appendix.**

### Can the output activation ratio be enforced?

During retrieval, since the distributions of dendritic sums are discrete distributions, it is not necessarily possible (or rather usually impossible) to choose an activation threshold such that exactly a number $M_{\text{out}}$ of output units is activated and exactly a number $N_{\text{out}} - M_{\text{out}}$ remains inactive. As both the input states and the synaptic states are binary, it is likely that several output units have the same dendritic sum. In particular, this happens if $M_{\text{in}}$ and $\eta$ are small because they limit the range of different values that the dendritic sums can take. Fig S17A shows an exemplary histogram of dendritic sums, where the activation threshold $T_{\text{in}}$ cannot be chosen such that exactly the desired number of active output units $M_{\text{out}} = 100$ is activated. We call the dendritic sums $d$ and we define the error

$$M_{\text{err}} := \min_{T_{\text{in}}} \left| M_{\text{out}} - \sum_{\{d \geq T_{\text{in}}\}} 1 \right|, \tag{53}$$

which is the smallest deviation from the exact $M_{\text{out}}$ that can be achieved with the best choice of activation threshold $T_{\text{in}}$. In Fig S17B, we see that the mean error (normalized by $M_{\text{out}}$) decreases with increasing $M_{\text{in}}$ and increasing $\eta$.

In this section, we first provide a theoretical upper bound of the error $M_{\text{err}}$ and compare this estimate to numerical averages. Then, we discuss possible practical ways of dealing with the issue of achieving a given output sparseness in numerical simulations.

Note that these considerations become unnecessary if we approximate the discrete distributions of dendritic sums by continuous normal distributions as in the analytical parts of the Methods Section of this paper.

#### Theoretical analysis

In the following, we derive an upper bound for the error $M_{\text{err}}$ in activating the right number of output units. We first discuss the case $P = 0$, i.e., retrieval immediately after learning a particular pattern, without additional patterns learned in between. The maximal error $M_{\text{err}}$ depends on how many additional units are activated (or deactivated) if the threshold is shifted to the left (or to the right) by 1. The smaller the values of the PMF, the smaller the error we potentially have to make. First, assuming $M_{\text{in}}(1 - \rho_g(0)) \gg 1$ and $M_{\text{in}} c \gg 1$, we approximate the distributions of the dendritic sums of genuine and spurious output units by normal distributions

$$\mathcal{N}\left(\mu_g^{[0]}, {\sigma_g^{[0]}}^2\right) = \mathcal{N}\left(M_{\text{in}}(c + (c_m - c)\eta), M_{\text{in}}(c + (c_m - c)\eta)(1 - c - (c_m - c)\eta)\right) \tag{54}$$

and

$$\mathcal{N}\left(\mu_s, \sigma_s^2\right) = \mathcal{N}\left(M_{\text{in}} c, M_{\text{in}} c(1 - c)\right), \tag{55}$$

respectively. We first consider the genuine distribution and derive an upper bound for the error by focusing on the largest value of the probability density function (PDF) which is at $x = \mu_g^{[0]}$:

$$\mathcal{P}(d_g = \mu_g^{[0]}) \approx \mathcal{N}\left(\mu_g^{[0]}, {\sigma_g^{[0]}}^2\right)(\mu_g^{[0]}) \tag{56}$$

$$= \frac{1}{\sigma_g^{[0]}\sqrt{2\pi}} \exp\left(-\frac{1}{2}\left(\frac{\mu_g^{[0]} - \mu_g^{[0]}}{\sigma_g^{[0]}}\right)^2\right) = \frac{1}{\sigma_g^{[0]}\sqrt{2\pi}}. \tag{57}$$

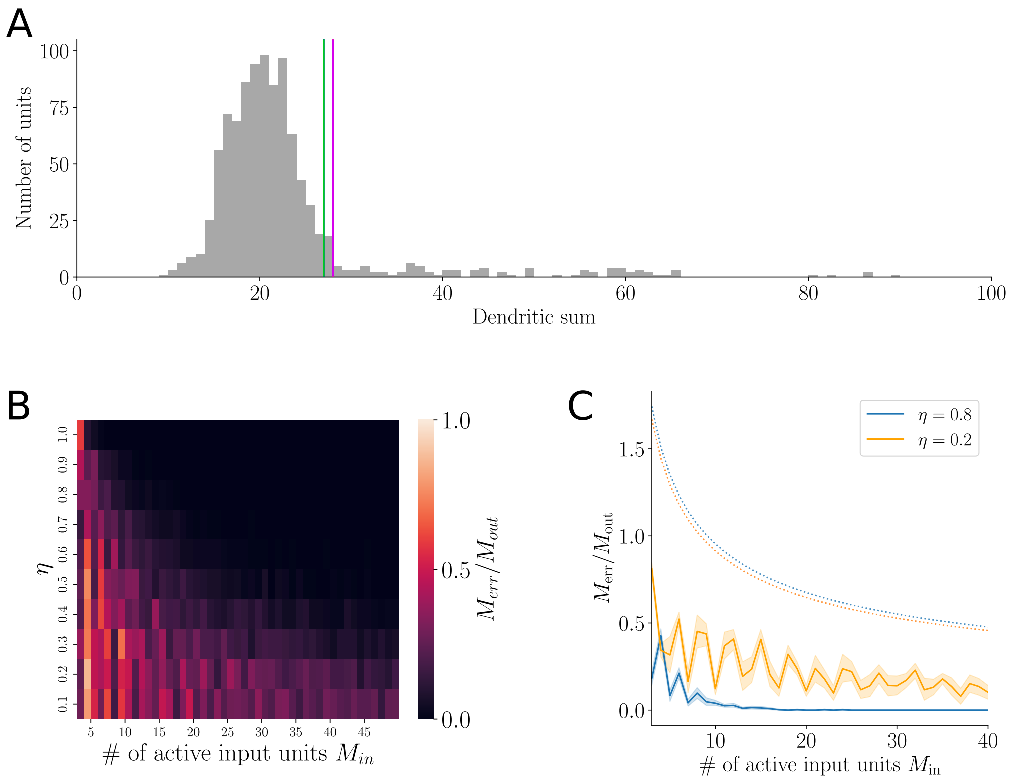


**Fig. S17. Error in number of active output units.** (**A**) In this histogram of dendritic sums (genuine and spurious together), there is no activation threshold $T_{\text{in}}$ that could activate exactly $M_{\text{out}} = 100$ units. If the magenta threshold ($T_{\text{in}} = 28$) is chosen, only 97 units are activated. If the threshold is reduced by one and chosen at the green line ($T_{\text{in}} = 27$), 119 units are activated. In (**A**), $P = 25, \eta = 0.8, N_{\text{in}} = N_{\text{out}} = 1000, f_{\text{in}} = f_{\text{out}} = 0.1, c = 0.2, c_m = 1$. (**B**) The normalized error $M_{\text{err}}/M_{\text{out}}$ decreases as a function of $M_{\text{in}}$ and as a function of $\eta$. (**C**) Comparison between theoretical upper bound of the error (dotted lines, Eq. (69)) and numerically calculated error (solid lines) for $\eta = 0.2$ (orange) and $\eta = 0.8$ (blue). In (**B**) - (**C**), $P = 0, N_{\text{in}} = N_{\text{out}} = 1000, M_{\text{out}} = 48, c = 0.2, c_m = 1$.

The number of genuine units is $M_{\text{out}}$, thus this value needs to be scaled by $M_{\text{out}}$ to describe the average number of genuine units that have a dendritic sum of $\mu_g^{[0]}$:

$$\frac{M_{\text{out}}}{\sqrt{M_{\text{in}}(c + (c_m - c)\eta)(1 - c - (c_m - c)\eta)}\sqrt{2\pi}}. \tag{58}$$

For the spurious distribution, we do the same and obtain the largest value of the PDF

$$\mathcal{P}(d_s = \mu_s) \approx \frac{1}{\sigma_s\sqrt{2\pi}}, \tag{59}$$

which has to be scaled by $N_{\text{out}} - M_{\text{out}}$:

$$\frac{N_{\text{out}} - M_{\text{out}}}{\sqrt{M_{\text{in}}c(1 - c)}\sqrt{2\pi}}. \tag{60}$$

The total change in number of activated units if the threshold is shifted by 1 is the sum of Eqs. (58) and (60). At worst, $M_{\text{err}}$ is half of this sum because if $M_{\text{err}}$ was larger, the activation threshold would not have been chosen optimally:

$$M_{\text{err}} \leq \frac{1}{2}\left(\frac{M_{\text{out}}}{\sqrt{M_{\text{in}}(c+(c_m-c)\eta)(1-c-(c_m-c)\eta)}\sqrt{2\pi}} + \frac{N_{\text{out}}-M_{\text{out}}}{\sqrt{M_{\text{in}}c(1-c)}\sqrt{2\pi}}\right). \tag{61}$$

For $M_{\text{out}} \ll N_{\text{out}}$, this upper bound can be approximated by

$$M_{\text{err}} \leq \frac{1}{2}\left(\frac{N_{\text{out}}}{\sqrt{M_{\text{in}}c(1-c)}\sqrt{2\pi}}\right). \tag{62}$$

It turns out that this simple approximation, which does not depend on $\eta$ often is appropriate even independent of the condition $M_{\text{out}} \ll N_{\text{out}}$: For small values of $\eta$ and a small enough functional connectivity $c$, we have

$$\frac{1}{\sqrt{M_{\text{in}}(c+(c_m-c)\eta)(1-c-(c_m-c)\eta)}} \leq \frac{1}{\sqrt{M_{\text{in}}c(1-c)}}, \tag{63}$$

which is sharper for $c_m$ small and thus close to $c$ but even for $c_m = 1$ and e.g. $\eta \leq 0.5$ it is true for any $c < \frac{1}{3}$.

For $0 < P \ll P^*$, the approximation of the distribution of the dendritic sums of genuine units by a normal distribution can be less accurate (in particular for large $\eta$) and the above analysis cannot easily be generalized. Nevertheless, we know that with increasing $P$ the distribution first becomes wider ($\sigma_g$ is increasing) while the distribution of the dendritic sums of the spurious units does not change with $P$, which decreases the contribution of the genuine units (first term in Eq. (61)) to the change in the number of activated units due to small shifts of the activation threshold. In this case, Eq. (62) is thus also a suitable approximate upper bound. After becoming wider, the distribution of the dendritic sums of the genuine units approaches the distribution of the dendritic sums of the spurious units, which means $1/\sqrt{M_{\text{in}}(c+(c_m-c)\eta)(1-c-(c_m-c)\eta)} \approx 1/\sqrt{M_{\text{in}}c(1-c)}$. For large $P$, we can thus again make use of the same upper bound (62).

For $P = 0$ and $\eta$ close to one, the inequality (63) that we used to simplify condition (61) to (62) is (especially for large $c_m$) usually not fulfilled. (In the limit of $\eta = 1$ and $c_m = 1$, we even have $\sigma_g = 0$ because $\mathcal{P}(d_g = M_{\text{in}}) = 1$.) However, in these cases, the distributions of dendritic sums of genuine units and spurious units are very far apart from each other and it is thus easy to set an activation threshold that will reliably activate the genuine units and deactivate the spurious units anyways. It is very unlikely that a small shift of the threshold will lead to an observable change in the number of activated units.

Although the above upper bound (Eq. (62)) is useful because it is a very simple expression, it relies on rather coarse approximations. In particular, the approximation for the spurious distribution in Eq. (59) can only be reached if $f_{\text{out}} = 0.5$ and a very large number of patterns $P$ such that the genuine distribution is essentially the same as the spurious distribution already. We can more closely approximate the largest relevant value of the spurious probability function $p_s$ by $\mathcal{P}(d_s = x_{f_{\text{out}}})$ with $F_s(x_{f_{\text{out}}}) = 1 - f_{\text{out}}$, where $F_s$ is the cumulative distribution function of the spurious distribution. We find

$$F_s(x_{f_{\text{out}}}) = 1 - f_{\text{out}} \tag{64}$$

$$\Leftrightarrow \frac{1}{2}\left(1 + \operatorname{erf}\left(\frac{x_{f_{\text{out}}} - \mu_s}{\sqrt{2}\sigma_s}\right)\right) = 1 - f_{\text{out}} \tag{65}$$

$$\Leftrightarrow x_{f_{\text{out}}} = \mu_s + \sqrt{2}\sigma_s \operatorname{erf}^{-1}(1 - 2f_{\text{out}}) \tag{66}$$

and hence

$$\mathcal{P}(d_s = x_{f_{\text{out}}}) = \frac{1}{\sigma_s\sqrt{2\pi}} \exp\left(-\frac{1}{2}\left(\frac{\mu_s + \sqrt{2}\sigma_s \operatorname{erf}^{-1}(1-2f_{\text{out}}) - \mu_s}{\sigma_s}\right)^2\right) \tag{67}$$

$$= \frac{1}{\sigma_s\sqrt{2\pi}\cdot\exp\left(\left(\operatorname{erf}^{-1}(1-2f_{\text{out}})\right)^2\right)}. \tag{68}$$

Together with the largest value of the genuine PDF (Eq (57)), this gives the closer upper bound

$$M_{\text{err}} \leq \frac{1}{2}\left(\frac{M_{\text{out}}}{\sigma_g^{[P]}\sqrt{2\pi}} + \frac{N_{\text{out}} - M_{\text{out}}}{\sigma_s\sqrt{2\pi}\cdot\exp\left(\left(\operatorname{erf}^{-1}(1-2f_{\text{out}})\right)^2\right)}\right). \tag{69}$$

Fig S17C compares the analytical upper bound of $M_{\text{err}}$ (Eq. (69), dotted lines) to the actual error in activation ratio obtained in numerical simulations (solid lines). The numerical $M_{\text{err}}$ was calculated by initializing a random weight matrix, presenting the network with a random input pattern with $M_{\text{in}}$ active input units, and measuring the discretization error (Eq. (53)). It is observed that the normalized mean error (as well as its theoretical upper bound) decreases as a function of $M_{\text{in}}$. This is because higher $M_{\text{in}}$ cause spurious and genuine distributions to be further apart, such that the number of units with dendritic sums close to the threshold $T_{\text{in}}$ are lower.

#### Practical solutions

There are several approaches to handle the fact that it is often not possible to activate exactly $M_{\text{out}}$ output units by choosing a fixed activation threshold $T_{\text{in}}$.

**Random sampling of activated units.** One way of achieving the exact given output sparseness is by randomly sampling some of the units that are activated. This method is used in all simulations of this paper unless stated otherwise (see Methods).

If the exact $M_{\text{out}}$ cannot be achieved, this is because any possible activation threshold $T_{\text{in}}$ yields too few or too many activated units. If the activation threshold was chosen as $a$, too few units would be activated, and if it was chosen as $a-1$, too many units would be activated. From all the output units that have the exact same dendritic sum (between $a-1$ and $a$), we randomly choose as many units as are needed to obtain $M_{\text{out}}$ active units. Their activity is set to one while the activity of the rest of the units (with the same dendritic sum) is set to zero. This random choice constitutes an additional (but minor, see Fig S18) source of noise in the simulations.

**Fixed noise mask on weight matrix.** A fixed noise mask on the weight matrix represents an alternative solution. The choice of the activation threshold is ambiguous only because the input activities and the weight values are all zero or one. This yields relatively few distinct dendritic sums. If the dendritic sums were distributed continuously or with a finer discretization, the desired number of active output units $M_{\text{out}}$ could be achieved more easily. The latter can, for example, be realized by point-wise multiplication of the weight matrix $J$ with a fixed noise mask. The noise mask could be defined such that each entry is sampled from a normal distribution with mean $\mu = 1$ and a standard deviation $\sigma \ll 1$. Since weight values now have a small variability, it is much less probable that several output units receive the same dendritic sum and the activation threshold can be easily placed such that exactly $M_{\text{out}}$ units are activated.

Fig S18 shows a comparison of average signal quality traces for the random sampling of activated units and a fixed noise mask on the weight matrix. When averaged across many patterns, there is hardly any difference between the two methods.

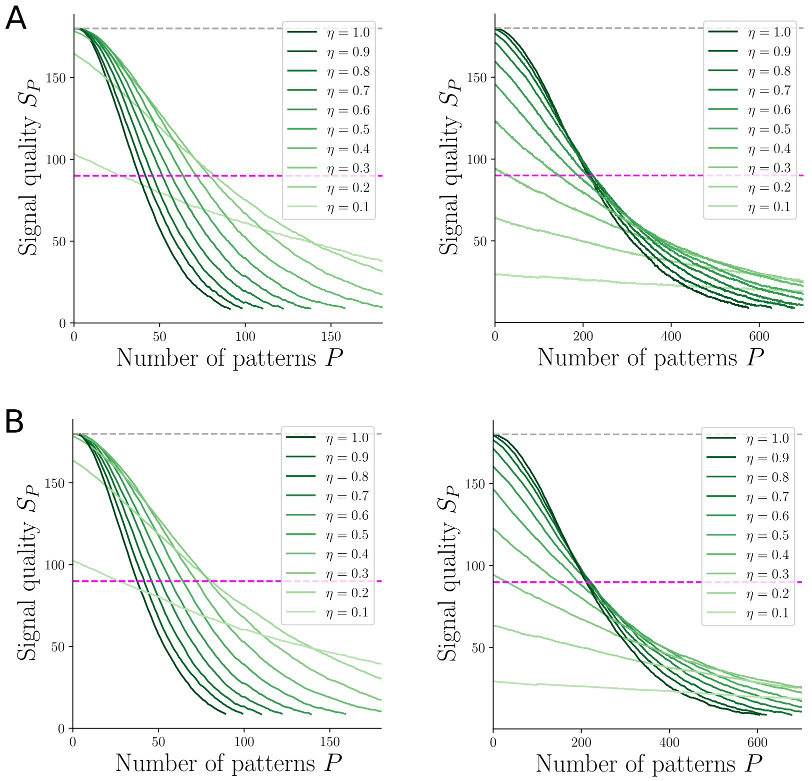


**Fig. S18. Random sampling of activated units or noise mask.** Comparison of signal quality for random sampling of activated units (**A**) and a fixed noise mask on the weight matrix (**B**) for two input activation ratios $f_{\text{in}} = 0.1$ (left) and $f_{\text{in}} = 0.012$ (right). Further parameters:
$N_{\text{in}} = N_{\text{out}} = 1000, f_{\text{out}} = 0.1, c = 0.2, c_m = 1, t_S = 0.5, N_{\text{avg}} = 200$.

**A less strict $f_{\text{out}}$.** Another obvious solution would be to implement a less strict output activation ratio $f_{\text{out}}$. Instead of enforcing an exact number of $M_{\text{out}}$ active output units, there could be a certain tolerance of deviation from this value, e.g., $M_{\text{out}} \pm M_\varepsilon$ with a small $M_\varepsilon \in \mathbb{N}$. In principle, this is a reasonable solution but, since the work in this manuscript is focused on analyzing the impact of sparseness on the optimal transition probability, we want to enforce a fixed $M_{\text{out}}$ and avoid such variabilities in activation ratios.
